# Weak and equally balanced synaptic inputs to interneurons in the CA1 hippocampus characterize *in vivo* rhythmic states

**DOI:** 10.1101/286369

**Authors:** Alexandre Guet-McCreight, Frances K. Skinner

## Abstract

Brain coding strategies are enabled by the balance of synaptic inputs that individual neurons receive as determined by the networks in which they reside. Inhibitory cell types contribute to brain function in distinct ways but recording from specific, inhibitory cell types during behaviour to determine their contributions is difficult. In particular, the *in vivo* activities of vasoactive intestinal peptide-expressing interneuron specific 3 (IS3) cells in the hippocampus that only target other inhibitory cells are unknown at present. We perform a massive, computational exploration of possible synaptic inputs to IS3 cells using multi-compartment models and optimized synaptic parameters. We find that asynchronous *in vivo*-like states that are sensitive to additional theta-timed inputs exist when excitatory and inhibitory synaptic conductances are equally balanced and there are low amounts of correlated inputs. Thus, using a generally applicable computational approach we predict the existence of balanced states in hippocampal circuits during rhythmic activities.

## Introduction

A dizzying array of morphological, molecular, and electrophysiological details for different cell types exist and appropriate classifications are being determined (***Zeng and Sanes, 2017***). How these different cell types contribute to brain function is challenging to determine, but it is clear that a homeostatic balance of cell excitability, together with excitatory and inhibitory synaptic inputs is essential for normal brain function (***Zhou and Yu, 2018; Denève and Machens, 2016; Chiu et al., 2018; Turrigiano, 2011***). The irregular firing of neurons *in vivo* is well-known and is believed to confer computational benefits, with inhibition being recognized as a crucial shaper of these asynchronous activities (***Isaacson and Scanziani, 2011; Treviño, 2016***). Recently, in directly fitting a deterministic firing network model to several sets of *in vivo* multi-neuron data, it was found that the intrinsically generated variability obtained in experiment was mainly due to feedback inhibition (***Stringer et al., 2016***). In essence, it is critical to understand these inhibitory components. However, we are cognisant of the much more diverse nature of inhibitory cells relative to excitatory cells in our brains, despite their smaller overall numbers (***Kepecs and Fishell, 2014; Markram et al., 2004; McBain and Fisahn, 2001***). While the examination of individual neuron activities in the behaving animal is becoming less uncommon, there are certainly more caveats and technical difficulties relative to *in vitro* studies. Further, the smaller numbers and sizes of inhibitory cells as well as being in hard to access locations create additional challenges for identification and patching. Indeed, the activity of several inhibitory cell types *in vivo* remains unknown.

One such cell type that suffers from these difficulties are hippocampal CA1 interneuron specific type 3 (IS3) interneurons. IS3 cells are a vasoactive intestinal polypeptide-positive (VIP+) and calretinin-positive (CR+) cell type with cell bodies found in the stratum radiatum and stratum pyramidale of the CA1 (***Acsády et al., 1996a***,b; ***Gulyás et al., 1996; Chamberland and Topolnik, 2012***), an area in CA1 more predominantly populated by pyramidal cells as well as some parvalbumin-positive (PV+) cell types. Compared to pyramidal cells, which make up approximately 80-90% of neurons in CA1 (***Pelkey et al., 2017; Vida, 2010***), IS3 cells make up less than one percent of the CA1 neurons (***Bezaire and Soltesz, 2013***), making them much more difficult to locate. Previous work has circumvented these issues through usage of GFP-VIP mouse lines such that IS3 cells can be readily identified *in vitro* (***Tyan et al., 2014***). Further, new calcium imaging techniques show promise in recording network activity of IS3 cells *in vivo* (***Villette et al., 2017***), but electrophysiological recordings during *in vivo* hippocampal rhythms such as theta has not been characterized. *In vivo* patch-clamp recordings have been obtained from other hippocampal inhibitory cell types such as oriens-lacunosum/moleculare (OLM) cells, ivy cells, bistratified cells, axo-axonic cells, and basket cells (***Klausberger et al., 2003; Lapray et al., 2012; Katona et al., 2014; Varga et al., 2012, 2014***), yet, due to the difficult nature of these techniques, these experiments often suffer from low numbers of recordings. Although IS3 cells only represent a small percentage of CA1 network neurons, understanding their contributions to network activity is a compelling exploration in hippocampal research due to their distinct connections exclusively onto other inhibitory interneurons, such as OLM and bistratified cells (***Tyan et al., 2014; Chamberland et al., 2010***). This unique circuitry suggests a potential function in the disinhibition of pyramidal cells. Such a contribution to network function has in fact been observed in VIP+ cells of different cortical areas during a variety of different behavioral contexts (***Pi et al., 2013; Lee et al., 2013; Fu et al., 2014; Zhang et al., 2014; Karnani et al., 2016a***,b; ***Jackson et al., 2016; Kamigaki and Dan, 2017***).

Despite the challenges of *in vivo* recordings, much can be said about the *in vivo* state. That is, *in vivo* recordings are irregular and asynchronous and constitute “high-conductance” states (***Destexhe et al., 2003***), where a high level of synaptic bombardment causes the cells’ subthreshold membrane potential to be more depolarized, to have larger fluctuations, and to have smaller input resistances (***Destexhe and Paré, 1999***). To move toward an understanding of IS3 cell contributions to brain function, we use a computational approach by necessity, as recording from these cell types *in vivo* is not possible at the present time. We explore what excitatory and inhibitory inputs could bring about asynchronous states in IS3 cells and also be sensitive to theta frequency inputs as has been shown *in vitro* (***Tyan et al., 2014***). We find that excitatory and inhibitory synaptic currents need to be equally balanced with correlated inputs (common inputs) and small numbers of excitatory and inhibitory synapses and low presynaptic input firing rates. This work contributes to an understanding of excitatory and inhibitory balances in hippocampal circuits.

## Results

Due to the diversity and details of inhibitory cells it is challenging to understand their roles and contributions in brain circuits and behaviours. We use a computational approach to simulate *in vivo* activities so as to determine how much and what balance of excitatory and inhibitory synaptic inputs inhibitory cell subtypes might receive in the behaving animal. We illustrate this in the ***Figure 1*** schematic for IS3 cells that we examine here.

**Figure 1.**
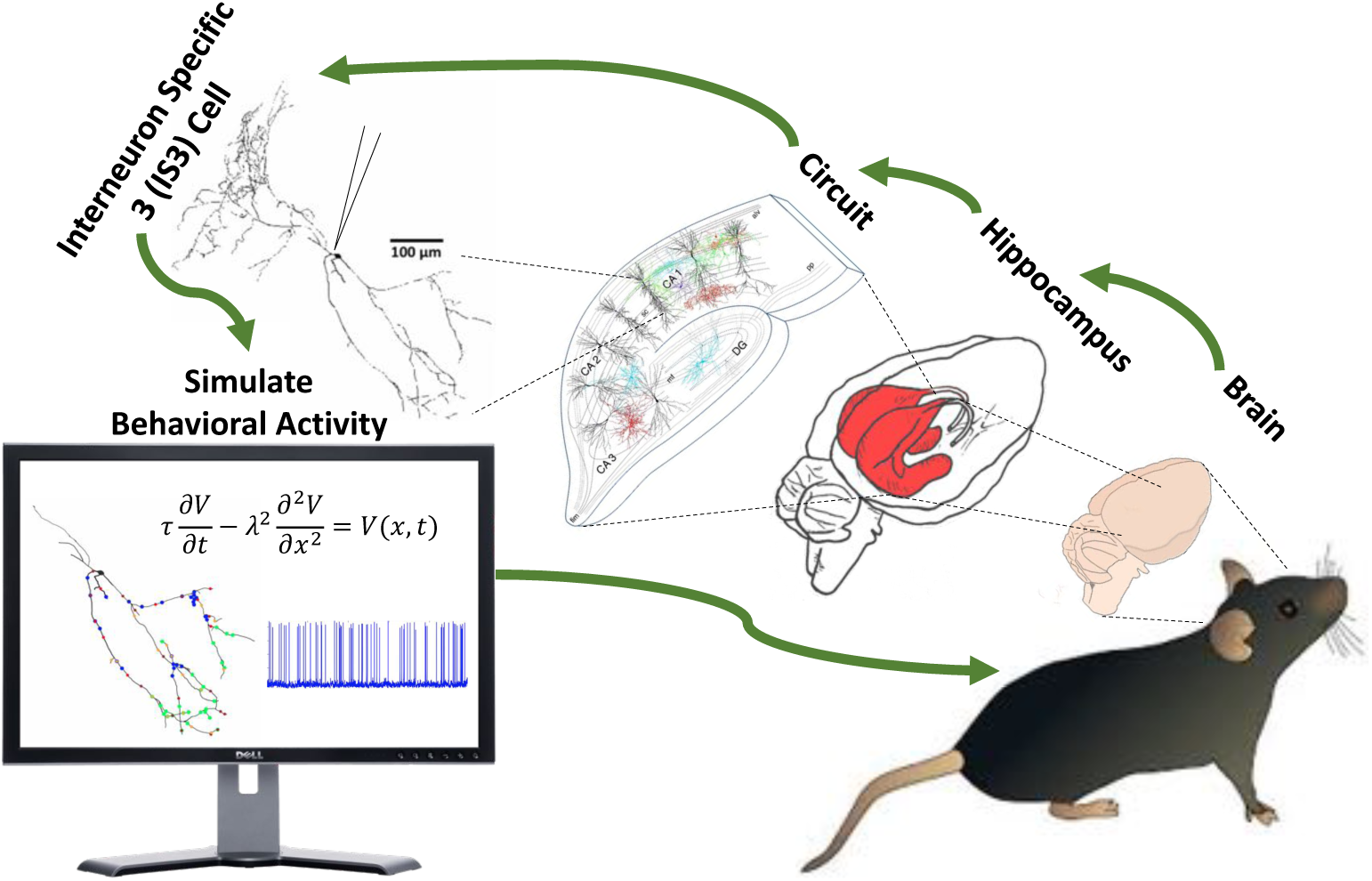
Schematic of computational approach. In behaving animals, it is difficult to perform patch-clamp experiments on cells in subcortical **brain** structures such as the **hippocampus**. Because **IS3 cells** are a small percentage of neurons in the CA1 **circuit**, and their morphologies are small relative to other cell types it is especially difficult to record from them. We use mathematical modeling to **simulate** *in vivo* activities so that predictions of circumstances under which these cell types are recruited *in vivo* can be made. The circuit image in this figure was taken from (***Ouyang et al., 2017***).

We use our previously developed multi-compartment models of IS3 cells in which the electro-physiological features of IS3 cells were captured (***Guet-McCreight et al., 2016***). Our models are detailed in terms of morphology and the inclusion of four types of ion channels, and we use two variant IS3 cell models. One model, SDprox1, has dendritic A-type potassium channels, and the other model, SDprox2, does not. Further model details and parameter values are given in the Methods section. To perform parameter explorations of input regimes that yield *in vivo*-like states, we first considered the types of layer-specific excitatory and inhibitory inputs to IS3 cells. Within the hippocampus, IS3 cells have dendrites that extend into the stratum radiatum and stratum pyramidale, which allows them to receive excitatory inputs from both CA3 and entorhinal cortex. Layer-specific inhibitory inputs to IS3 cells are not characterized, though there are several possible inhibitory presynaptic populations to IS3 cells. For this reason we consider one generic population of inhibitory inputs, one population of proximal excitatory inputs, and one population of distal excitatory inputs. This is illustrated in the schematic of ***Figure 1, Figure Supplement 1***. Using our two IS3 multi-compartment models, we investigated a wide range of parameter values characterizing excitatory and inhibitory synaptic inputs onto these cell models. We explored the full range of possible input parameter combinations based on what is known experimentally to obtain *in vivo*-like states. This is illustrated in the schematic of ***Figure 1, Figure Supplement 2*** and details of our parameter exploration strategy and setup is given in the Methods. Based on what is known, we devised an HC metric (***Equation 5***) to identify *in vivo*-like or high-conductance (HC) states. As detailed in the Methods, output from our IS3 cell models is considered to appropriately represent an *in vivo*-like state if our HC metric has a value of three. A negative or zero HC metric indicates that the output is, respectively, nearing depolarization block (DB) or in a non-high-conductance (NHC) state. Consideration of these additional states will help us better interpret our results.

### Common inputs and an inhibitory dominance promote *in vivo*-like states

We display our results from parameter explorations using clutter-based dimensional reordering (CBDR) plots (***Taylor et al., 2006***). This enables us to plot multi-dimensional parameter sets as two-dimensional plots. ***Figure 2*** illustrates this for a parameter exploration using the SDprox1 model with common excitatory and inhibitory inputs. Note that white pixels represent HC or *in vivo*-like states, orange pixels represent NHC states and dark pixels (i.e. red to black) represent scenarios nearing DB. Voltage output for a particular set of parameters when it is in a HC state (i.e., HC metric = 3) is also shown in ***Figure 2***.

**Figure 2.**
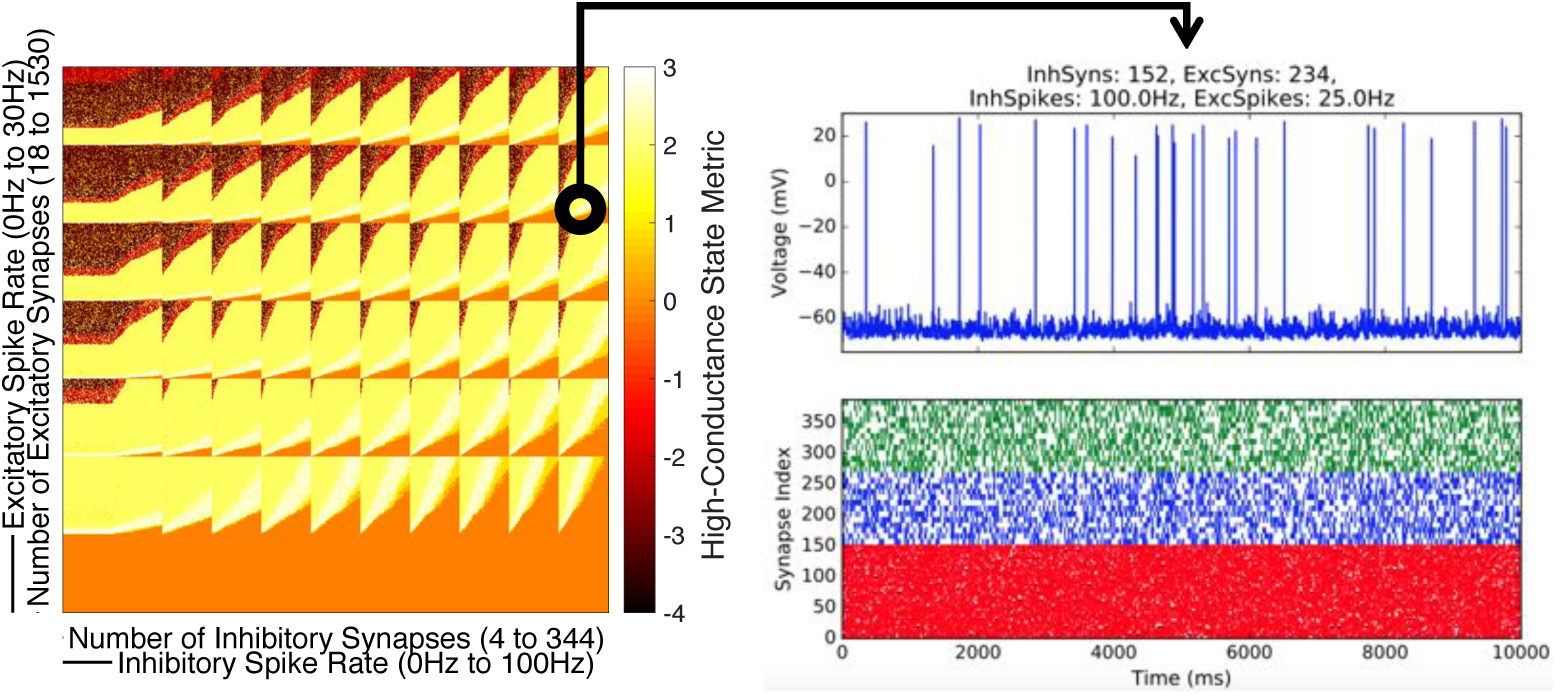
Display of Results. **(Left):** A clutter-based dimensional reordering (CBDR) plot of a parameter exploration for the SDprox1 model with common excitatory and inhibitory inputs. Excitatory input parameters are indicated by the scale bars on the y-axis and inhibitory input parameters are indicated by the scale bars on the x-axis, with parameter ranges shown in parentheses. Each pixel represents a 10 second simulation where the color of the pixel indicates the high-conductance (HC) metric score for the particular set of parameters. Using the scale bars, one can extrapolate the precise parameters of each individual pixel. Note that the height and width of the pixels are of equal size to the lengths of the smaller scale bars on the y- and x-axes. For example, going from bottom to top at an interval of the length of the larger scale bar, the excitatory spike rate increases in increments of 5 Hz. Likewise, going from bottom to top at an interval of the length of the smaller scale bar, the number of excitatory synapses increases in increments of 18 synapses, until it reaches the length of the larger scale bar, at which point it restarts the count. Similarly, the inhibitory spike rate increases in increments of 10 Hz going from left to right, and the number of inhibitory synapses increases in increments of 4 synapses. **(Right):** Output from one of the parameter sets (HC metric=3, white pixel) as indicated by the arrow. The top subplot shows the 10 second simulated voltage trace from the pixel indicated in the Left CBDR plot. Bottom subplot shows the raster plot of the presynaptic inputs where inhibitory inputs are shown in red, proximal excitatory inputs are shown in blue and distal excitatory inputs are shown in green. Note that these plots do not contain any information regarding the synaptic locations of the inputs, which are chosen randomly.

In ***Figure 3*** we show results from our full set of parameter explorations using SDprox1 and SD-prox2 models and with common or independent inputs. From our CBDR plots we can immediately observe that common excitatory inputs promote *in vivo*-like or HC states since there are more white pixels (HC metric=3) in the two right columns of ***Figure 3***. The exact number of scenarios for HC states are given in the caption of ***Figure 3***, and they indicate that having common inputs for both excitatory and inhibitory synapses maximizes the number of HC states.

**Figure 3.**
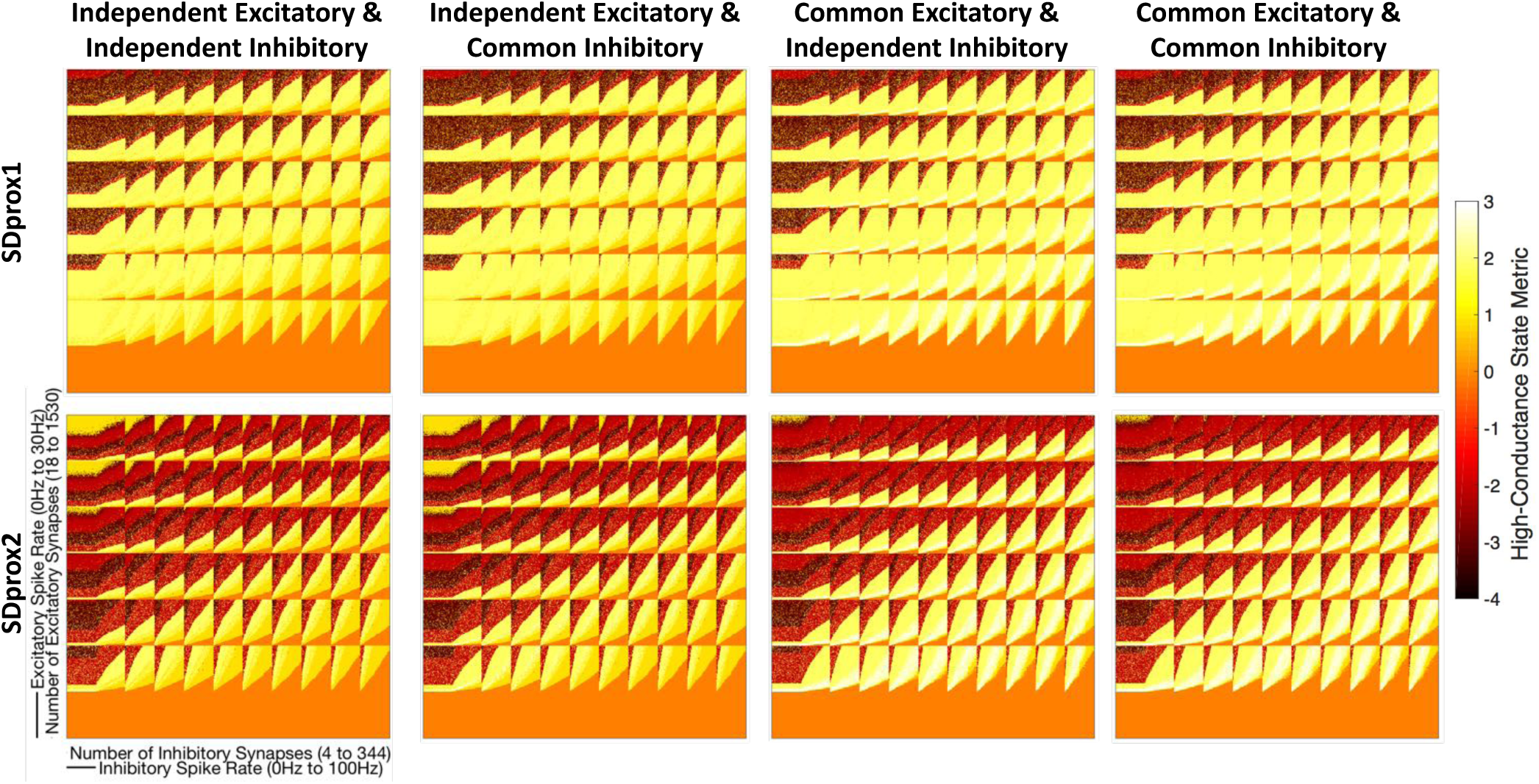
CBDR plots for SDprox1 and SDprox2 models and for all possible combinations of common and independent excitatory and inhibitory inputs. In these plots the high-conductance (HC) metric is plotted, where white pixels (HC metric=3) signify that the model is in an *in vivo*-like state. The number of HC or *in vivo*-like scenarios found for each SDprox1 independent/common condition is (from left to right): 920, 2414, 38785, 39939. Similarly for SDprox2: 1994, 7152, 44642, 46655.

The separate parts of our HC metric (***Equation 5***) can be seen in supplementary figures of ***Figure 3*** (***Figure Supplement 1, Figure Supplement 2, Figure Supplement 3***). Looking specifically at the subthreshold membrane potential standard deviation 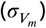 and interspike interval coefficient of variation (*ISICV*) metrics (***Figure Supplement 2, Figure Supplement 3*** respectively), it is clear that the larger number of HC states with common inputs that we observe in ***Figure 3*** is mainly due to these characteristics having a larger parameter space that exceed their chosen thresholds to represent an *in vivo*-like or HC state. This makes sense since common excitatory or inhibitory inputs will result in a single presynaptic spike train causing larger deflections in the IS3 cell model (i.e. larger 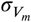). Also, it is more likely that irregularly timed spiking in the IS3 cell model (i.e. larger *ISICV*) would occur since the presynaptic spike times are sampled randomly and are irregular, and excitatory input arriving at multiple synaptic locations simultaneously would increase the likelihood that the IS3 cell model could evoke a spike (***Figure 1, Figure Supplement 2***A).

A clear observation when comparing SDprox1 and SDprox2 models is the much larger number of darker pixels for the SDprox2 model relative to the SDprox1 model which indicates that the SDprox2 model enters depolarization block much more readily than the SDprox1 model. This is likely due to the lack of dendritic A-type potassium in the SDprox2 model making it more excitable. To this point, consider the spike rates of the IS3 cell models as shown in ***Figure 3, Figure Supplement 4***. For larger excitatory spike rates and number of excitatory synapses (i.e., the top left regions of the CBDR plots), the spike rates are closer to zero for the SDprox2 models but not for the SDprox1 models.

Moving forward, we focus only on scenarios in which there are common inputs since it is clear that having common inputs maximizes the space of parameter sets that could generate *in vivo*-like states. However, we retain our consideration of both SDprox1 and SDprox2 models. To get a handle on these different scenarios we examine the balance of excitation and inhibition using two different EI metrics as given by ***Equation 6*** and ***Equation 7*** in the Methods. For both metrics, a negative value implies a dominance of inhibitory inputs. A metric value of zero implies that excitation and inhibition are approximately equal. We note that since synaptic inputs are not present at the same location and our metrics do not take spatial aspects into consideration, an EI metric value of zero does not necessarily mean that inputs are perfectly balanced in terms of amount and spatial spread. In ***Figure 3, Figure Supplement 5*** we show histogram distributions of our EI metrics for the HC states. From these plots, it is clear that the majority of HC state scenarios exhibit an inhibitory dominance. Histogram distributions for DB and NHC states are shown together with HC states in ***Figure 3, Figure Supplement 6***. Predictably, the DB distributions tend to lean toward excitatory-dominant regimes with low amounts of inhibition as represented by positive EI metric values. These plots also highlight the larger number of DB scenarios generated when using the SDprox2 model rather than the SDprox1 model potentially due to the lack of dendritic A-type potassium in SDprox2 as noted above. We note that the HC distributions fall within the NHC distributions for both metrics and that the NHC distributions tend to lean much more towards inhibitory dominant-regimes with low amounts of excitation. Further, looking at EI metric #2, we can make two observations. First, in the NHC distributions there are large peaks at zero (***Figure 3, Figure Supplement 6***) indicating balanced excitation and inhibition, which likely represents a pool of scenarios which have zero Hz spike rates which would not be present in HC scenarios since the requirements for an *in vivo*-like state would not be satisfied. Second, we also notice that the HC distributions for EI metric #2 have peaks just above zero (***Figure 3, Figure Supplement 5***), whereas the NHC distributions do not (***Figure 3, Figure Supplement 6***), suggesting a subpopulation of HC scenarios that require balanced input parameters to satisfy our requirements for an *in vivo*-like state.

### Low amounts of balanced synaptic inputs and high amounts of inhibitory-dominant synaptic inputs can both generate *in vivo*-like states

To determine subspaces of parameter sets as well as uncover relationships between HC states and balances between excitation and inhibition, we divided the parameter space into 16 pools by splitting each parameter into high and low ranges as specified in ***Table 1***. The acronym naming scheme we use is given in ***Table 2*** and we use these acronyms in subsequent figures.

**Table 1.** Ranges for each parameter when splitting the parameter space into 16 pools.

| Parameter | Low | High |
| --- | --- | --- |
| Excitatory Spike Rate (Hz) | 0-15 | 20-30 |
| Inhibitory Spike Rate (Hz) | 0-50 | 60-100 |
| Number of Excitatory Synapses | 18-765 | 783-1530 |
| Number of Inhibitory Synapses | 4-172 | 176-344 |

**Table 2.**
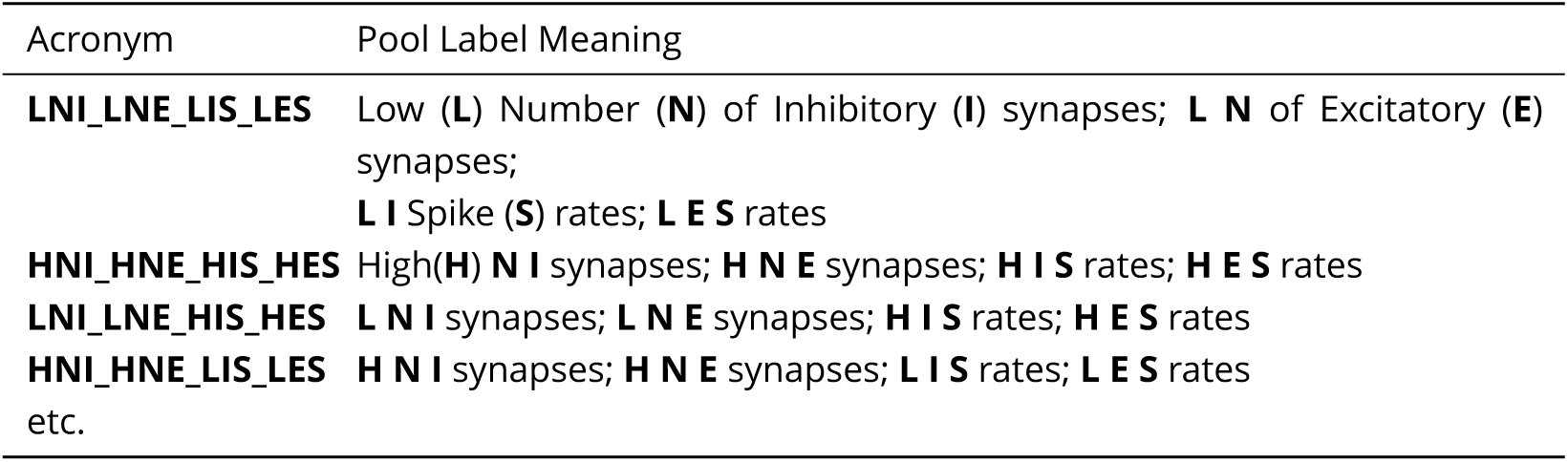
Naming scheme for the 16 different pool labels, see ***Figure 4***.

In ***Figure 4*** we show these 16 pools as red or green squares where red squares represent parameter sets where no HC states were found. For both SDprox1 and SDprox 2, there are three pools where no HC states were found and they encompass parameter sets with high excitatory spike rates and high numbers of excitatory synapses. For the other 13 pools the number of HC scenarios is specified, and for each pool in ***Figure 4*** the pool label (***Table 2***) is specified. The delineation of the parameter values of the 16 pools is shown in ***Figure 4, Figure Supplement 1*** plots. From these plots it is clear that pools with high amounts of inhibition and low amounts of excitation have large numbers of NHC scenarios, and there are large amounts of DB scenarios for the inverse. As noted earlier, scenarios for HC states tend to occur in pools with larger amounts of inhibition. We illustrate this using colors and the 16 pool representation described in ***Figure 4, Figure Supplement 2***. Also shown are the IS3 cell model spike rates where there does not appear to be a large variation across pools. It is interesting to note that having a larger amount of inputs did not necessarily result in higher mean spike rates.

**Figure 4.**
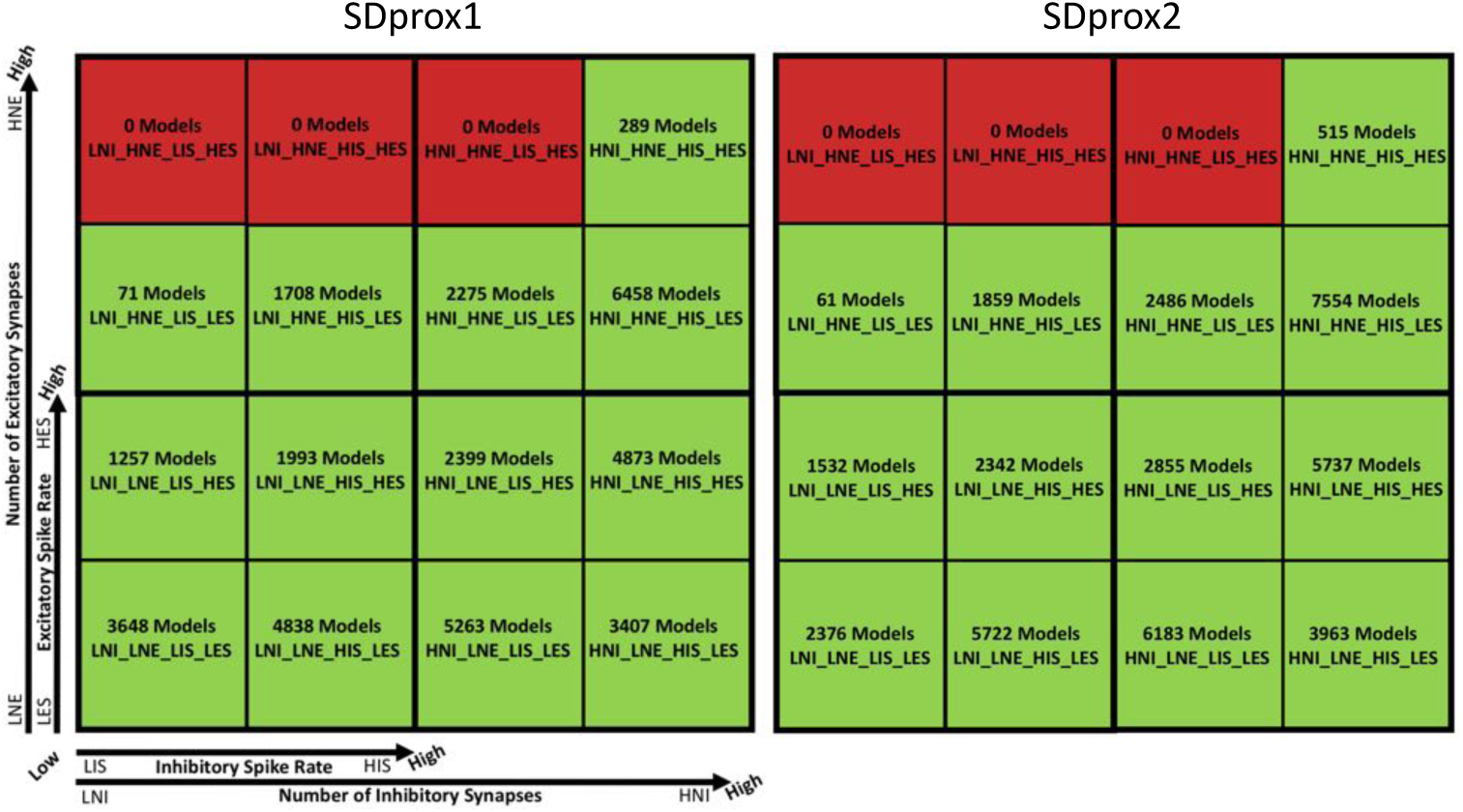
Schematic of parameter space pool divisions. Pools in green generated HC states while pools in red did not generate any. Number of HC scenarios for each pool are indicated in each square as well as the pool labels. As detailed in ***Table 2***, labels L or H represent “low” or “high” parameter ranges, NI or NE represents “number” of “inhibitory” or “excitatory” synapses, and IS or ES represents “inhibitory” or “excitatory” “spike” rates. The smaller axes refer to each quarter of the entire large square. Axis labels are the same for SDprox1 and SDprox2.

We plot our EI metrics for the 16 pools in ***Figure 5***, and ***Figure 5, Figure Supplement 1, Figure Supplement 2*** for HC, NHC and DB scenarios. From close observation, we find that the pool mostly straddling a zero value EI metric is the one with low inputs (LNI_LNE_LIS_LES). We plot this pool on its own in ***Figure 5*** for the two EI metrics and SDprox1 and SDprox2 models. From the supplemental figures of ***Figure 5*** we can once again observe that the SDprox2 model produces a considerably larger number of DB scenarios relative to the SDprox1 model. We further observe that the HC pool distributions are shifted towards more balanced metric values (i.e., EI metric values of zero), when compared to their NHC pool counterparts. Furthermore, the largest NHC pool distribution (brown - HNI_LNE_HIS_LES), is not prominent in the HC pool distributions plots. Likewise, the largest HC pool distribution (grey - HNI_HNE_HIS_LES), is not prominent in the NHC pool distribution plots. This provides evidence that, despite having overlap in inhibitory-dominant regimes (***Figure 3, Figure Supplement 6***), there are differences in input parameter balances that differentiate HC states versus overly inhibited NHC states. Together, these findings suggest that HC states with low amounts of inputs will be more likely to have balanced excitatory and inhibitory inputs, and HC states with large amounts of inputs will tend to have inhibitory-dominant inputs.

**Figure 5.**
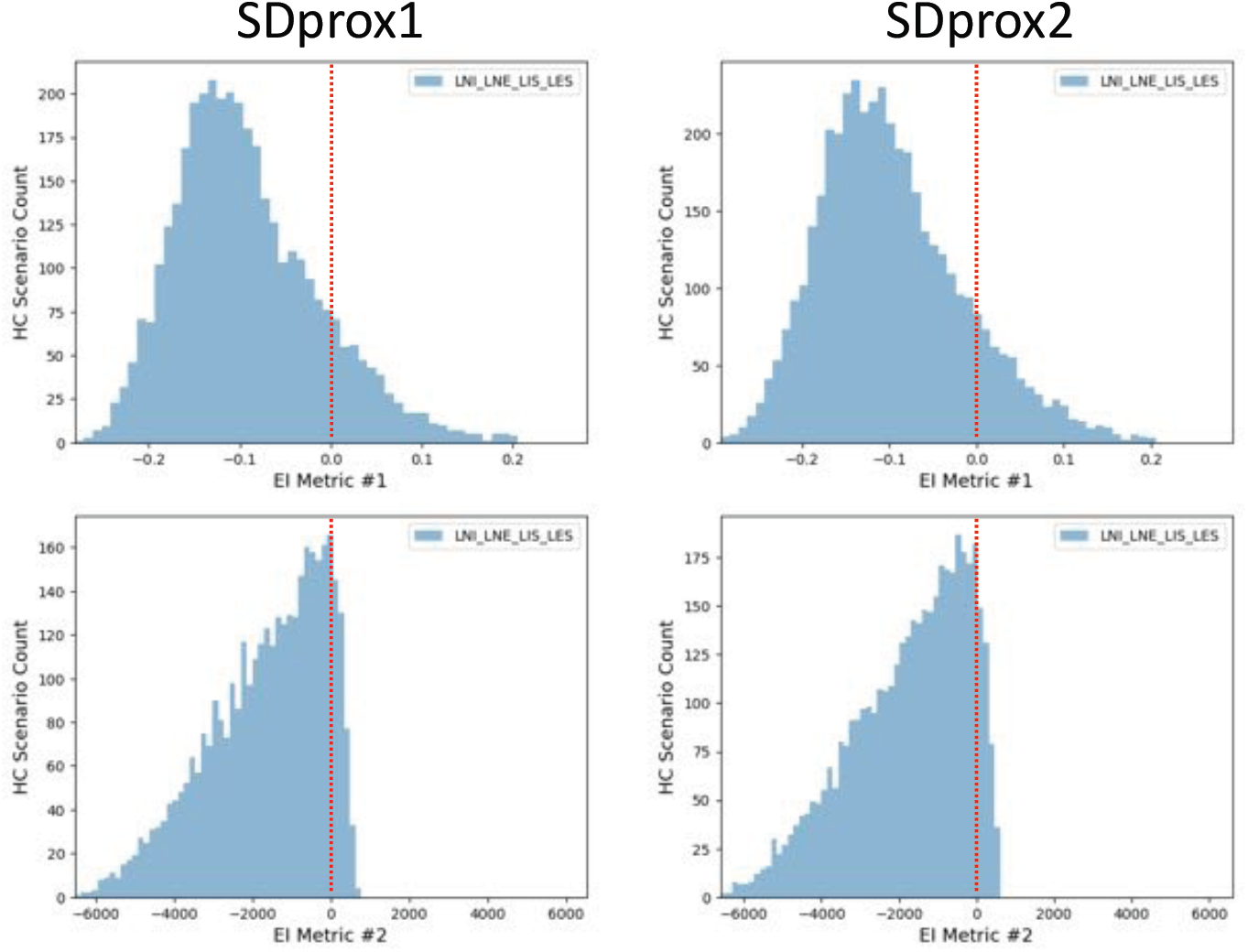
Zoomed in distributions of EI metric values for the LNI_LNE_LIS_LES pool for both SDprox1 and SDprox2 models. Red dashed lines indicate the EI value of zero (for both metrics) where excitation and inhibition are even.

### Moving closer to predicting synaptic inputs to IS3 cells *in vivo*

To determine what sort of inputs IS3 cells might receive *in vivo*, we perform a deeper exploration of our parameter sets. That is, despite already doing millions of simulations (see Methods), we perform additional simulations to probe and understand the robustness of our HC or *in vivo*-like states. We do this by obtaining a representative scenario from each pool that is robust, as determined by using a procedure that cycles through several random seeds. The procedure is described in the Methods, and in ***Figure 6*** we show the voltage output from a representative scenario from each pool that is robustly HC for a given set of random seeds. Also shown in ***Figure 6*** are raster plots of the excitatory and inhibitory synaptic inputs. The synaptic input parameters for each of the representative scenarios is given in ***Figure 6, Figure Supplement 1***, and the synaptic locations for them are shown in ***Figure 6, Figure Supplement 2*** for a given random seed. We already know that 3 of the 16 pools did not produce HC states (red squares), and in searching for robust, representative scenarios, another pool (LNI_HNE_LIS_LES) is removed (yellow square) for each of the SDprox1 and SDprox2 models as a representative scenario was not found. We note that this pool actually had the smallest number of HC states to begin with (see ***Figure 4***). Thus we have now reduced the possible *in vivo*-like states to being from 12 pools and we have chosen a representative scenario from each of these pools. Additional considerations regarding intrinsic cell noise (see ***Figure 6, Figure Supplement 3***) and common inputs (see ***Figure 6*** figure supplements) are described in the Methods.

**Figure 6.**
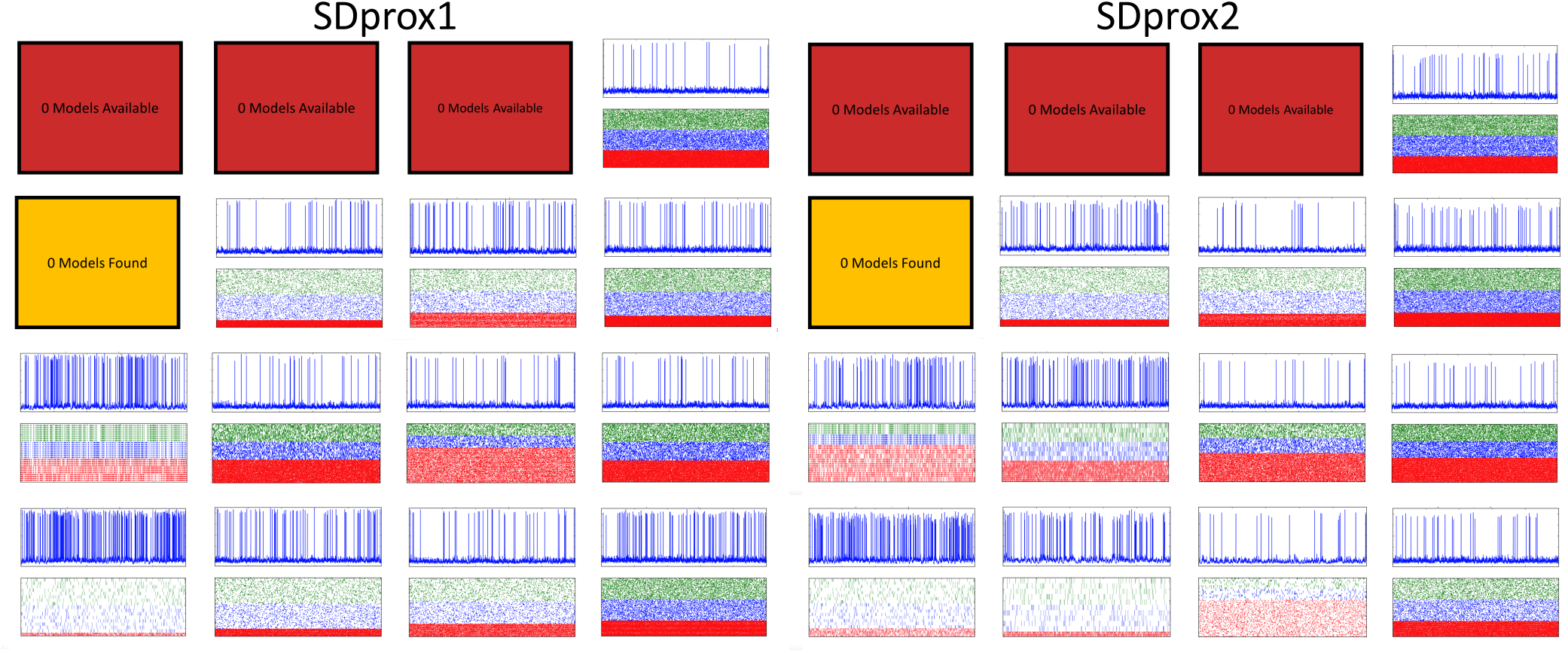
Representative HC scenario from each pool. Voltage traces (top subplot y-axes: voltage from -80 to +30 mV) and input rasters (bottom subplots: see ***Figure 6, Figure Supplement 1*** for synapse index y-axis ranges) for one of the random seeds. Traces and raster plot descriptions are the same as in ***Figure 2*** (x-axis: time from 0 to 10,000 ms).

For each representative scenario we re-do the simulation ten times with different random seeds to examine the consistency of their HC states. In ***Figure 7*** we show that the input resistance and mean spike rates of IS3 cell models vary for the representative scenarios. The input resistances are smaller compared to our model IS3 cells when not receiving synaptic input. The specific values are given in the caption of ***Figure 7***. Previous reports have in fact indicated that there are decreases in input resistance during *in vivo* states (***Destexhe and Paré, 1999; Monier et al., 2008***), which makes sense due to the larger amount of synaptic inputs being received *in vivo* relative to *in vitro*. Looking at the mean spike rates of the IS3 cells, we observe that they are larger when there are low amounts of inputs (***Figure 7***, bottom plots; see LNI_LNE_LIS_LES, and also see ***Figure 6***). This is logical and consistent with our results above that show that there is an inhibitory dominance with high amounts of inputs. In ***Figure 6, Figure Supplement 3*** we dissect the consistency of the HC states of the representative scenarios by plotting the different parts of the HC metric along with the % of simulations that the HC metric remained at 3. We see that the representative scenario from pool LNI_LNE_LIS_LES is consistently HC for both SDprox1 and SDprox2 models for most of the re-done simulations, and when it is not, it is due to the *ISICV* dipping a bit below the chosen threshold. This is also true for the representative HC scenario from pool LNI_LNE_LIS_HES for the SDprox1 model and from pool LNI_LNE_HIS_LES for the SDprox2 model. Interestingly, these four pools are those that have the largest input resistances and mean spike rates (see ***Figure 7***). For the representative scenarios in other pools, changing the random seed had a larger impact on the consistency of the HC state because the *ISICV* fell quite a bit below the chosen threshold of 0.8 on some of the trials, although we note that it has been demonstrated that *ISICV* values lower than our chosen threshold value of 0.8 can occur (***Softky and Koch, 1993***).

**Figure 7.**
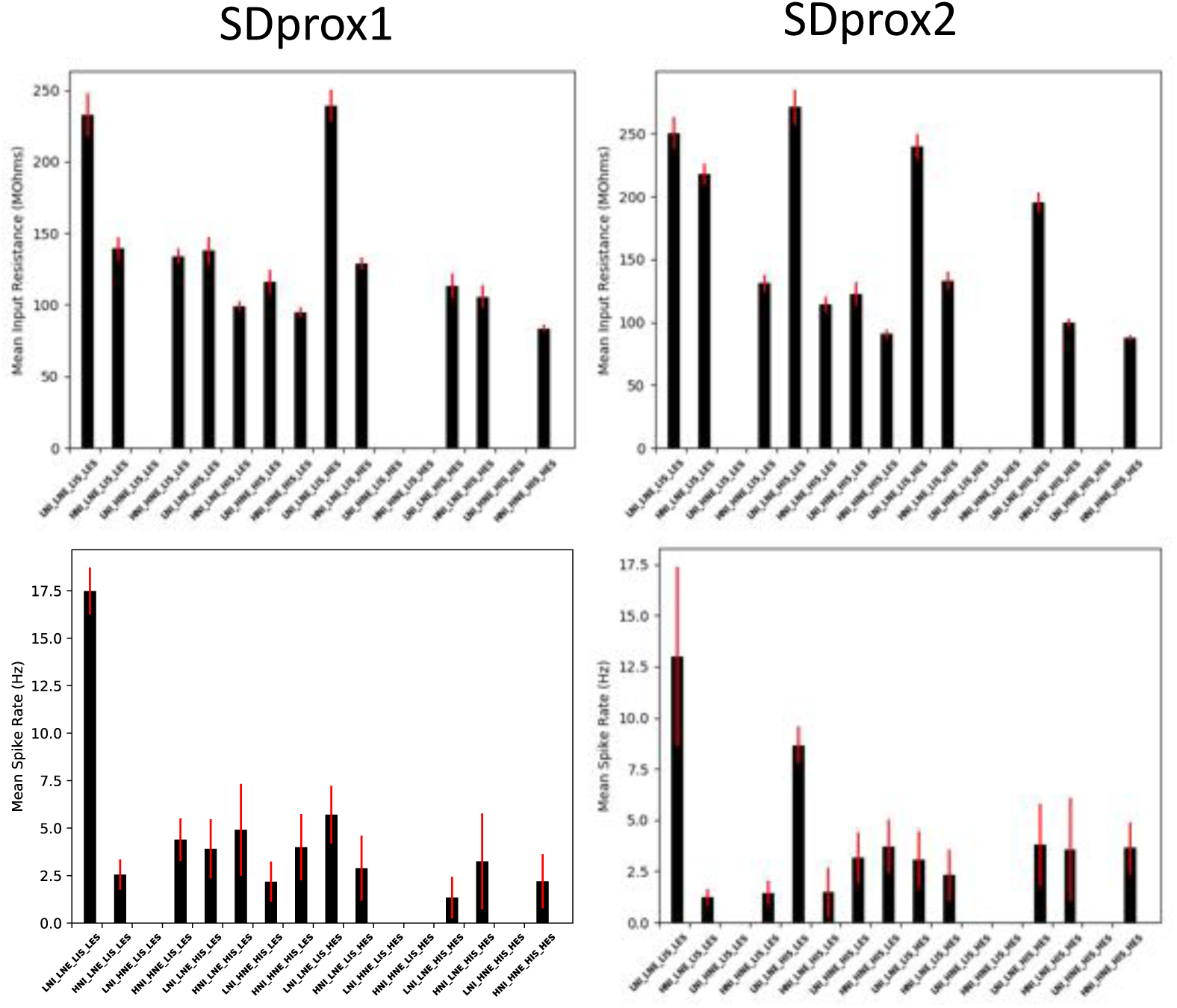
Input resistances and spike rates of IS3 cells in the representative HC scenarios. Top plots: Mean input resistance values, as computed across 10 simulations with different random seed values on each iteration. Standard deviations of the input resistance values are shown as red error bars. Bottom plots: Mean spike rates, as computed across 10 simulations with different random seed values on each iteration. Standard deviations of the spike rates are shown as red error bars. When not receiving any synaptic input, models show no spiking and input resistance is: 388.71 MOhms (SDprox1), 406.57 MOhms (SDprox2).

#### HC scenarios with low amounts of synaptic input have balanced synaptic currents and conductances

Though our EI metrics have shown that HC scenarios with low amounts of synaptic input are more likely to have balanced excitation and inhibition, our EI metrics do not include differences in distance-dependent synaptic parameters. Our inhibitory and excitatory synapses have different distance-dependent kinetics, as determined from optimized fits to experiment (see ***Equation 3*** and ***Equation 4*** in the Methods). Additionally, inhibitory synapses in our models have slower time constants than excitatory synapses. We thus directly examine the balance of excitation and inhibition by isolating excitatory and inhibitory currents (see Methods for details) in our representative HC scenarios from the different pools. In viewing these results, we see that the representative HC scenarios for the pools with lower amounts of synaptic input (***Figure 8***, bottom left panels) exhibit more balanced magnitudes of excitatory and inhibitory currents and conductances than those with larger amounts of inputs, which are all inhibitory-dominant.

**Figure 8.**
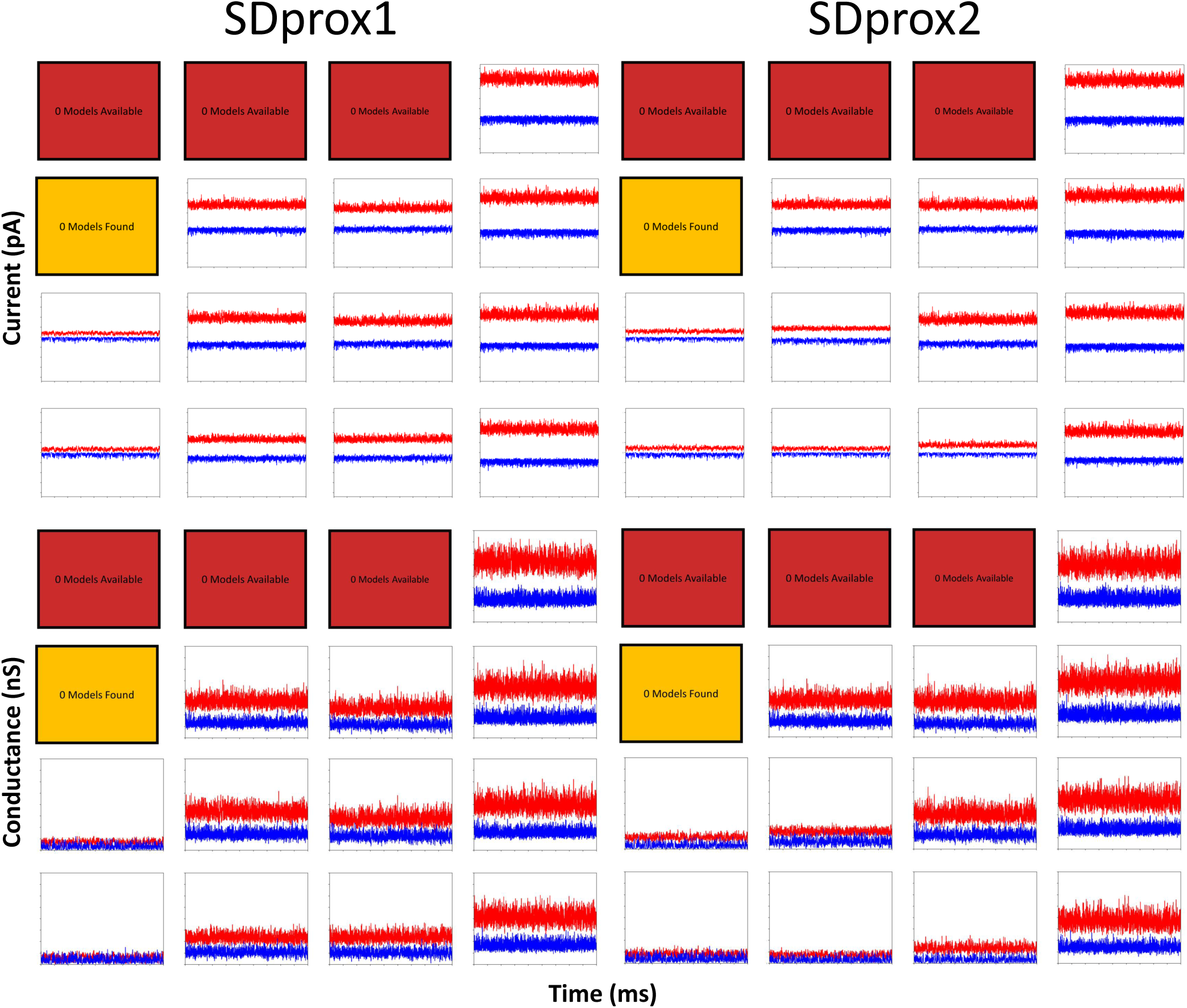
Inhibitory or excitatory currents (y-axes: -1100 pA to +1100 pA) and conductances (y-axes: 0 nS to 16 nS) recorded when excitatory or inhibitory currents are respectively removed completely (x-axes: 1000 ms to 10,000 ms). Top panels: excitatory (blue) and inhibitory (red) currents recorded in representative HC scenarios of the SDprox1 and SDprox2 models. Bottom panels: excitatory (blue) and inhibitory (red) conductances recorded in representative HC scenarios of the SDprox1 and SDprox2 models.

It is interesting to consider potential confounding effects that would be present in experimental situations. That is, to separate out excitatory or inhibitory currents, one would hold the cell at either the inhibitory or excitatory reversal potential respectively. Besides not necessarily knowing the appropriate reversal potential in experiment, one would also have space clamp issues. Of course, we do not have these limitations in models so that we can be sure about the excitatory and inhibitory currents and conductances plotted in ***Figure 8***. In ***Figure 8, Figure Supplement 1*** we show the synaptic currents and conductances that are obtained in our models if we separate excitatory and inhibitory currents as would be done experimentally. There is a clear difference between ***Figure 8*** and ***Figure 8, Figure Supplement 1***. Because the model is only clamped at the soma, inhibitory currents can still have an effect of reducing the current generated by excitatory synapses (i.e. space-clamp issues). This is most apparent when comparing the excitatory currents and conductances since the excitatory conductances remain ‘glued’ to zero using the experimental approach which is a clear artifact. Since we do not have this artifact we can clearly state that *in vivo*-like states in our IS3 cell models when there is a low amount of synaptic input have balanced excitatory and inhibitory inputs, and having a larger amount of synaptic input requires a shift towards an inhibitory dominance to maintain an *in vivo*-like state.

### IS3 cells need to have balanced excitation and inhibition to exhibit sensitivity to theta-timed inputs

The lack of *in vivo* recordings of IS3 cells means that it is unknown how they might contribute to brain function. However, given their connections to other inhibitory cell types that are involved in theta rhythms, and that they can preferentially influence OLM cells at theta frequencies *in vitro* (***Tyan et al., 2014***), it is reasonable to hypothesize that IS3 cells need to be responsive to theta-timed inputs. To examine this, we implemented theta-timed layer-specific inputs at 8 Hz. In order to define these theta-timed input populations, we approximated synapse numbers (see Methods) as well as the relative timing of both known and hypothetical excitatory and inhibitory cell type inputs to IS3 cells during a theta-cycle (***Bezaire et al., 2016; Mizuseki et al., 2009***) (***Figure 9***).

**Figure 9.**
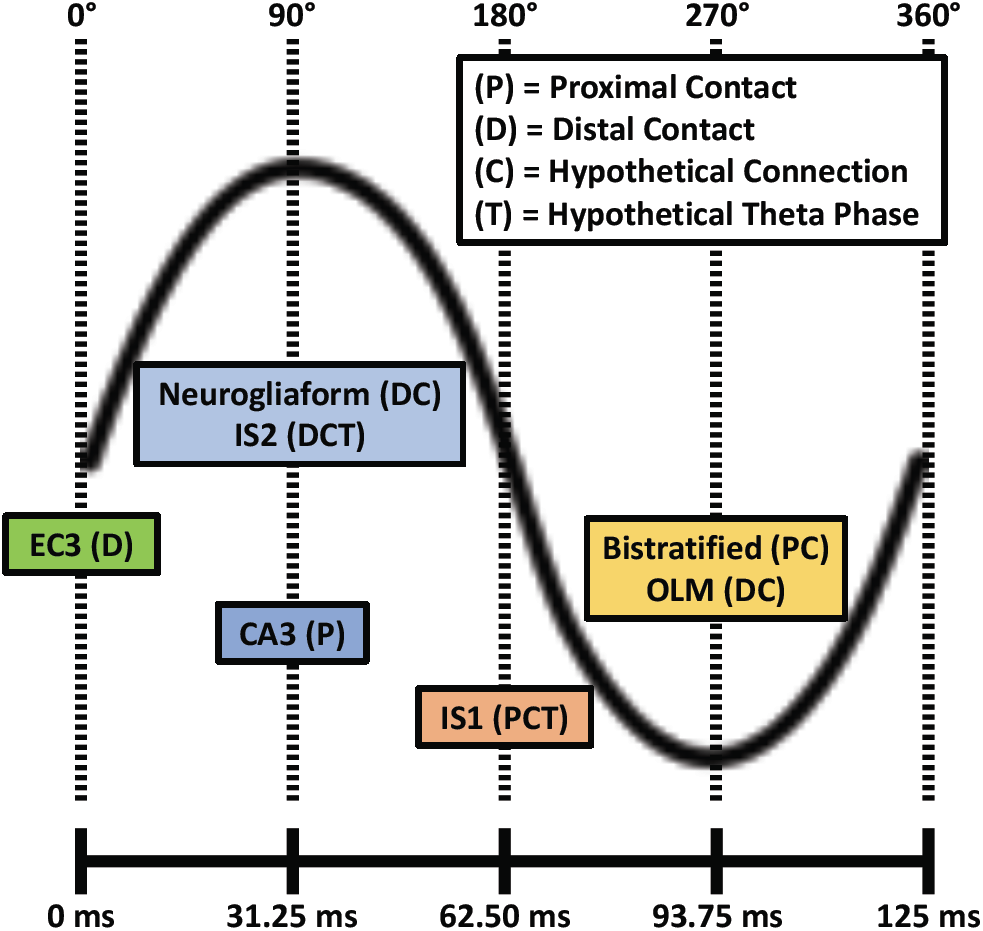
Schematic showing the relative timing of excitatory input populations (i.e. EC3 and CA3) and hypothetical local inhibitory input populations (e.g. bistratified, OLM, IS1, IS2, and neurogliaform) to IS3 cells. The legend denotes inputs that would connect to IS3 cell proximal dendrites (P), inputs that would connect to IS3 cell distal dendrites (D), connections that are hypothetical (i.e. non-confirmed; C), as well as inputs with relative timing during the theta-cycle phase that are hypothetical (T). For the input populations with hypothetical relative theta-cycle timing, we estimated that they would spike 1/4 of a phase after either CA3 or EC3 input populations spike, depending on where the dendrites of those cell types are likely to be positioned. EC3 = entorhinal cortex, layer III pyramidal cells; OLM = oriens lacunosum moleculare interneurons; IS1 = interneuron specific, type 1 interneurons; IS2 = interneuron specific, type 2 interneurons.

In ***Figure 10*** we show IS3 cell output when theta inputs are added to a low (LNI_LNE_LIS_LES) or to a high (HNI_HNE_HIS_HES) amount of synaptic input representative scenario. Also shown are the excitatory and inhibitory currents and conductances, computed in the same way as done for ***Figure 8***. Excitatory and inhibitory currents and conductances for all of the representative scenarios are shown in ***Figure 10, Figure Supplement 1***. We note that the scenarios with lower amounts of inputs (top plots in ***Figure 10***, and bottom left panels in ***Figure 10, Figure Supplement 1***) exhibit the most rhythmic currents when compared to scenarios with larger amounts of inputs. This is borne out when computing the power spectral densities (PSDs) of the spike trains before and after the addition of theta-timed inputs, as shown in ***Figure 10, Figure Supplement 2***. The representative scenario from pool LNI_LNE_LIS_LES (i.e. lowest amount of inputs) shows a much more appreciable increase in the PSD at 8 Hz when compared to the representative scenario from pool HNI_HNE_HIS_HES. As well, when looking at averaged excitatory and inhibitory conductances during a theta cycle, the low input representative scenario is more responsive to theta-timed inputs (***Figure 10, Figure Supplement 3***). As might be expected, excitation leads inhibition at the quarter cycle due its more proximal input locations (***Figure 9***) and faster synaptic time constants. Inhibitory or excitatory peaks at other quarter cycles are also clearly apparent in the representative scenario from pool LNI_LNE_LIS_LES. To ensure the robustness of our results, we re-do these theta-timed simulations for the representative scenario from pool LNI_LNE_LIS_LES five times and the averaged voltage output from them are shown in ***Figure 10, Figure Supplement 4***.

**Figure 10.**
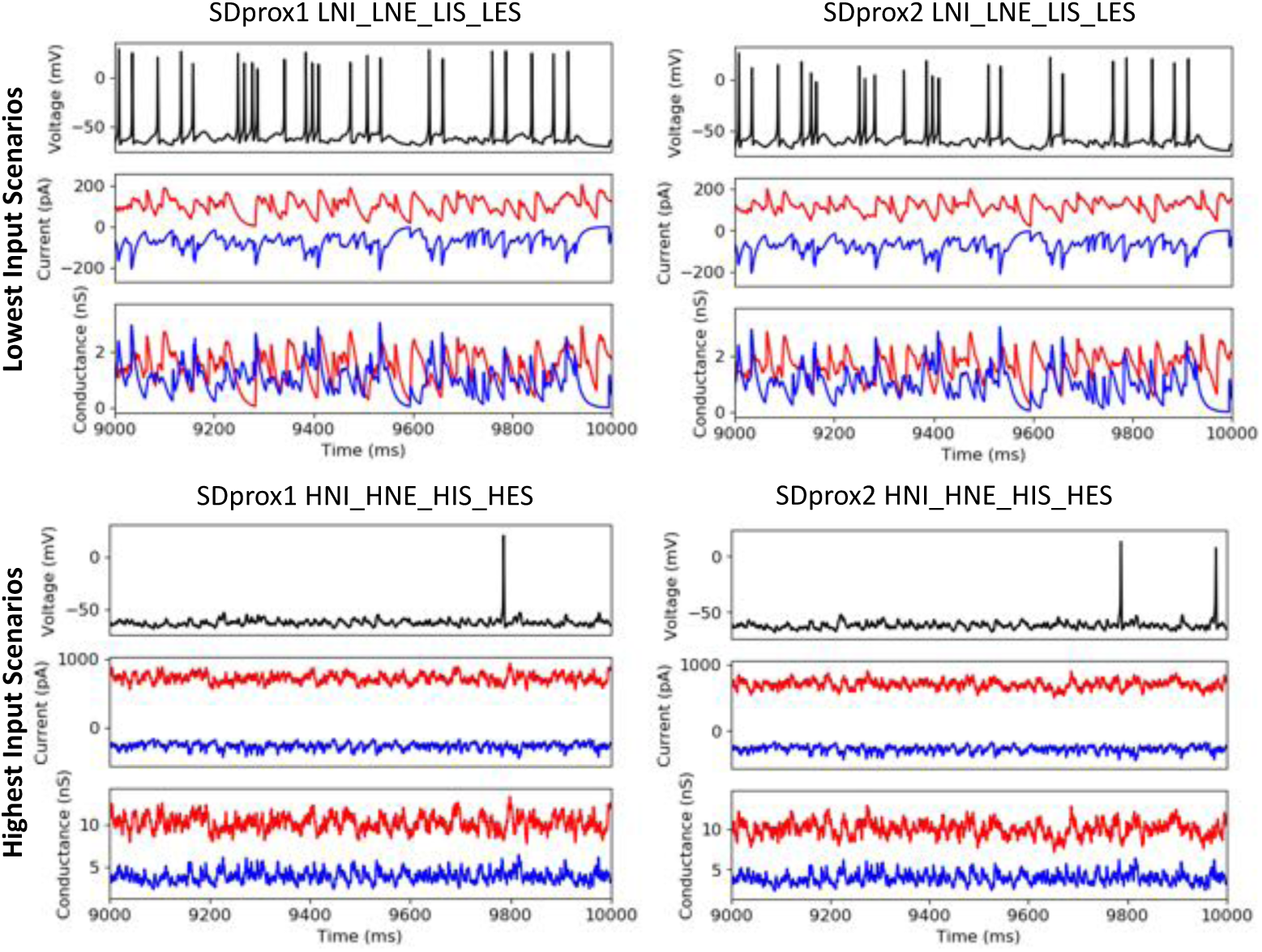
Example voltage traces, and inhibitory (red) and excitatory (blue) currents and conductances during the addition of theta timed inputs. The top row of plots show results using the representative HC scenarios with the lowest amounts of inputs (LNI_LNE_LIS_LES). The bottom row of plots show the results using the representative HC scenarios with the highest amount of inputs (HNI_HNE_HIS_HES).

These results suggest that the synaptic inputs in the baseline HC scenarios with larger amounts of inputs are drowning out the influence of the comparatively smaller theta-timed inputs. This can be appreciated by noting that the representative HC scenario with highest input resistance also responded the most to theta-timed inputs (***Figure 7***, top plots; see LNI_LNE_LIS_LES and LNI_LNE_LIS_HES for SDprox1; see LNI_LNE_LIS_LES, LNI_LNE_LIS_HES, and LNI_LNE_HIS_LES for SDprox2). Additionally, having inhibitory dominance when there are larger amounts of inputs will decrease neuron sensitivity to new input by increasing the threshold for spiking. In line with this, we observed considerably larger spike rates in the representative HC scenario with low amounts of inputs (***Figure 7***, bottom plots; see LNI_LNE_LIS_LES for both models). In summary, our findings demonstrate that having low amounts of inputs specifically in terms of number of synapses (i.e., LNI and LNE), combined with balanced inhibition and excitation, allow our IS3 cell models to exhibit both *in vivo*-like states as well as sensitivity to additional theta-timed inputs.

## Discussion

In this work we explored synaptic input parameter spaces that generate *in vivo*-like scenarios for IS3 cell models. We found that when there is a low amount of common inputs with equally balanced excitatory and inhibitory conductances, *in vivo*-like states are present as well as a sensitivity to additional theta-timed inputs. It was possible to have *in vivo*-like states with a larger amount of input, but then there is an inhibitory dominance and the sensitivity to additional inputs is lost. We found this to be due to lower input resistances with the larger synaptic input currents. It is interesting to note that IS3 cells have been found to have high input resistances relative to other cell types (***Tyan et al., 2014***). If IS3 cells are to preferentially control other inhibitory cell types at theta frequencies *in vivo* as they do *in vitro* (***Tyan et al., 2014***), then it is reasonable to expect that IS3 cells need to be able to be recruited with preferred firing at theta to be able to exert this influence. Therefore, we predict that IS3 cells *in vivo* have weak inputs with equally balanced conductances. We note that the vast majority of studies regarding *in vivo* excitatory-inhibitory balances have been in cortical circuits. In the hippocampus, ***Atallah and Scanziani*** (***2009***) show balancing of excitation and inhibition on pyramidal cells during gamma oscillation amplitude modulations. In cultured cells, a local structural excitation/inhibition balance on hippocampal pyramidal dendrites has been shown (***Liu, 2004***). Overall, our work suggests that balanced states exist in hippocampal circuits during rhythmic activities *in vivo*.

### Intrinsic properties and comparison to pyramidal cells

That the synaptic inputs could be low and still have an *in vivo*-like scenario is suggestive of the importance of *both* intrinsic and synaptic homeostatic balances of IS3 cells in their contribution to hippocampal circuitry. In our explorations we used two IS3 cell models from our previous work that captured IS3 cell features (***Guet-McCreight et al., 2016***). Due to its intrinsic properties, the SDprox2 model (A-type potassium channels restricted to soma) generated larger numbers of HC scenarios than the SDprox1 model (A-type potassium channels in dendrites), as well as a larger number of models nearing depolarization block. Although at present we do not know the distribution of A-type potassium channels in IS3 cells, their presence or absence in IS3 cell dendrites clearly would affect cell excitability. Further, our IS3 cell models currently have a minimal set of ion channel types and other channel types could clearly make additional contributions. For example, calcium channels are not included in our models even though there is evidence for them (***Vinet and Sík, 2006***), and they could potentially be distributed to more distal IS3 cell dendrites. If included, dendritic calcium channels could potentially serve to help amplify synaptic inputs and possibly shift the high input inhibitory-dominant HC regimes towards more balanced regimes. In essence, our prediction of low synaptic inputs *in vivo* highlights the importance of having a biophysical characterization of ion channels in IS3 cells.

Pyramidal cells have been studied much more extensively *in vivo* (***Destexhe and Paré, 1999; Harvey et al., 2009; Adesnik, 2017***) relative to inhibitory cells or specifically IS3 cells. Relative to their hippocampal counterparts, IS3 cells have much smaller synaptic densities (***Gulyás et al., 1999; Megias et al., 2001***), and thus we would expect pyramidal cells to have even lower input resistances *in vivo* due to larger synaptic densities. However, pyramidal cells are considerably larger than IS3 cells and so would have smaller axial resistances, which would promote signal propagation and better counteract the decreases in input resistance. As well, pyramidal cell dendrites have several active voltage-gated channels spread throughout the further lengths of their dendrites (***Magee et al., 1998***). This would also promote signal propagation along pyramidal cell dendrites relative to IS3 cells, which only have spike-propagating active properties in proximal dendrites as modelled here.

### Theoretical and biological balances

Although principles of brain coding remain to be clearly deciphered, they involve firing rates, spike timing and correlations, and repeating patterns in various capacities (***Luczak et al., 2015***). The variability of neural firing increases information content and theoretical studies have shown that networks with balanced excitation and inhibition exhibit irregular spiking or an asynchronous state (***Vreeswijk and Sompolinsky, 1996***). Correlations in these balanced networks can vary depending on spatial connectivities (***Rosenbaum et al., 2017***). Neurons would be considered to be in a fluctuation-driven regime, as opposed to a mean-driven regime (***Schreiber et al., 2009***), with spike firing occurring randomly due to a loose balance between excitation and inhibition. From a coding perspective, efficiency and information content favours having neither an excitatory nor an inhibitory dominance (***Denève and Machens, 2016; Zhou and Yu, 2018***). Further, having a loose or global excitatory/inhibitory balance is thought to promote rate coding (***Denève and Machens, 2016; Shadlen and Newsome, 1994; Zhou and Yu, 2018***) and having a tight balance would facilitate spike time coding. Tight balances between excitation and inhibition have been observed *in vitro* (***Xue et al., 2014***), during rhythmic contexts (***Atallah and Scanziani, 2009; Huh et al., 2016***), as well as *in vivo* (***Higley and Contreras, 2006; Monier et al., 2008; Adesnik, 2017***). How tightly excitation and inhibition are balanced *in vivo* would depend on the specificities of the anatomical circuit and the behavioural context. From an individual cell perspective, experimental studies indicate that excitation and inhibition are proportionally balanced *in vivo* (***Haider et al., 2006***), although there could also be an inhibitory dominance (***Piwkowska et al., 2008***). However, extracting these balances from experiment can be tricky due to space clamp and reversal potential estimations. We directly showed the effect of space clamp in excitatory and inhibitory balances in our explorations here where excitatory conductances were underestimated.

In turtle lumbar spinal circuits, it has been reported that neurons spend a component of their time in fluctuation-driven regimes and another component in mean-driven regimes (***Petersen and Berg, 2016***). Neurons in fluctuation-driven regimes were found to have enhanced sensitivity, whereas neurons in mean-driven regimes had reduced sensitivity. This highlighted a network-level ‘Goldilocks Zone’ where neurons could be both stable and sensitive to new inputs (***Humphries, 2016***). Our *in vivo*-like states are suggestive of such a scenario in hippocampal circuits as we have found fluctuation-driven regime with enhanced sensitivity to additional inputs for IS3 cells. Increased excitation would move one toward a mean-driven regime. We achieved our *in vivo*-like state using a loose balance between excitation and inhibition, but tighter balances are warranted when introducing a rhythmic or transient context such as during theta rhythms or sharp wave-associated ripples (***Klausberger and Somogyi, 2008; Huh et al., 2016; Bezaire et al., 2016***).

The prominent theta rhythm in the hippocampus, associated with spatial navigation and memory consolidation, is observed during REM sleep and movement (***Buzsáki, 2002; Colgin, 2016***). In CA1 this rhythm is generated through a combination of external inputs from entorhinal cortex and CA3 (***Kamondi et al., 1998; Mizuseki et al., 2009***), recurrent inputs from medial septum (***Boyce et al., 2016***), as well as intrinsically (***Goutagny et al., 2009***) through inhibitory cell interactions (***Ferguson et al., 2015; Huh et al., 2016***). During intrinsically generated theta rhythms in CA1, there is a tight balance between excitation and inhibition for different neuron types (***Huh et al., 2016***). In our explorations we found that common inputs were important in generating *in vivo*-like states for our IS3 cells as there was a noticeable increase in the size of the parameter spaces that generated high-conductance, fluctuation-driven regimes when common inputs were present. This suggests that correlations might be generically present in hippocampal microcircuits. Our prediction of evenly balanced excitation and inhibition of *low* amounts to IS3 cells confers high input resistances and sensitivity to additional theta-timed inputs.

### Balance and disease states

In considering disease states, a consideration of excitation/inhibition balance on its own is over-simplistic (***Marín, 2012***). For example, it was recently shown in a mouse model of Fragile-X Syndrome that correlations and firing rate changes were insufficient to capture the shifts seen in the population activities with disease when comparing a spiking circuit model with analyses of calcium imaging data (***O’Donnell et al., 2017***).

While we found balanced regimes representative of neural states in live animals, we also highlighted inhibitory-dominant and excitatory-dominant regimes in our models that could be present during disease states. For example, at many levels of inputs (i.e. the different parameter range pools), excitatory-dominant input scenarios can generate models nearing depolarization block, thus demonstrating that excitatory shifts in balance could lead to inactivation of particular cell types via depolarization block if given enough excitation. This was suggested to be the case for fast spiking interneurons during the spread of epileptic waves across cortex (***Trevelyan et al., 2006; Ahmed et al., 2014; Ho and Truccolo, 2016***). Interestingly, inhibitory drive from IS3 cells has been shown to be reduced in a pilocarpine-induced model of epilepsy (***David and Topolnik, 2017***), suggesting a shift towards excitatory-dominant, depolarization block-generating regimes in IS3 cell postsynaptic targets.

### Limitations

Although we have done an extensive exploration to examine how much and what balance of excitatory and inhibitory synaptic inputs IS3 cells might receive *in vivo*, we did limit our exploration by grossly separating proximal and distal excitatory inputs and not controlling the spatial locations of common input. However, that we separate proximal and distal locations at all will allow us to examine how inputs from different regions could control IS3 cells during different behavioural states. We based our HC metric from other cell types since *in vivo* information is not available for IS3 cells directly although our consideration of threshold values did take IS3 cell specifics into consideration. While changing the specific threshold values used in our HC metric would change the details of our results, we do not expect that our general results that predict low amounts of equally balanced excitation and inhibition to change considering our examination of unpacking the components of the HC metric.

We used a simple first-order synaptic kinetic model to simulate our postsynaptic effects. Although there is evidence that short-term synaptic depression exists for inputs onto VIP+ cells in neocortex (***Karnani et al., 2016b***), it is unknown whether this is also the case for VIP+ IS3 cells in hippocampus. Given the high levels of random activity in our simulations, it is unclear how implementing synaptic depression would change our results, but it is likely that the overall synaptic currents and conductances generated during higher spike rate input regimes will be smaller. As well, in our synaptic model, we assume that there is a 0% failure rate (i.e. all common synapses succeed at transmitting an EPSC/IPSC at the same time), which is unlikely to be the case. If implemented, this would probably reduce the number of HC scenarios being generated, as failure rates would decrease the probability of postsynaptic spiking. Finally, we have noted that our cellular models are limited in the types of ion channels incorporated, but their development ensured that IS3 cell features were captured (***Guet-McCreight et al., 2016***).

### Concluding remarks and future work

While excitation and inhibition balance is central to brain coding, it is important to reconcile the specifics associated with theoretical insights and biological realities. By doing an extensive computational exploration here we have been able to *both* predict synaptic inputs to IS3 cells *in vivo* as well as to unpack some of these ‘balance’ aspects.

Using our predicted *in vivo*-like states we will investigate the conditions and contexts under which IS3 cells can be recruited to spike and contribute to theta rhythms and sharp wave-associated ripples in the hippocampus. Ongoing experimental work shows promise in recording from IS3 cells *in vivo* (***Villette et al., 2017***) and our modeling work can help guide such studies. In combination, we aim to achieve an understanding of how IS3 cells could critically contribute to hippocampal activities and behaviours. As IS3 cells specifically target OLM cells, we also aim to explore *in vivo*-like states in previously developed OLM cell models (***Sekulic and Skinner, 2017***) and create *in vivo*-like microcircuits. Similar to cortex, perhaps IS3 cells “open holes in the blanket of inhibition” (***Karnani et al., 2016a***) to bring about and control theta rhythms and sharp wave-associated ripples in the hippocampus.

## Methods

### IS3 cell models

We use two multi-compartment model variants of IS3 cells developed in our previous work (***Guet-McCreight et al., 2016***) using the NEURON software environment (***Carnevale and Hines, 2006***). The morphological and electrophysiological properties of the models are directly based on IS3 cell recordings. The model consists of 221 somatic, dendritic, and axon initial segment compartments (see ***Figure 1, Figure Supplement 2***B), and exhibit firing frequency and depolarization block characteristics of IS3 cells. The ion channels types and distributions in our IS3 cell models reflect interneuron specific characteristics as channel data specific to IS3 cells is not available at present. They include transient and persistent sodium channels and A-type and delayed rectifier potassium channels. Also, to reflect subthreshold activities observed in IS3 cells, intrinsic noise was included as done in our previous models (***Morin et al., 2010***). Although it is expected that there are more ion channel types in IS3 cells, we did not include any additional ones in building these models, but rather focused on a minimal set since this was sufficient to capture IS3 cell features. Direct immunological evidence was obtained for delayed rectifier potassium channels in dendritic portions of IS3 cells but it is at present unknown about other channel types (***Guet-McCreight et al., 2016***). Here, we use two of our models that captured IS3 cell electrophysiological features, one that has A-type potassium channels in the dendrites (SDprox1) and one that does not (SDprox2). The specifics of these models with their somato-dendritic ion channel distributions are given in ***Table 3***.

**Table 3.** Intrinsic properties of IS3 cell model variants. *G*_*Na,t*_ = transient sodium channel conductance; *G*_*Na,p*_ = persistent sodium channel conductance; *G*_*Ka*_ = A-type potassium channel conductance; *G*_*Kdrf*_ = fast delayed rectifier potassium channel conductance. **Table 3–source data 1.** *Guet-McCreight et al*. (*2016*)

| Model | Distribution | $G_{Na,t}$<br>(mS/cm<sup>2</sup>) | $G_{Na,p}$<br>(mS/cm<sup>2</sup>) | $G_{Ka}$<br>(mS/cm<sup>2</sup>) | $G_{Kdrf}$<br>(mS/cm<sup>2</sup>) |
| --- | --- | --- | --- | --- | --- |
| SDprox1 | Uniform across soma and first 70 μm of dendrites<br>($G_{Na,p}$: soma only) | 70 | 0.075 | 70 | 250 |
| SDprox2 | Uniform across soma and first 70 μm of dendrites<br>($G_{Na,p}$ and $G_{Ka}$: soma only) | 55 | 0.15 | 30 | 295 |

### Synaptic model and parameters

We use excitatory and inhibitory synaptic parameters that were obtained from optimal fits to the experimental data (***Guet-McCreight et al., 2017***). We use NEURON’s Exp2Syn synapse model, which describes synapses as two-state kinetic schemes:

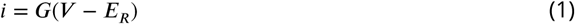

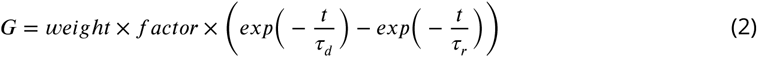

Where *i* is the synaptic current, *G* is the synaptic conductance, *E* is the reversal potential, *V* is the membrane potential, *weight* is the synaptic weight, *factor* is a NEURON process that is used to normalize the peak synaptic conductance to the value of *weight, t* is time, *τ*_*r*_ is the rise time, and *τ*_*d*_ is the decay time. Both model variants (SDprox1, SDprox2) were used in obtaining synaptic parameters as derived from optimizations that fit passive model responses to those seen in layer-specific-evoked excitatory postsynaptic currents (EPSCs) and spontaneous inhibitory postsynaptic currents (IPSCs) (***Guet-McCreight et al., 2017***). The layer-specific evoked EPSCs were obtained under minimal stimulation. Evoked EPSCs generated from stimulation in stratum radiatum were assumed to be Schaeffer collateral inputs from CA3 to IS3 cell proximal dendrites. Evoked EPSC responses generated from stimulation in stratum lacunsom-moleculare were assumed to be inputs from entorhinal cortex layer III (ECIII) to IS3 cell distal dendrites (***Witter, 2010***).

For excitatory synapse models, we applied a reversal potential of 0 mV, and applied a linear distance-dependent weight rule, according to the weight of the best fit for proximal dendrites (i.e. < 300 µm from the soma), and the weight of the best fit for distal dendrites (i.e. > 300 µm from the soma). This border was based on the layout of the dendritic morphology of our reconstructed models, with proximal dendrites in the Stratum Radiatum (SR), and distal dendrites extending to the Stratum Lacunosum Moleculare (SLM). Excitatory rise and decay time constants (i.e. *τ*_*r*_ and *τ*_*d*_) for either proximal or distal synapses were then fixed to the optimized time constants of the best fit in proximal or distal dendrites, respectively.

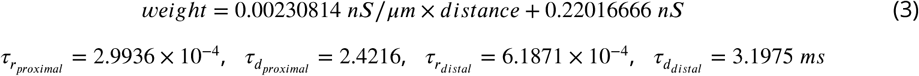

For the inhibitory synapse models, we applied a reversal potential of -70 mV, and applied a linear-distance-dependent weight rule, according to the optimized weight values of the two best fits, regardless of their synaptic locations. Inhibitory time constants for all synaptic locations in the model were then fixed to the optimized time constants of the best inhibitory synapse fit.

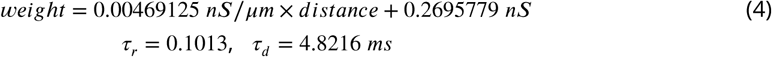

More details on synaptic parameter optimizations are given in ***Guet-McCreight et al***. (***2017***).

### Parameter exploration ranges for excitatory and inhibitory synapses, strategy and setup details

In our parameter explorations, we keep the individual synaptic amplitude and time course parameters (i.e. weight, rise times, and decay times) fixed, see above section. However, we vary the inhibitory and excitatory spike rates and numbers of synapses (i.e. four parameters in total). Since these parameters are variable *in vivo*, depending on the context and the neuron type of interest, we aim to explore full ranges of possible input parameter combinations, based on what is known experimentally.

#### Excitatory spike rates

We first looked at the spike rate distributions of CA3 pyramidal cells, as well as ECIII pyramidal cells. While CA3 pyramidal cells tend to occasionally show high spike rate distributions (i.e. >100 Hz) during anaesthesia, which could correspond to bursting activity (***Frerking et al., 2005; Kowalski et al., 2016***), on average, spike rates tend to be below 30 Hz, with peak firing rates occurring during behavior while the animal moves through place fields (***Leutgeb et al., 2004***) or time fields (***Salz et al., 2016***). Similarly, pyramidal cells in ECIII (i.e. in both lateral and medial ECIII) exhibit below 30 Hz spiking during 200 pA stimulation *in vitro* (***Canto and Witter, 2012b***,a), as well as during grid field traversals *in vivo* (***Domnisoru et al., 2013***). Considering these observations, we set maximum excitatory spike rates of 30 Hz for both proximal and distal excitatory inputs.

#### Inhibitory spike rates

Though the spike rates of fast-spiking parvalbumin-positive (PV+) cells *in vivo* during theta or low oscillations tend to be below 40 Hz, during sharp wave-associated ripples these tend to spike at up to 160 Hz (***Katona et al., 2014***). Since it is unclear which groupings of local CA1 inhibitory neurons target IS3 cells and we are simulating baseline *in vivo* states (i.e. without sharp-wave associated ripples), we set the maximum inhibitory spike rate to be 100 Hz.

#### Synapse numbers

The maximum synapse numbers in the model were deduced from excitatory and inhibitory synaptic densities onto calretinin-positive (CR+) cells in hippocampal CA1 (***Gulyás et al., 1999***). Using these maximal densities, as well as the average length of each compartment in the IS3 cell model, we estimated there to be approximately 9 excitatory synapses and 2 inhibitory synapses per compartment. In total this estimated maximums of 1530 excitatory synapses and 344 inhibitory synapses spread across the dendritic arbor of the model.

We also investigate whether synaptic inputs are common or independent (***Figure 1, Figure Supplement 2****A*). Inputs that are common means that a presynaptic input forms multiple synaptic inputs onto the IS3 cell model, and thus these common inputs are assigned identical spike trains. Independent inputs, simply means that each synapse has a unique spike train (i.e., a different random seed is used). For common input parameter searches, we use numbers of 9 common excitatory inputs and 4 common inhibitory inputs, although different numbers are also examined (see below). These numbers are chosen because a) they fall well within ranges of common inputs seen in somatosensory cortex according to findings by Blue Brain Project (***Ramaswamy et al., 2015; Reimann et al., 2015***), and b) they allow a high parameter search resolution based on the maximal excitatory and inhibitory synapses. That is, if looking at larger numbers of common inputs, the parameter search resolution becomes smaller because each time we increase the number of synapses we have to add an amount of synapses equal to or divisible by the number of common inputs. Note that the maximum numbers of excitatory and inhibitory synapses are 1530 and 344 respectively, which are exactly divisible by 9 and 4. The parameter ranges explored in our parameter searches are shown in ***Table 4***.

**Table 4.**
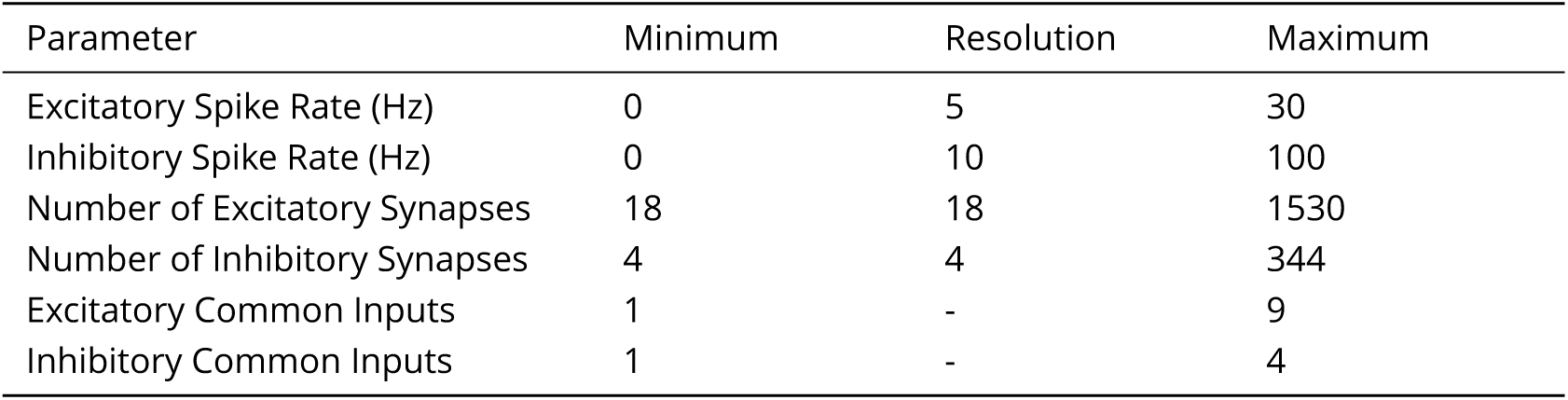
Parameter exploration ranges. Output from a total of 4,502,960 ten-second simulations are obtained when considering all possible parameter combinations and the two model variants.

We perform our parameter search in the following manner. The locations of inhibitory and excitatory synapses are chosen randomly but with proximal and distal locations having an equal number of excitatory synapses. Throughout our parameter explorations, we adjust the number of synapses by selecting synapses and assigning them spike trains. Each presynaptic spike train is obtained by sampling spike times randomly from a uniform distribution. The mean frequency (spike rate) of the presynaptic spike train is set by ensuring that there is an appropriate number of spikes for the length of time of the simulation. For example, for 100Hz, 1,000 spike times would be sampled for a 10 second simulation. This presynaptic spike rate is adjusted by changing the number of spike times being sampled. Further implementation details are provided in the Methods. In ***Figure 1, Figure Supplement 2****B,C* we show an example for one set of parameters - 8 inhibitory synapses activated with 30 Hz presynaptic spike trains, and 144 excitatory synapses activated with 5 Hz spike trains. ***Figure 1, Figure Supplement 2****B* shows the random distribution of 8 inhibitory (red), 72 proximal excitatory (blue) and 72 distal excitatory (green) synapses on the dendritic tree of the IS3 cell model, and ***Figure 1, Figure Supplement 2****C* shows a raster plot of the presynaptic spike trains. In this example set of parameters, the common input scenario (***Figure 1, Figure Supplement 2****A*, left) is shown.

### Implementation details

The synapse locations are determined as follows: The 1530 excitatory and 344 inhibitory synapses are fully distributed throughout the dendritic tree with 9 excitatory and 4 inhibitory synapses per compartment. To activate the given number of synapses being explored, vectors of presynaptic spike times are assigned to the selected synapses using NEURON’s VecStim function. Non-assigned synapses remain silent over the course of the simulation since they do not receive any input spike trains. The locations of the active synapses are determined using a customized function for random index selection, which chooses indices by sampling integers from a discrete uniform distribution. In this function each group of synapses are organized in vectors (proximal excitatory, distal excitatory, and inhibitory). For each synapse vector a random indexing vector of the same length is created where the content of each index (in the random indexing vector) is an integer value sampled using NEURON’s random.discunif(min,max). A loop that keeps sampling from random.discunif(min,max) is done until every number in the random indexing vector is unique, so that synapse locations are not represented more than once.

Each presynaptic spike train is determined by sampling spike times from a uniform distribution of length 0 to 10,000 msec (the simulation length that we use). We use NEURON’s default random number generator, which is a variant of the Linear Congruential Generator combined with a Fibonacci Additive Congruential Generator (ACG), to sample from a discrete uniform distribution. The spike rate was adjusted by changing the number of spike times being sampled within this time window. Once the spike times are selected and sorted, we use NEURON’s VecStim function to assign the custom-built spike time vectors into the different synaptic locations. The spike trains are irregular and our spike rates are given by # of spikes/10 (Hz), since our simulation lengths are 10 seconds.

To obtain a representative scenario from each pool, we use the following procedure: For each pool, we go through the list of HC scenarios obtained for that pool (see ***Figure 4***). The order in which we do this is, excitatory spike rates ⇒ inhibitory spike rates ⇒ number of excitatory synapses ⇒ number of inhibitory synapses, from lowest to highest values. For each scenario, a 10 second simulation is done up to 10 times, where on each iteration, presynaptic spike times and locations of active synapses are re-sampled using new random seed values. If on any of the 10 iterations, the input scenario does not satisfy the criteria for HC, we skip to the next HC scenario in the list and start over. If an HC scenario satisfies the criteria for high-conductance 10 times in a row, we choose that scenario as the representative scenario for that pool, and move on to the next pool.

Output measurements of spike amplitude, coefficient of variation of the inter-spike-interval (*ISICV*) and number of spikes are acquired using the Efel module in Python (see http://bluebrain.github.io/eFEL/efeature-documentation.pdf), as well as custom python code for other measurements. Specifically, measurements for subthreshold voltage mean and standard deviation, where Efel code for finding the spike-begin and spike-end indices is used to cut out all spikes from the traces in order to be left with the subthreshold portions, from which means and standard deviations are computed. All custom codes are accessible through https://github.com/FKSkinnerLab/IS3-Cell-Model. Clutter-based dimensional reordering (CBDR) plots were generated in Matlab using code obtained from Adam Taylor (***Taylor et al., 2006***). An illustrative example CBDR plot is shown in ***Figure 2***.

Input resistance, *R*_*i*_, is computed using an injected current, *I*_*inj*_, of -100 pA starting at 4.5 sec and lasting for 1 sec in the 10 sec simulation. The difference between the mean voltage during hyperpolarization and the mean voltage during baseline (0-4.5 sec, and 5.5-10 sec, with spikes cut) is divided by the current input magnitude: *R*_*i*_ = (*V*_*m,hyperpolarized*_ − *V*_*m,baseline*_)/*I*_*inj*_.

Power spectral densities (PSDs) are computed from the model’s spike train in the last 9 seconds of the simulation, using the Scipy module in Python for estimating the PSD using Welch’s method. For our bar plots, we sample the PSD magnitude specifically at 8 Hz.

Parameter explorations are done via simulations using NEURON’s Python interface through the Neuroscience Gateway (NSG) for high-performance computing (***Sivagnanam et al., 2013***).

### High-conductance (HC) metric

The IS3 cell model outputs from our simulations are analyzed for the last 9 seconds of our 10 second simulations. The measurements include average subthreshold membrane potential 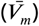, standard deviation of the subthreshold membrane potential 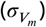, the interspike interval coefficient of variation (*ISICV*), and the average spike amplitude 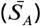. We use these four measurements in each simulation to compute our “high-conductance” (HC) metric as a conditional statement.

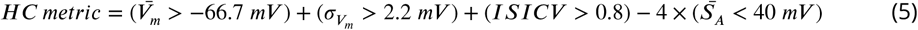

Using this equation we identify synaptic input scenarios that are high-conductance (HC metric = 3), non-high-conductance (NHC; HC metric = 0), as well as models nearing or in depolarization block (DB; where the average spike amplitude is less than 40 mV, making the HC metric negative). Note that the NHC state is interpreted as not being in DB and not satisfying any of the other conditions.

We chose the threshold values in our metric based on literature values seen in other cell types *in vivo*. For example, we know from ***Destexhe and Paré (1999)*** that neocortical pyramidal cells *in vivo* show elevated average subthreshold membrane potentials and subthreshold membrane potential fluctuations. We specify that the subthreshold membrane potential standard deviation should be greater than 2.2 mV. This is based on there being a 10-fold decrease in cat neocortical neuron membrane potential standard deviations *in vivo* upon application of tetrodotoxin (***Destexhe and Paré, 1999***). Since the membrane potential standard deviation of the IS3 cell model without synapses is 0.22 mV (note that there is intrinsic noise applied at the soma), we could therefore expect a 10-fold increase (i.e. 2.2 mV) in an *in vivo*-like scenario. We specify that the average subthreshold membrane potential should be greater than -66.7 mV. This is based on the change in membrane potential of mouse CA1 pyramidal cells during place field entry (***Harvey et al., 2009***), where we would expect to see at minimum, an approximate 3 mV depolarization in the membrane potential. Alternatively, if basing this criterion on results from neocortical cells in cats from ***Destexhe and Paré (1999)***, we would expect a depolarization of at least 10 mV (i.e. an average membrane potential greater than -60 mV in our model). We also know that neurons *in vivo* tend to show more irregular spiking activity (***Destexhe et al., 2003***). We specify that the interspike interval coefficient of variation (*ISICV*) must be greater than 0.8. This is based on a lower limit for the *ISICV* of neurons in middle temporal cortex of alert macaque monkeys during constant-motion stimulation (***Stevens and Zador, 1998***). Alternatively, for neurons in visual cortex and middle temporal cortex of cats and macaque monkeys, it had also been found that *ISICV*s greater than 0.5 could be expected *in vivo* for neurons that are spiking above 30 Hz (***Softky and Koch, 1993***).

### Excitatory and inhibitory balance metrics

To get a sense of the balance between excitation and inhibition in our explorations, we define two EI metrics. In the first metric (***Equation 6***), we normalize each parameter to their maximal possible value and scale the range such that values fall between -1 (i.e. purely inhibitory-dominant) and 1 (i.e. purely excitatory-dominant).

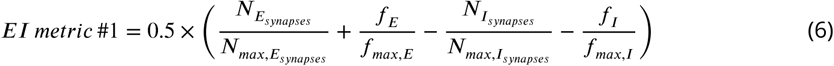

Where *N* is number of synapses and *f* is spike rate for excitatory (*E*) and inhibitory (*I*) inputs. The “max” stands for the maximum value looked at within the parameter search (***Table 4***). In the second metric (***Equation 7***) we chose not to normalize them at all as this can unfairly penalize parameters with larger parameter ranges. Note however that while excitatory synapses have larger parameter ranges for numbers of synapses, inhibitory synapses have larger parameter ranges for presynaptic spike rates, so this possibly balances out the normalization. As a secondary E/I balance metric, we chose to simply look at the difference in cumulative spike rates across all excitatory and inhibitory synapses. Once again, any negative values represent inhibitory-dominant parameter combinations, and any positive values represent excitatory-dominant regimes.

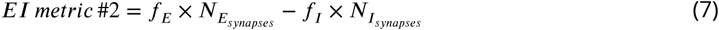

### Isolating excitatory and inhibitory currents

To isolate the excitatory and inhibitory currents in our IS3 model cells, we run the simulation twice, a first time where we remove inhibitory inputs, and a second time where we remove excitatory inputs. We apply a voltage clamp at the soma of the model (NEURON’s VClamp), set the leak reversal potential in all compartments to the holding potential (-70 mV for recording excitatory currents, and 0 mV for recording inhibitory currents), and remove ion channel conductances in all compartments. We also convert current traces into conductance traces according to: *G*_*trace*_ = *I*_*trace*_/(*V*_*hold*_ − *E*_*R*_)

This protocol is similar, but not identical, to experimental protocols that measure excitatory and inhibitory currents without the use of synaptic blockers (***Atallah and Scanziani, 2009; Xue et al., 2014; Huh et al., 2016***). One notable difference is that during experiments excitatory or inhibitory currents are not removed entirely using these experimental protocols (unlike what we can easily do in our simulations by not giving input spike trains to the synapses). Rather, their activity is simply rendered ineffective by voltage clamping the cell at their supposed reversal potentials (***Zhou and Yu, 2018***). To examine how outputs might differ, we also isolate excitatory and inhibitory currents by doing the voltage clamp and without removing excitatory or inhibitory inputs directly (***Figure 8, Figure Supplement 1***).

### Estimating theta-timed synapse numbers

To determine how many excitatory and inhibitory synapses should be used for theta-timed inputs (8 Hz), we estimate the number of excitatory and inhibitory proximal and distal inputs that are necessary to elicit theta-timed spiking outputs in the *in vitro* models (i.e. when the model IS3 cell is not in an *in vivo*-like state). We do the following: (1) Approximate the number of synaptic inputs for excitatory and inhibitory synapses in either proximal (i.e. <300 *µm* from the soma) or distal (i.e. >300 *µm* from the soma) dendrites independently of each other. (2) For excitatory inputs, the number of inputs is increased until the IS3 cell models’ spike train PSD at 8 Hz is larger than 50 spikes2_Hz and there are more than 10 spikes. (3) To ensure comparability between SDprox1 and SDprox2 when estimating numbers of inhibitory inputs, the magnitude of the current injection in each model is first estimated by gradually increasing the current amplitude until a spike rate of at least 35 Hz (i.e. 350 spikes in 10 seconds) is obtained in both models. Using this method, the current clamp magnitude was determined to be 27.5 pA for SDprox1 and 24 pA for SDprox2. (4) For inhibitory inputs, the number of inputs is increased until the spike train PSD at 8 Hz is larger than 80 *spikes*2/*Hz* and there are less than 240 spikes. The results from doing this is given in ***Table 5***. Given these estimates, we apply 27 excitatory theta-timed synapses per layer-specific excitatory population (i.e. CA3 and ECIII, in ***Figure 9***). Likewise, for inhibitory cell populations we chose to apply 8 synapses per layer-specific population (i.e. bistratified, neurogliaform, OLM, IS1, and IS2, in ***Figure 9***).

**Table 5.** Estimates of numbers of layer-specific excitatory and inhibitory theta-timed inputs required to recruit the starting IS3 cell models to spike appreciably at 8 Hz

| Model | Proximal<br>Excitatory | Proximal<br>Inhibitory | Distal<br>Excitatory | Distal<br>Inhibitory |
| --- | --- | --- | --- | --- |
| SDprox1 | 27 Synapses | 8 Synapses | 27 Synapses | 8 Synapses |
| SDprox2 | 27 Synapses | 4 Synapses | 18 Synapses | 4 Synapses |

### Further explorations

#### Intrinsic cell noise

As described above, there are several sources of noise caused by random processes in our parameter explorations. In addition to the random choosing of synaptic locations and the random choosing of presynaptic spike times, our SDprox1 and SDprox2 models have Gaussian noise current injected directly at the soma. This was originally included in our IS3 cell models in order to capture subthreshold membrane potential fluctuations that are seen experimentally (***Guet-McCreight et al., 2016***). To ensure that this intrinsic noise in our models was not having a major effect on our our HC state scenarios, we re-did our simulations without the intrinsic noise while keeping the random seeds for synaptic locations and presynaptic spike times fixed. As shown in ***Figure 6, Figure Supplement 3***, this did not affect the consistency of our HC scenario results.

#### Changing the number of common inputs

As described above, we estimate the number of common inputs to use for excitatory and inhibitory synapses at 9 and 4 respectively, and our examinations focused on scenarios with common inputs as they promoted *in vivo*-like states (see Results). However, these particular numbers are estimates and it is unclear whether they might strongly affect *in vivo*-like states. Thus, as described below, we did further investigations using our robust, representative scenarios. We varied the number of excitatory and inhibitory common inputs from 1 (i.e. independent) to 10, and in these additional parameter explorations we kept the random seed values fixed between simulations so that only changes in the numbers of common inputs could affect the output of the simulations. The results are shown in ***Figure 6, Figure Supplement 4, Figure Supplement 5, Figure Supplement 6, Figure Supplement 7, Figure Supplement 8***. We note that there is a bit of inexactitude in these additional simulations. That is, if the total number of active synapses in the representative HC scenario is not divisible by the number of common inputs, this leaves a remainder group of synapses of a size smaller than the number of common inputs. An example schematic of this is shown in ***Figure 6, Figure Supplement 9***. Another inexactitude with these parameter searches is that if the total number of synapses is smaller than the set number of common inputs, then increasing the number of common inputs to a larger value than the total number of available synapses would essentially change nothing (e.g. see the inhibitory common inputs axis in bottom left panel figures of ***Figure 6, Figure Supplement 5***).

From this analysis, we further confirmed that the number of common excitatory inputs does in fact have a strong influence on our HC metric (***Figure 6, Figure Supplement 4***), whereas the impact of the number of common inhibitory inputs is comparatively milder. Interestingly, the number of common inputs shows an impact not only on the subthreshold membrane potential standard deviation (***Figure 6, Figure Supplement 6***), but also on the mean subthreshold membrane potential (***Figure 6, Figure Supplement 5***), and the mean spike rate (***Figure 6, Figure Supplement 8***). Counter-intuitively,the subthreshold membrane potential appears to mildly decrease as the number of common excitatory inputs is increased. This may be because when excitatory presynaptic spikes are more distributed (i.e. more independent) the membrane potential will be more consistently larger. On the other hand, when excitatory presynaptic spike times are correlated (i.e. more common), the increase in membrane potential caused by synaptic events will be more occasional and transient (and possibly even be partially removed from the analysis when spikes are cut from the traces), leaving the mean subthreshold membrane potential more hyperpolarized.

As was observed previously (i.e. ***Figure 3, Figure Supplement 2***), the subthreshold membrane potential standard deviation increases as the number of excitatory (and in some cases inhibitory) common inputs is increased (***Figure 6, Figure Supplement 6***). This is likely due to presynaptic spike trains causing larger deflections in the model’s membrane potential when there are larger numbers of common inputs. On the other hand, here we do not see any clear relationship between number of common inputs and the *ISICV* values, aside from occasionally regulating a border at which *ISICV* values jump from values of zero to values typically larger than 0.5 (***Figure 6, Figure Supplement 7***). In fact, these areas in the parameter space with *ISICV* values of zero appear to correspond with areas of the parameter space where there is no spiking present (i.e. ***Figure 6, Figure Supplement 8***). Coincidently, in ***Figure 6, Figure Supplement 8*** we observed a positive relationship between mean spike rate and number of common inputs, which demonstrates that having common correlated synaptic inputs will increase the probability of spiking. This finding is in line with work from ***Stevens and Zador (1998)*** which shows that inputs need to be correlated in order to generate the irregular spiking that is often seen *in vivo*. Furthermore, the common input numbers that we use in the main text Results (i.e. 9 common excitatory inputs, and 4 common inhibitory inputs) fall within the parameter regimes that generate HC metric values of 3.

## Acknowledgments

We would like to thank Dr. L. Topolnik and her lab for feedback and their excellent collaboration, and Dr. J. Lefebvre and A. Chatzikalymniou for comments on the manuscript. Special thanks to A. Sherrington for contributing the “Hippocampus” drawing in ***Figure 1***.

---

**Figure 1–Figure supplement 1.**
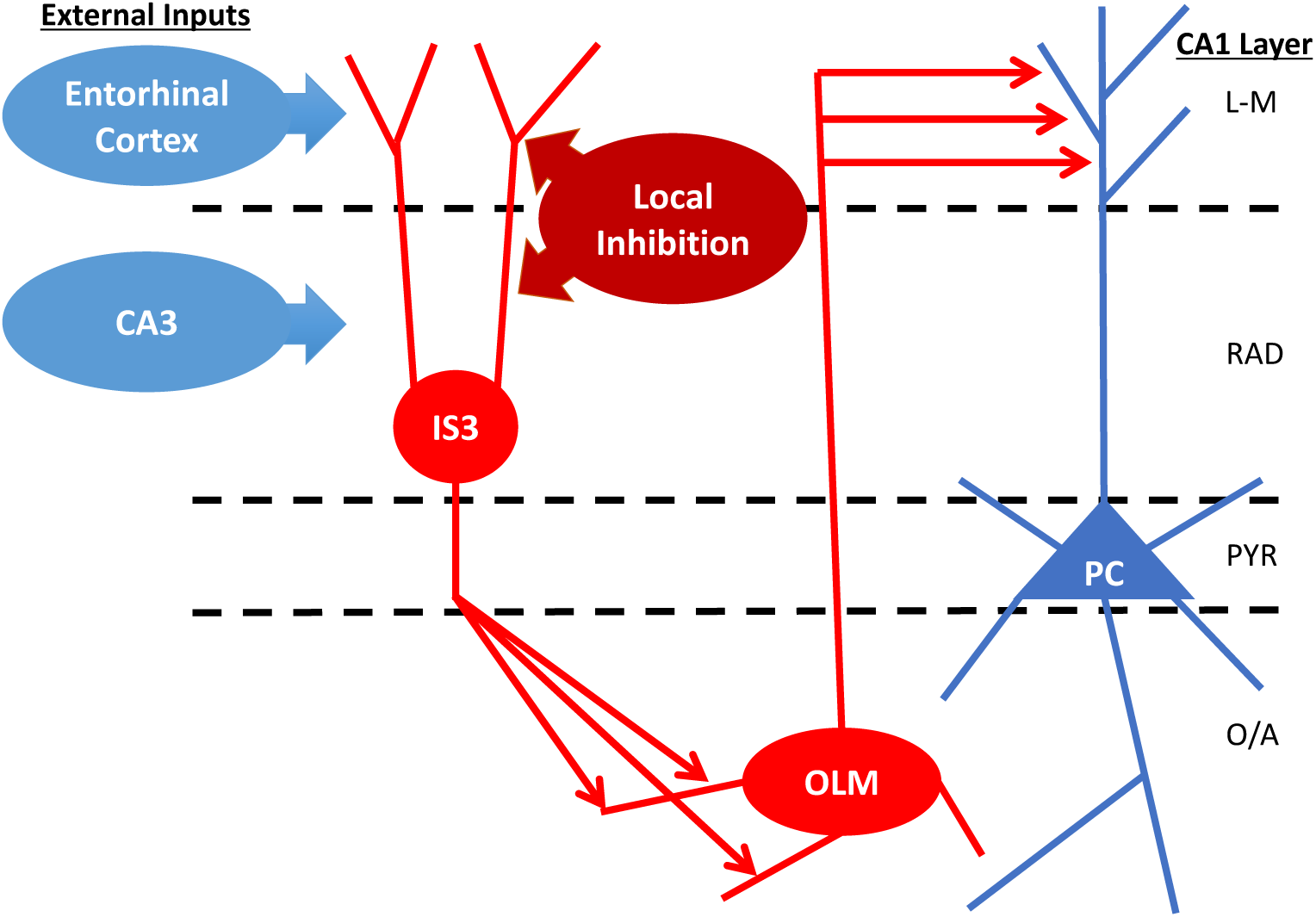
Schematic of layer-specific inputs to IS3 cells in hippocampal CA1. IS3 cell have dendrites that extend into RAD and L-M and receive excitatory inputs from entorhinal cortex and CA3. They also receive inhibitory input from various local inhibitory cell types. Acronyms: CA3 - Cornu Ammonis area 3; IS3 - Interneuron Specific type 3 cell; OLM - Oriens Lacunosum Moleculare cell; PC - Pyramidal cell; L-M - stratum lacunosum moleculare; RAD - stratum radiatum; PYR - stratum pyramidale; O/A - stratum orians/alveus.

**Figure 1–Figure supplement 2.**
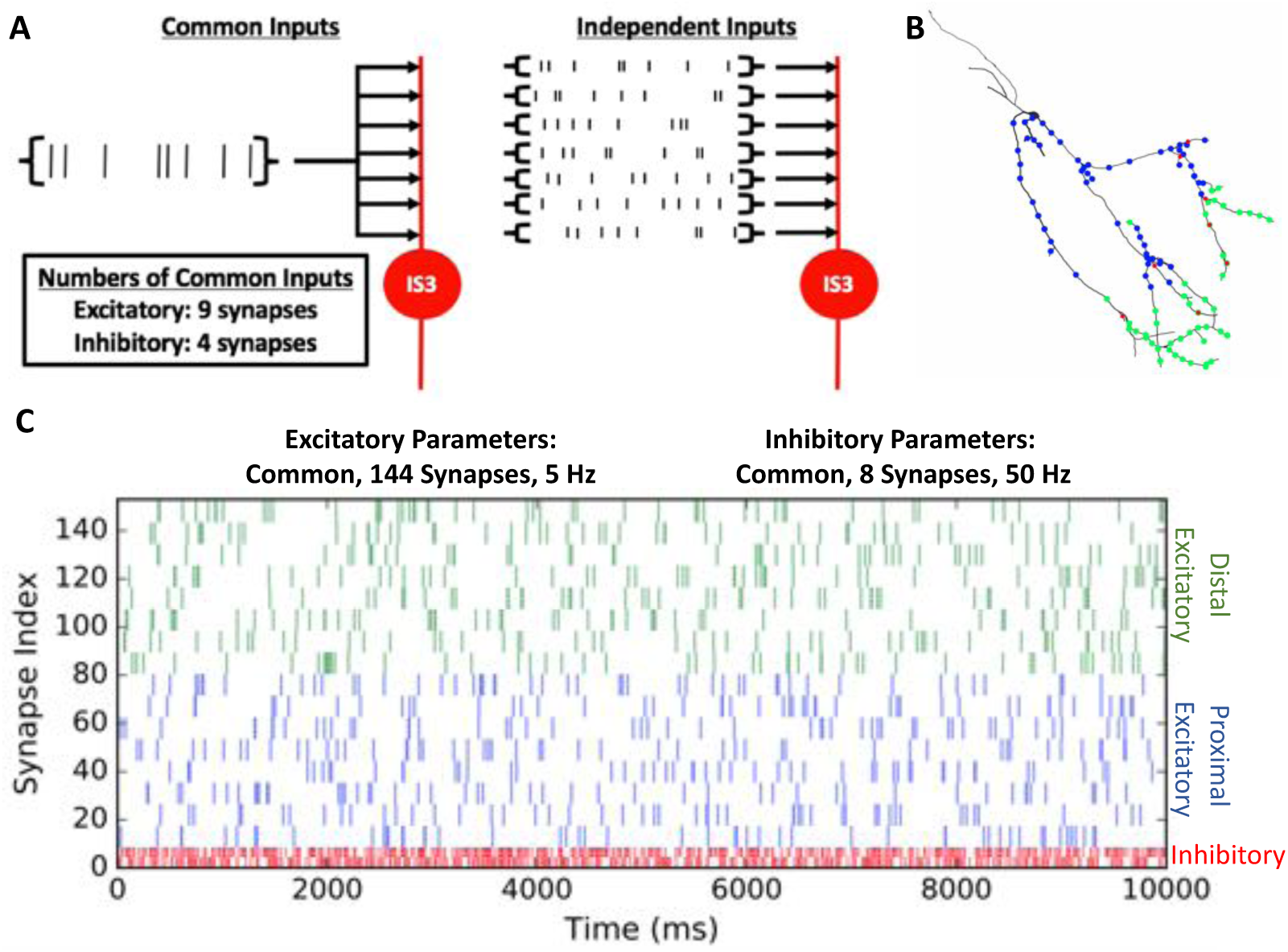
Parameter search setup where we vary the numbers of synapses, the presynaptic spike rates, and whether the inputs are common or independent. **(A)** Schematic of common inputs versus independent inputs. **(B)** Example of randomly chosen synaptic locations along the dendritic compartments of the model, according to the parameters shown in C. Blue are proximal excitatory synapses, green are distal excitatory synapses, and red are inhibitory synapses. Note that synaptic locations can overlap on the same compartments. Apart from separating proximal and distal excitatory synaptic locations, the synaptic locations are chosen randomly. **(C)** Example raster plot of the spike times of synapses using parameter values as shown. Color scheme is the same as in B. Since this plot shows a common input scenario, note that the ‘lines’ in the raster plot are actually a series of dots (9 dots for excitatory and 4 dots for inhibitory) representing groups of synapses receiving the same (i.e., common) presynaptic spike trains.

**Figure 3–Figure supplement 1.**
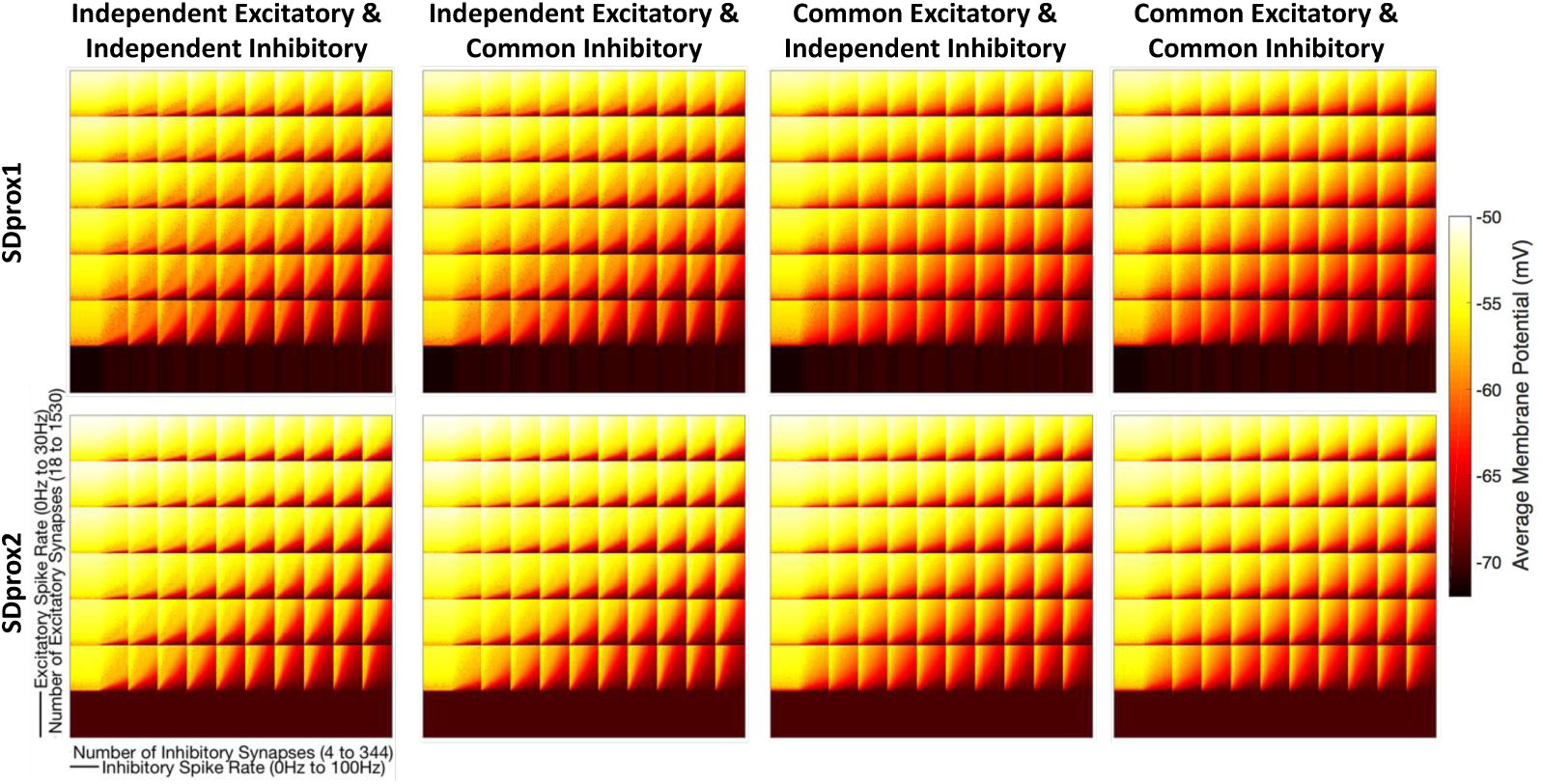
CBDR plots showing the mean subthreshold membrane potential 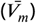. Note that the threshold 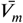 in the HC metric equation is -66.7 mV.

**Figure 3–Figure supplement 2.**
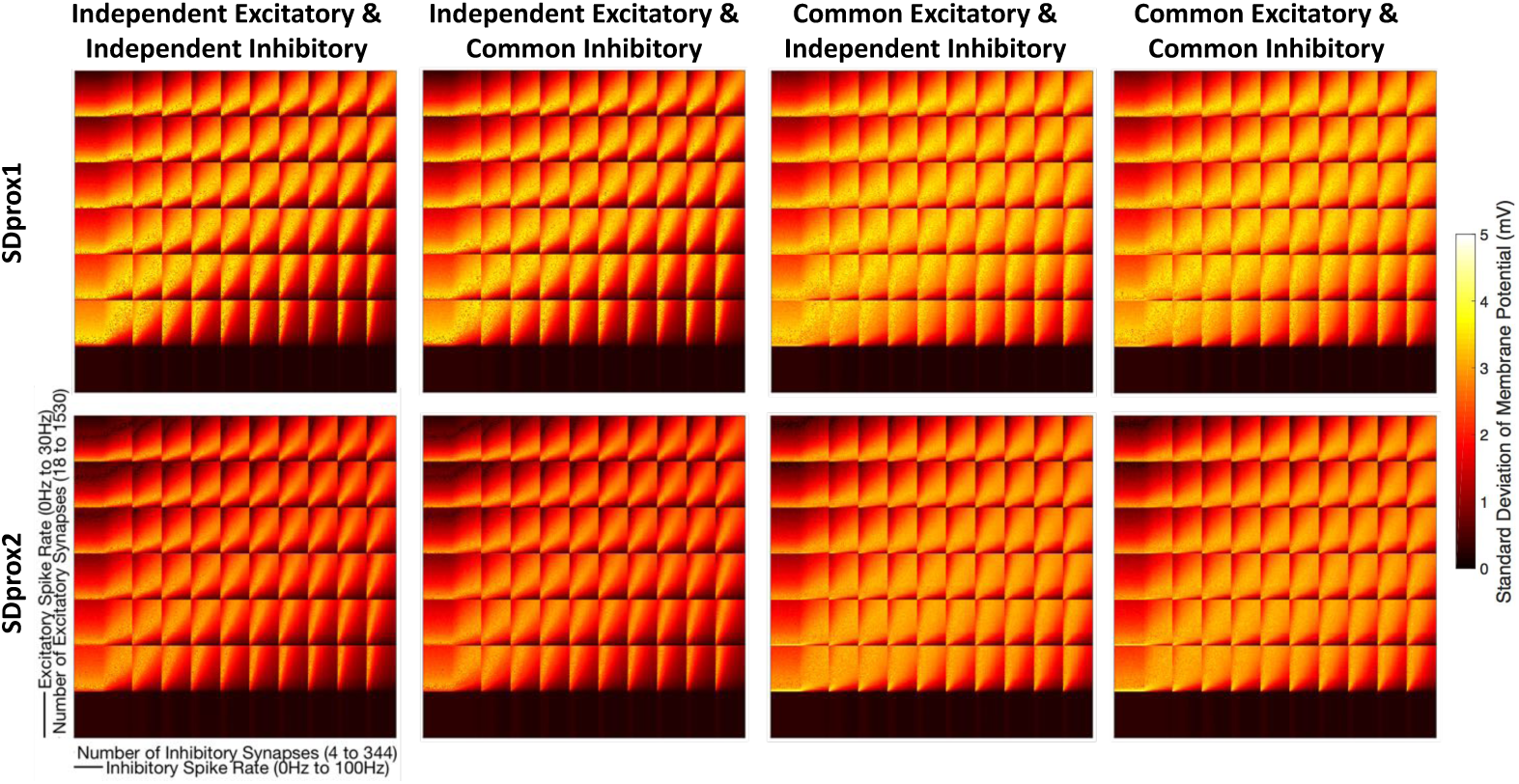
CBDR plots showing the subthreshold membrane potential standard deviation 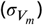. Note that the threshold 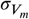 in the HC metric equation is 2.2 mV.

**Figure 3–Figure supplement 3.**
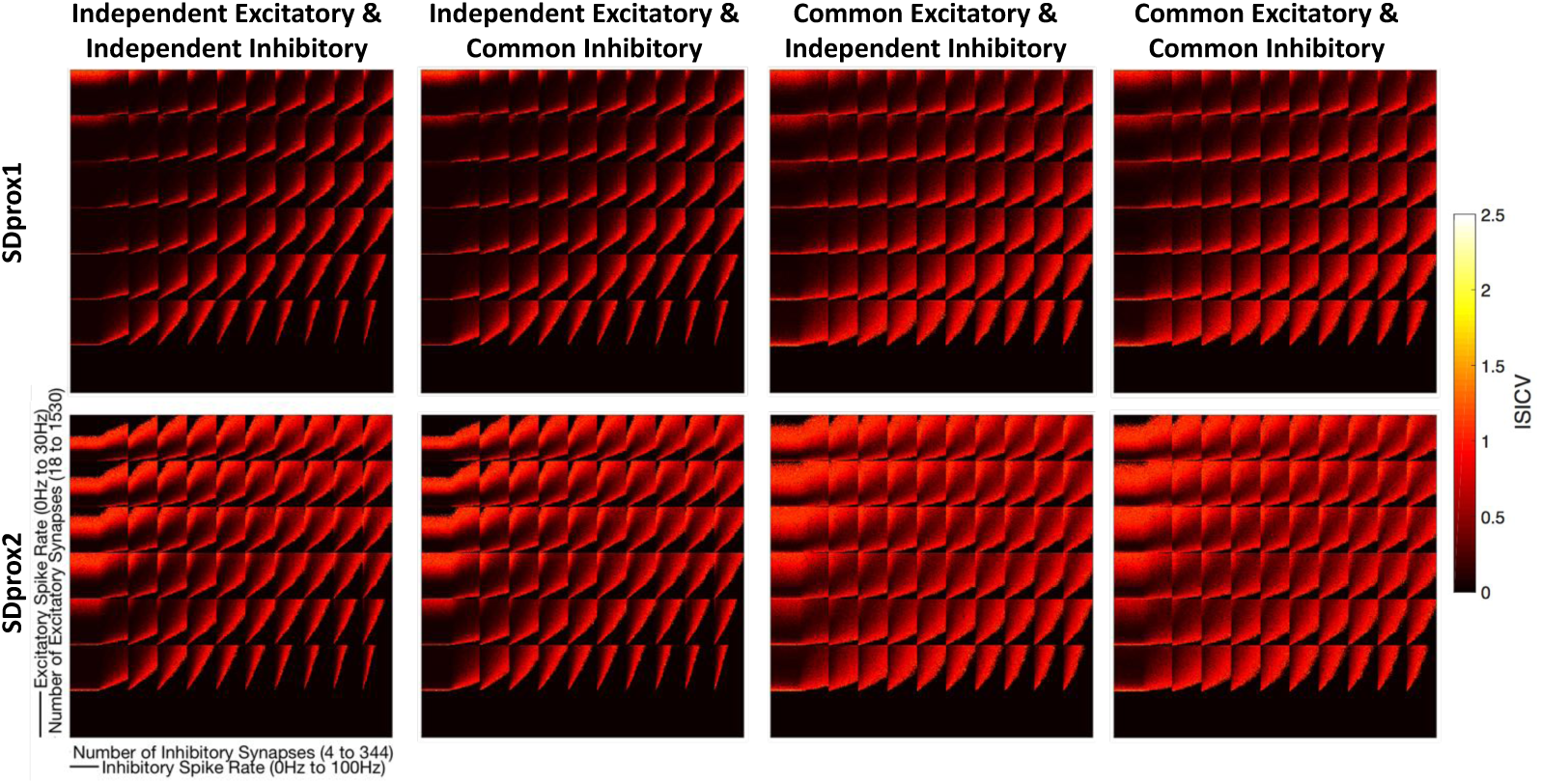
CBDR plots showing the interspike interval coefficient of variation (*ISICV*). Note that the threshold *ISICV* in the HC metric equation is 0.8.

**Figure 3–Figure supplement 4.**
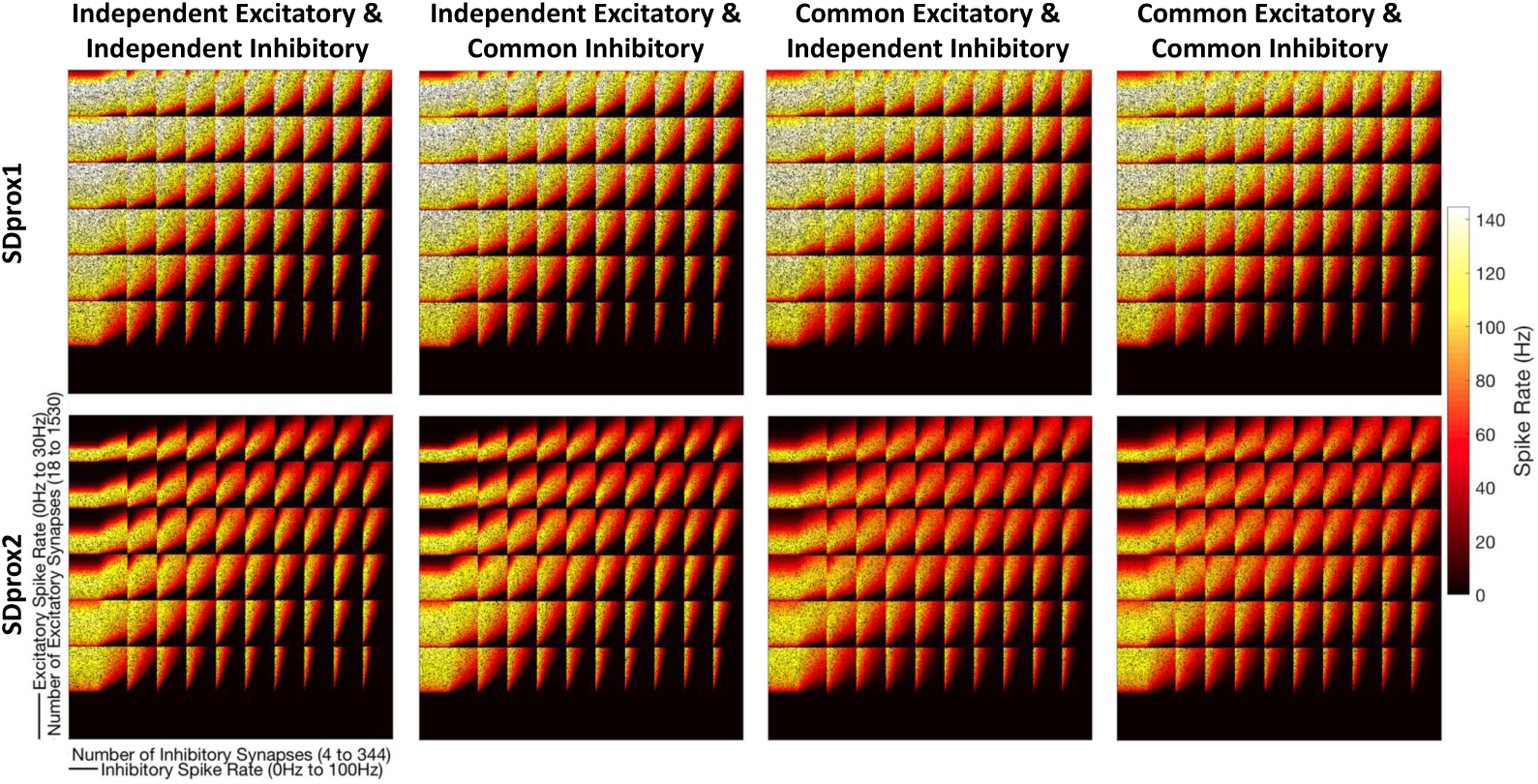
CBDR plots showing the spike rates produced by the IS3 cells in each simulation.

**Figure 3–Figure supplement 5.**
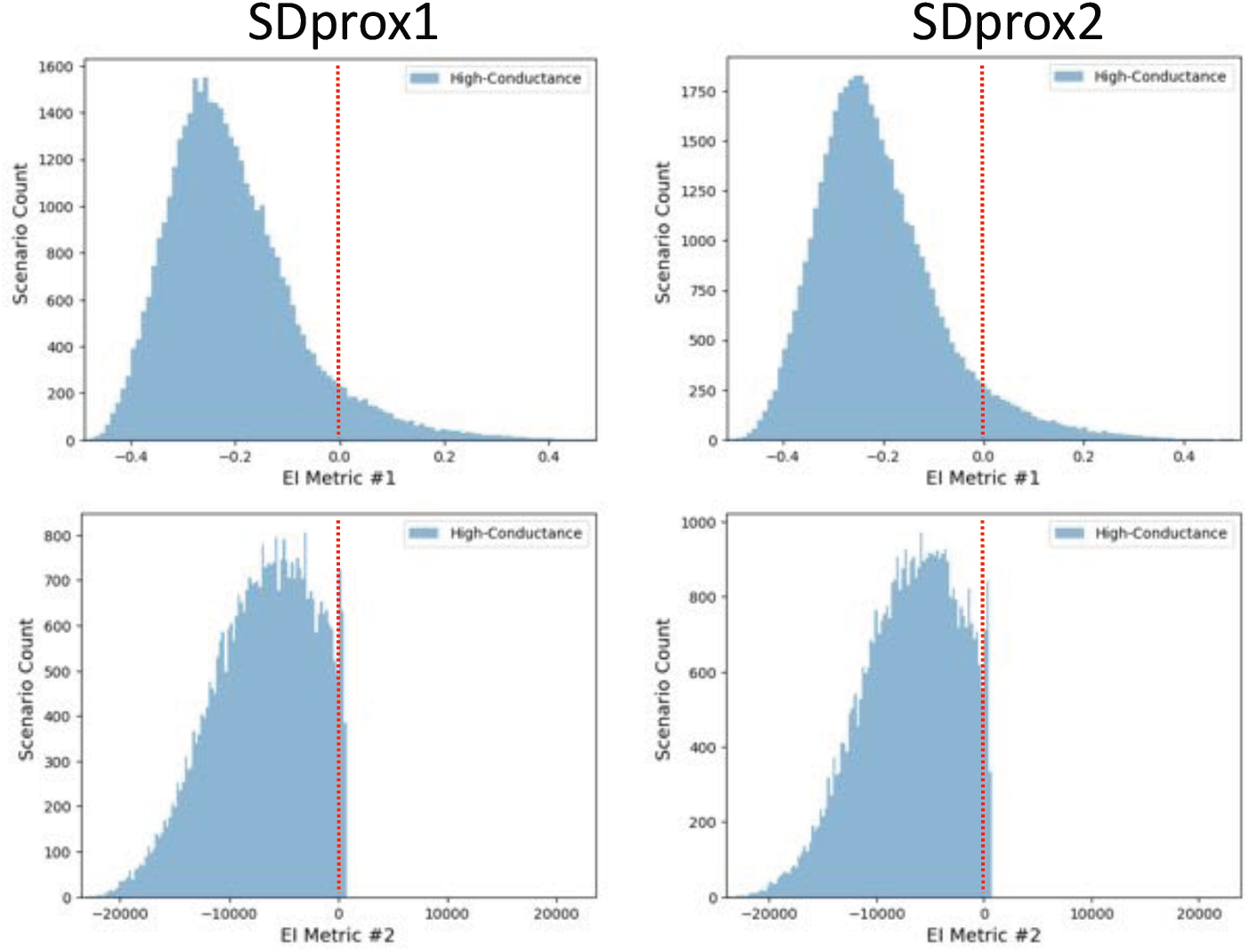
Histogram distributions of EI Metric #1 (***Equation 6***) and #2 (***Equation 7***) for HC states of the SDprox1 and SDprox2 models. Red dashed lines indicate EI metric values of zero where excitation and inhibition are approximately even.

**Figure 3–Figure supplement 6.**
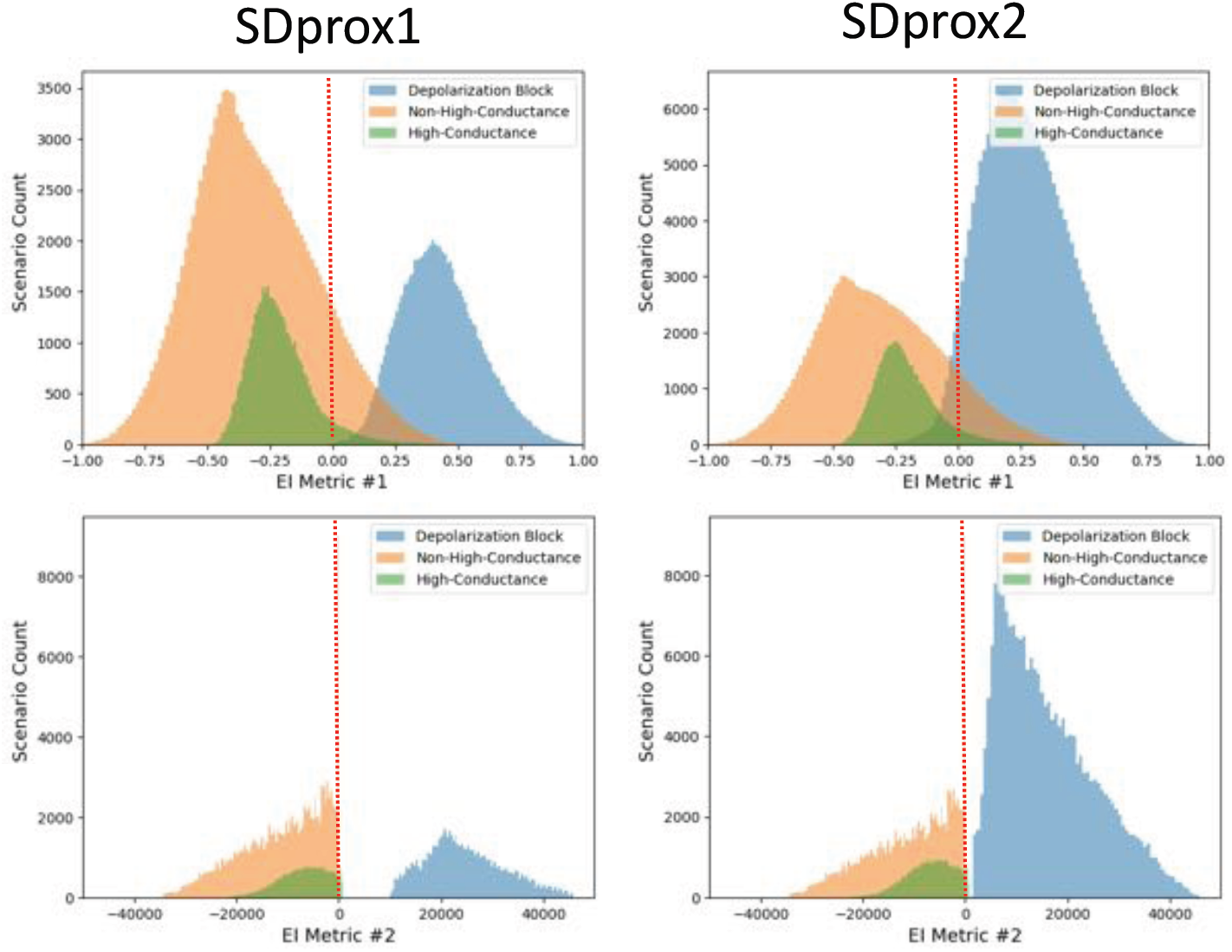
Histogram distributions of EI metrics for each of the defined states (i.e. HC, NHC, and DB) on the same plot, and for SDprox1 and SDprox2 models. Red dashed lines indicate EI metric values of zero (for both metrics) where excitation and inhibition are approximately even.

**Figure 4–Figure supplement 1.**
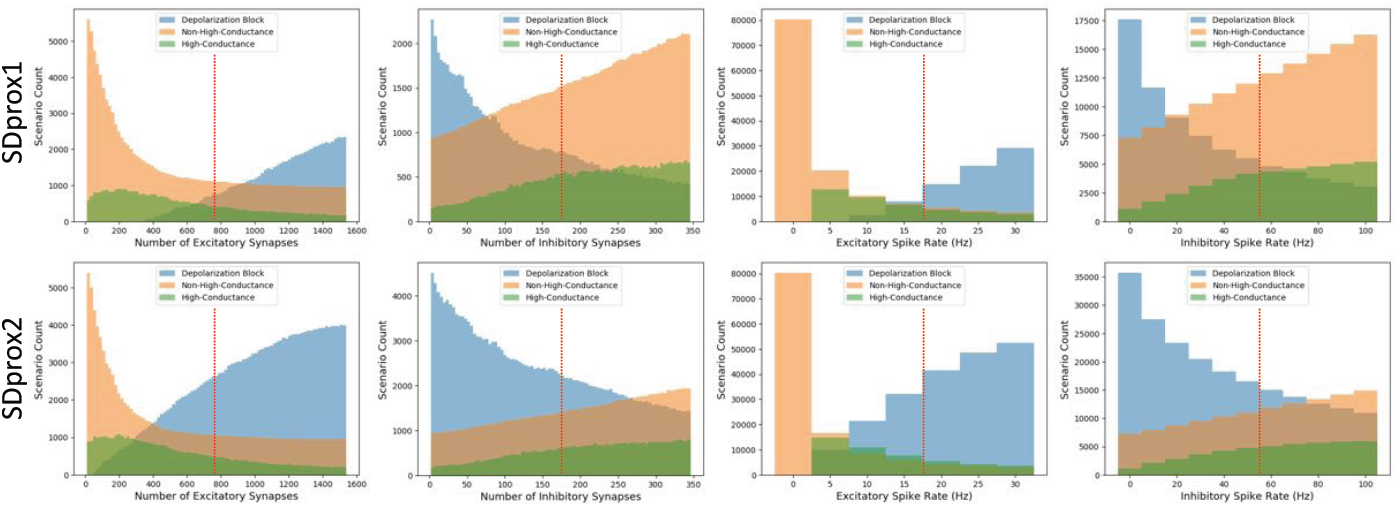
Histogram distributions of input parameters for each of the defined states (i.e. HC, NHC, and DB) for SDprox1 and SDprox2 models. Red lines delineate the ranges of parameters when splitting into 16 pools (see ***Table 1***).

**Figure 4–Figure supplement 2.**
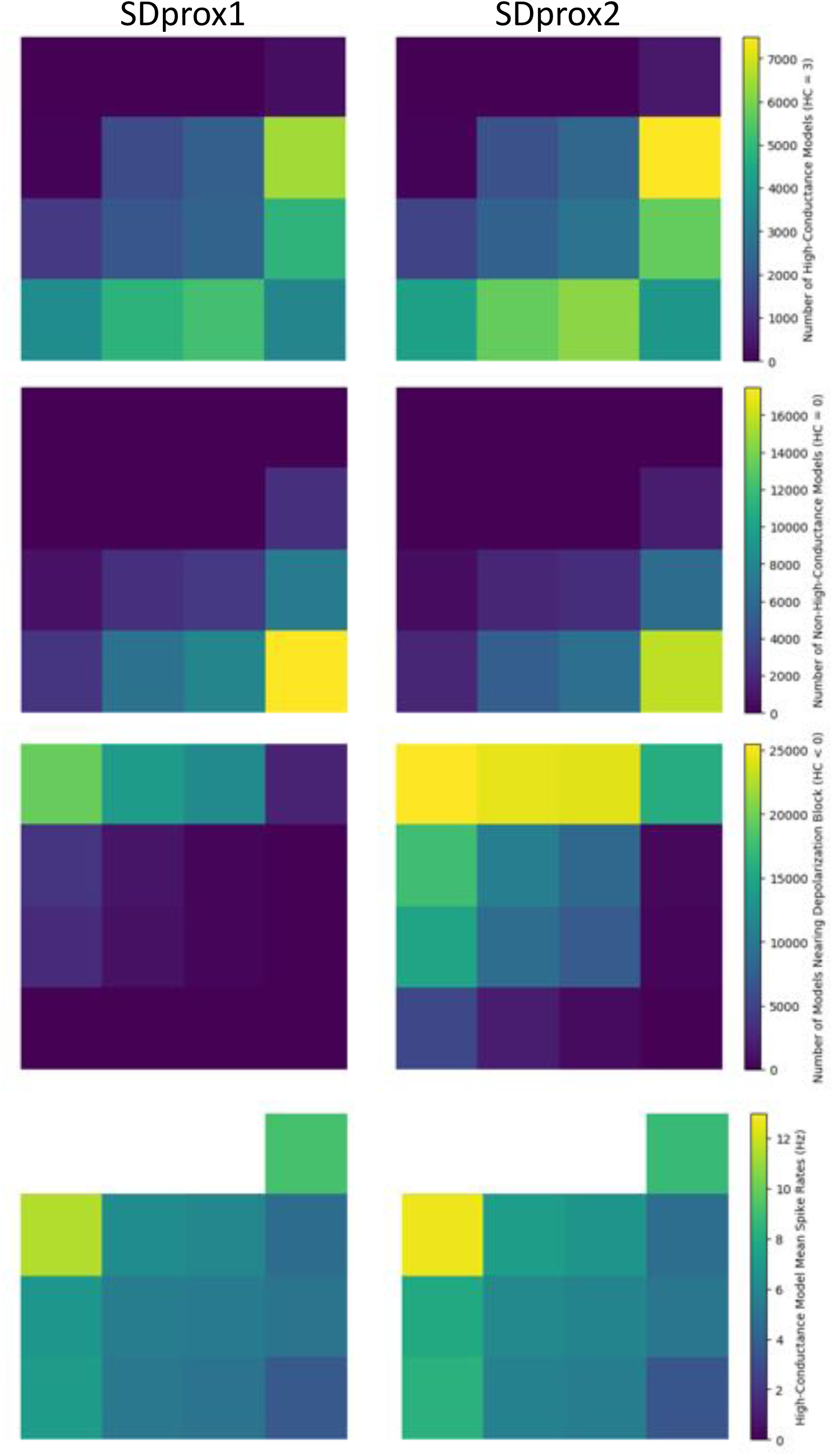
Top three rows of plots: Numbers of scenarios for each defined state (row 1: HC; row 2: NHC; row 3: DB). Bottom row shows the average spike rates of the IS3 cell models in a HC state. Each colored square in each plot represents one of the 16 pools organized as described in the layout of ***Figure 4***.

**Figure 5–Figure supplement 1.**
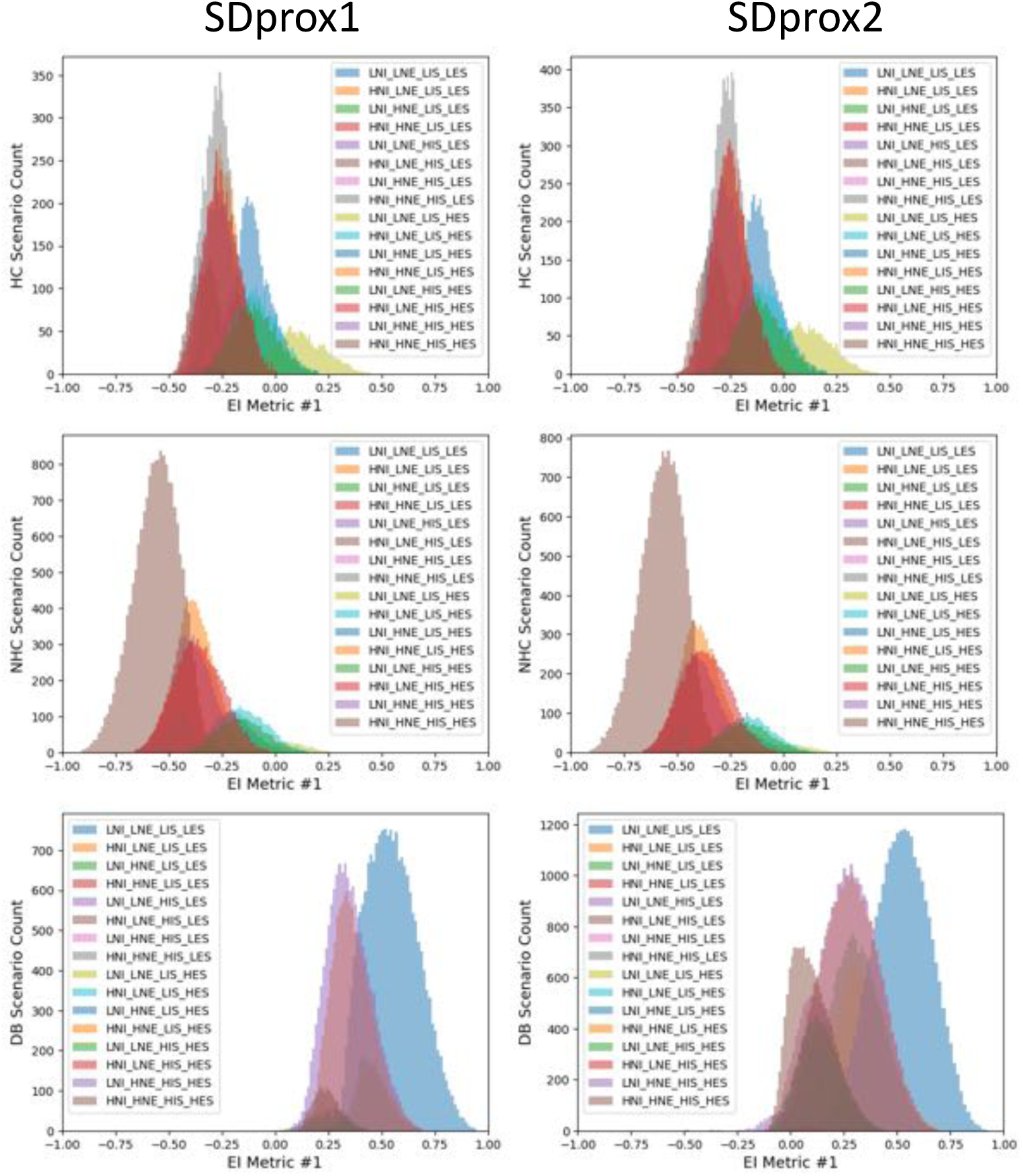
Distributions of EI metric 1 values for each pool for both SDprox1 and SDprox2 models. Going from top to bottom: HC pool distributions, NHC pool distributions, DB pool distributions. Consult ***Figure 4*** for description of pool labels shown in the legend.

**Figure 5–Figure supplement 2.**
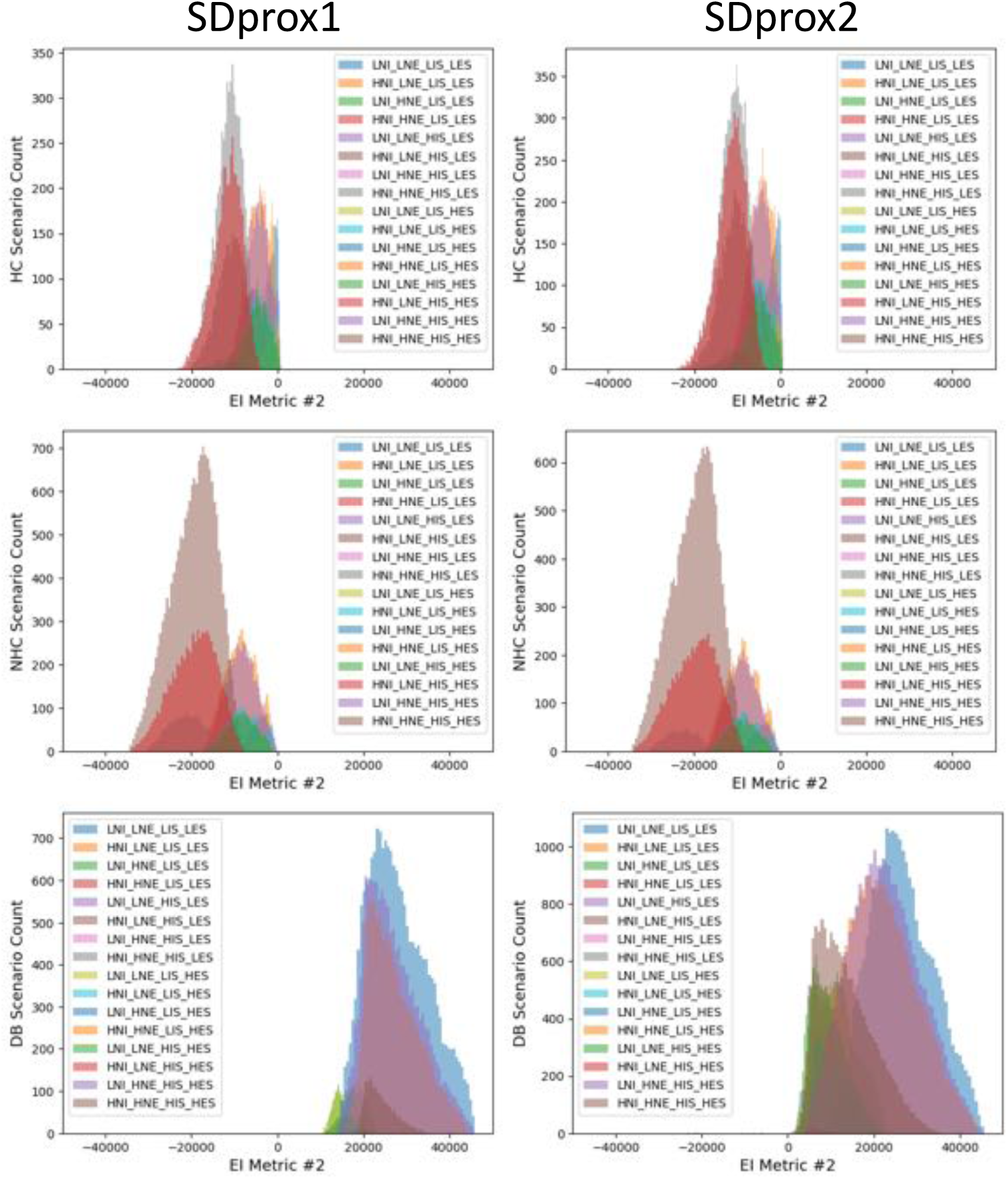
Distributions of EI metric 2 values for each pool for both SDprox1 and SDprox2 models. Going from top to bottom: HC pool distributions, NHC pool distributions, DB pool distributions. Consult ***Figure 4*** for description of pool labels shown in the legend.

**Figure 6–Figure supplement 1.**
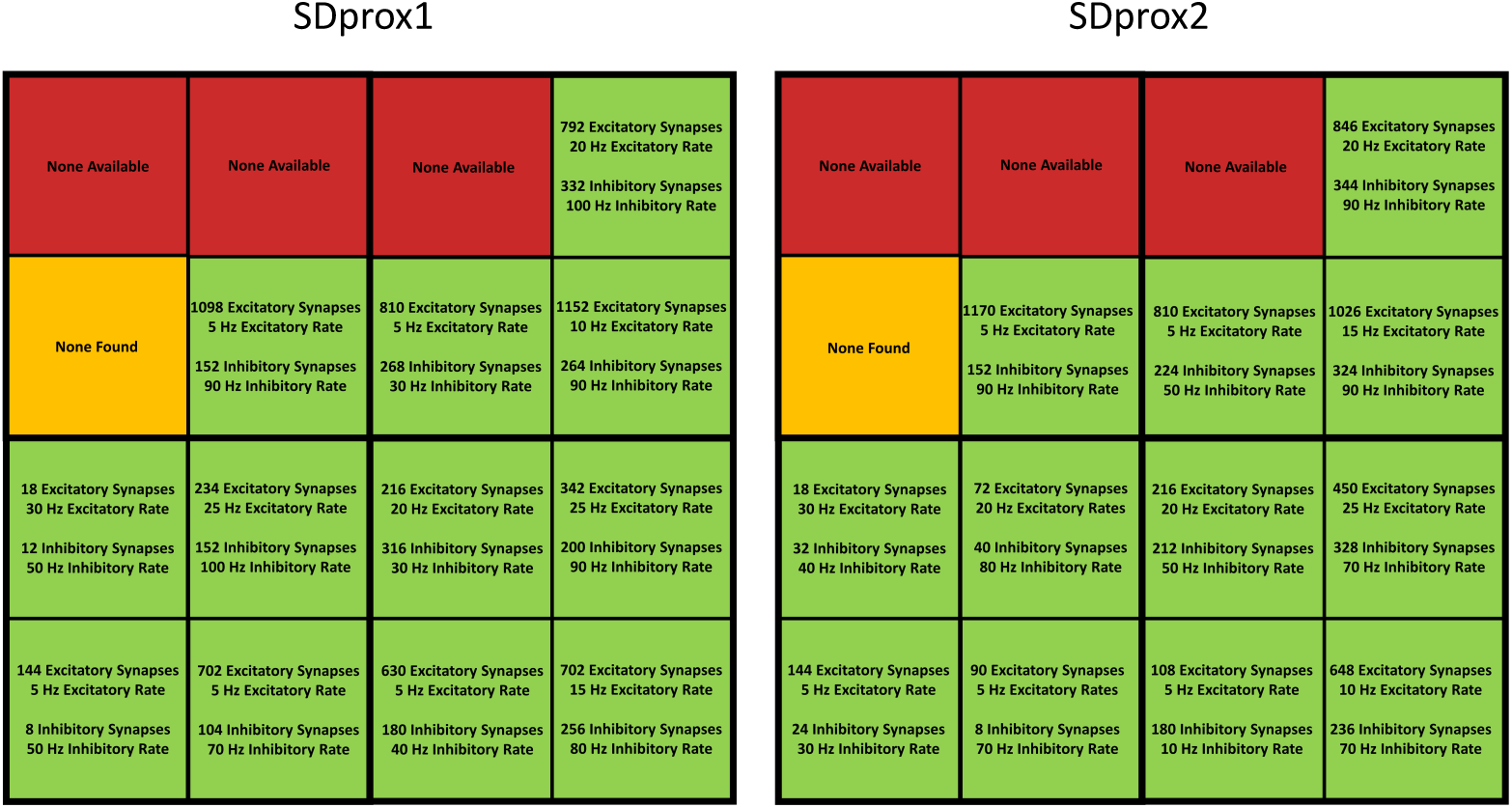
Representative HC scenario input parameter values for each pool.

**Figure 6–Figure supplement 2.**
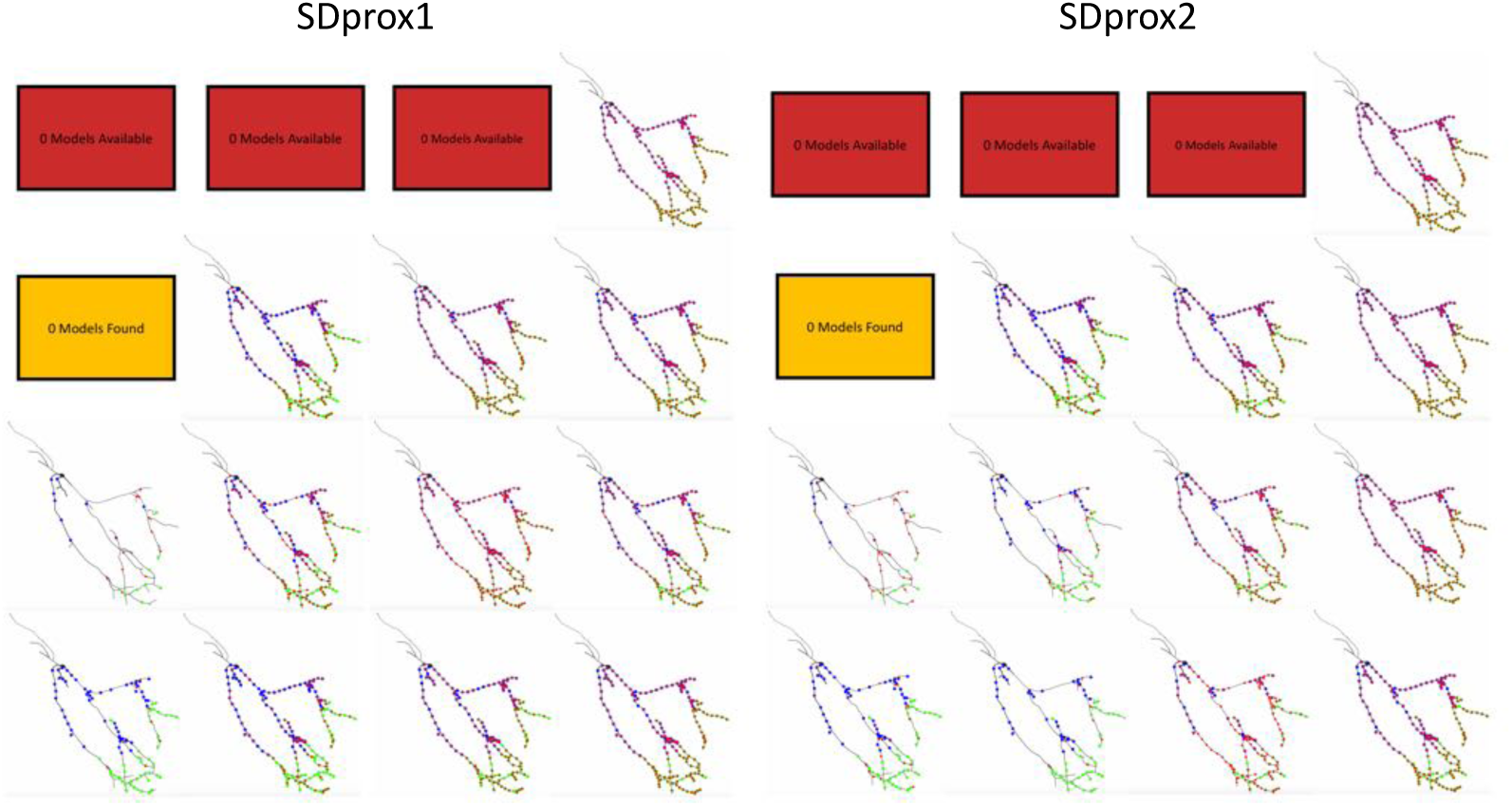
Representative HC scenario locations and numbers of active inputs for each pool. Proximal excitatory inputs are shown as blue dots, distal excitatory inputs as green dots, and inhibitory inputs as red dots (i.e. same as in ***Figure 1, Figure Supplement 2***).

**Figure 6–Figure supplement 3.**
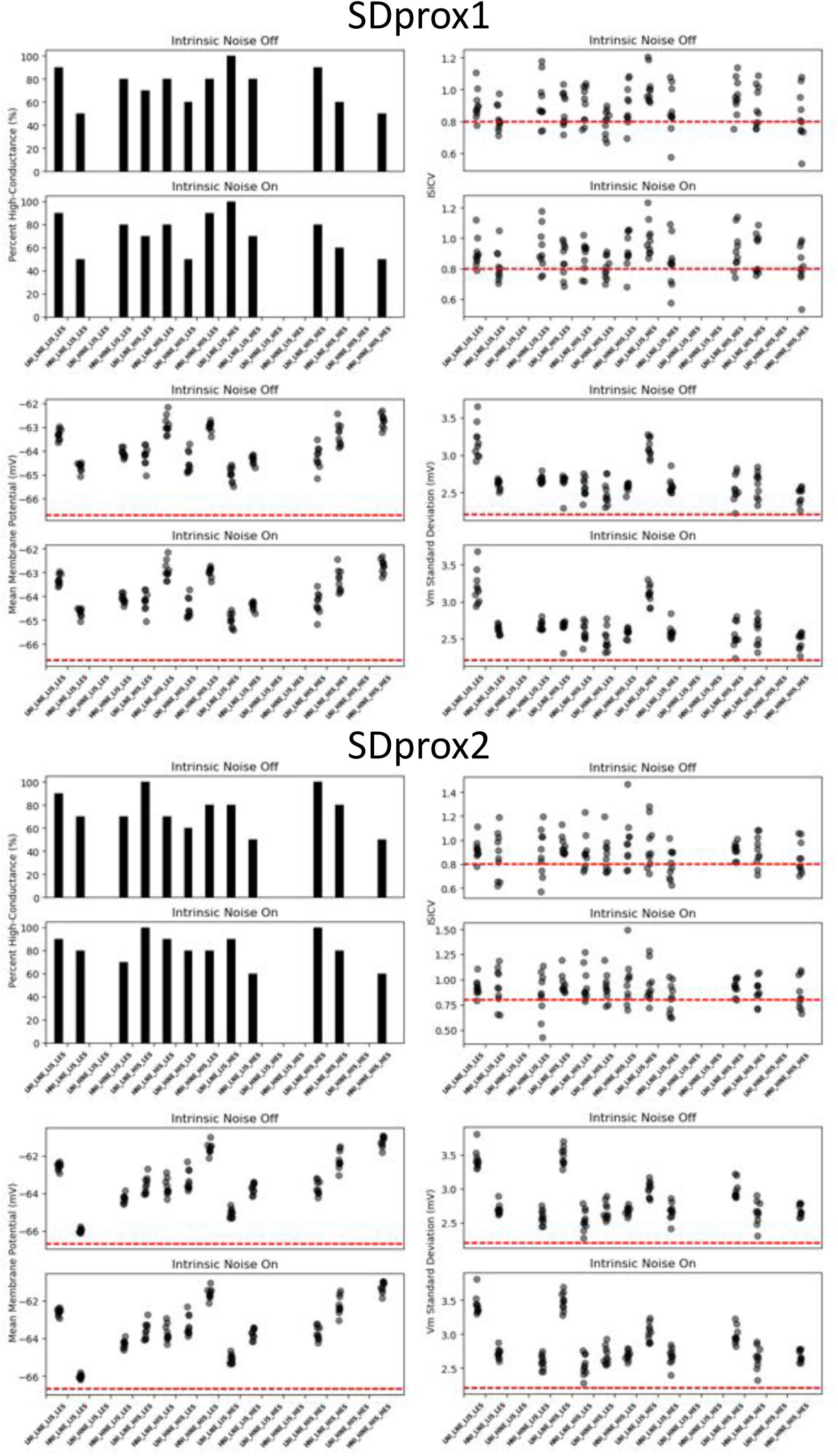
HC consistency of the representative HC scenarios with and without intrinsic noise. Top left panels: % of simulations that each HC representative scenario remained as a HC state when re-run 10 times with 10 different seeds. Other panels: *ISICV*, mean *V*_*m*_, and standard deviation of the *V*_*m*_′ for the 10 simulations of each representative scenario. Red lines indicate the threshold values used in the HC metric. Consult ***Figure 4*** for pool labels.

**Figure 6–Figure supplement 4.**
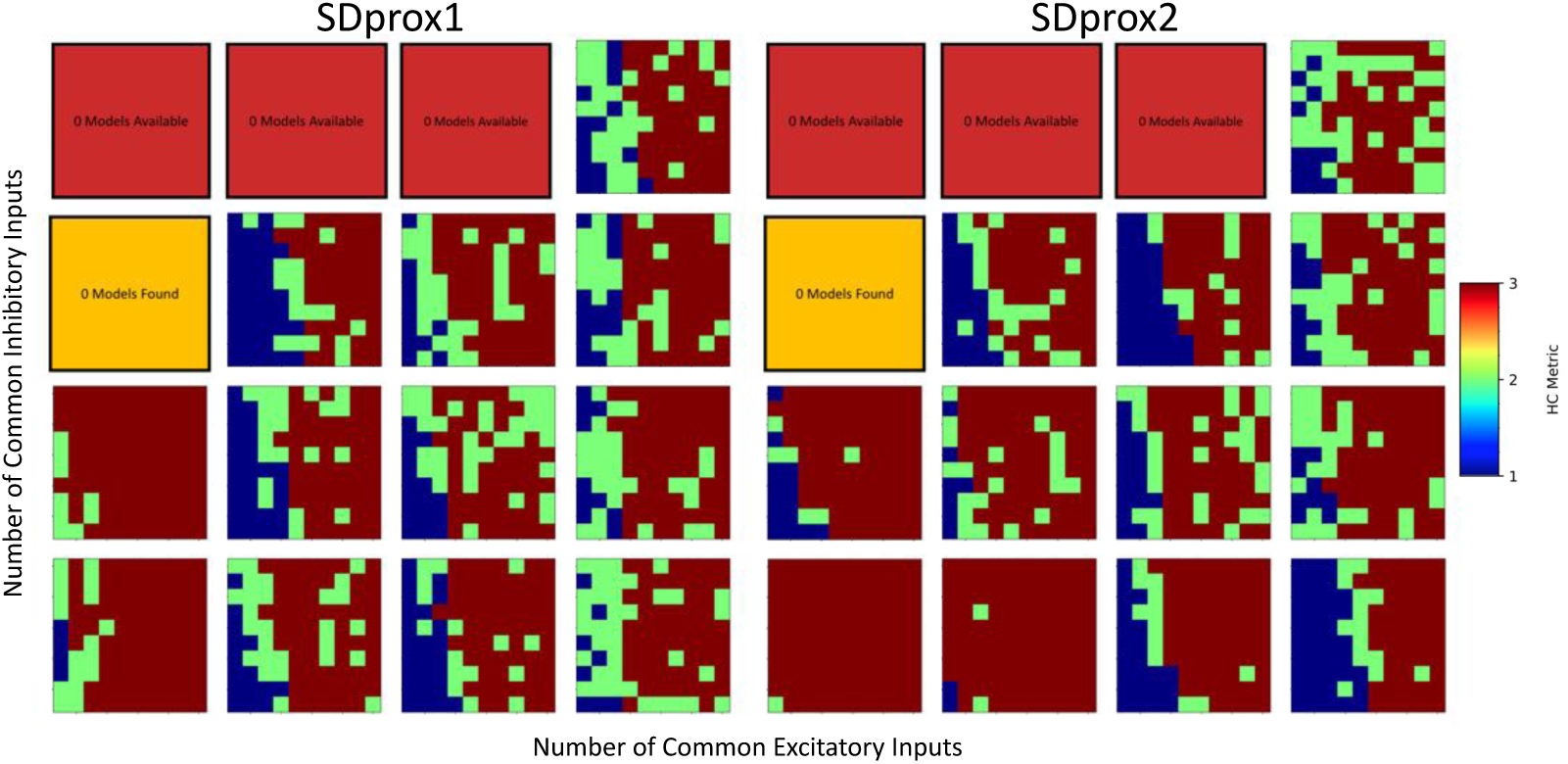
The influence of the number of common inputs on the HC metric for each representative HC scenario from the given pool (see ***Figure 6***). Note that excitatory common inputs are plotted on the x-axis and inhibitory common inputs are plotted on the y-axis. (x-axis and y-axis ranges: 1 to 10 common inputs).

**Figure 6–Figure supplement 5.**
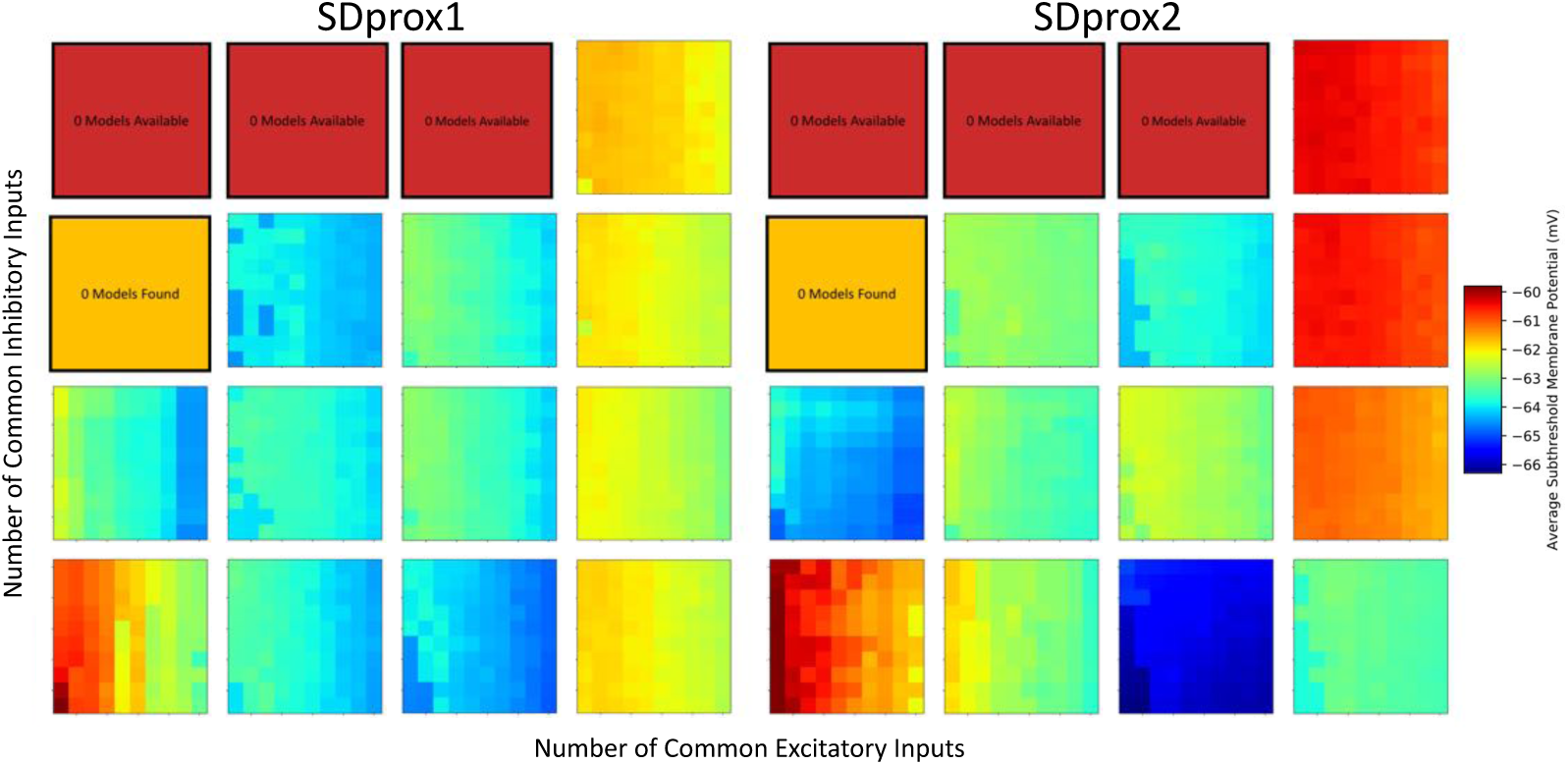
The influence of the number of common inputs on the mean subthreshold membrane potential (part of the HC metric) for each representative HC scenario from the given pool (same format as in ***Figure 6,Figure Supplement 4***).

**Figure 6–Figure supplement 6.**
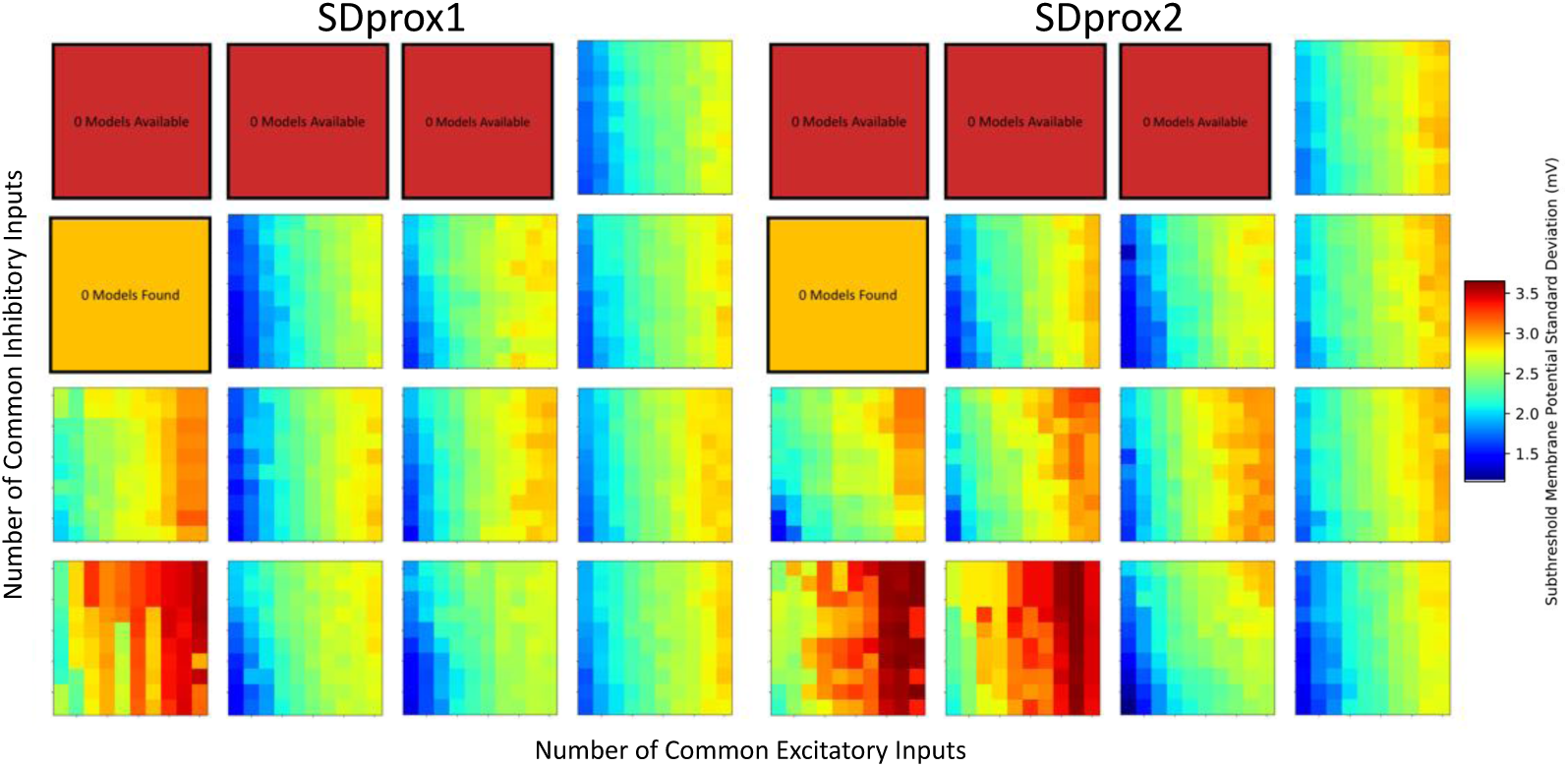
The influence of the number of common inputs on the subthreshold membrane potential standard deviation (part of the HC metric) for each representative HC scenario from the given pool (same format as in ***Figure 6,Figure Supplement 4***).

**Figure 6–Figure supplement 7.**
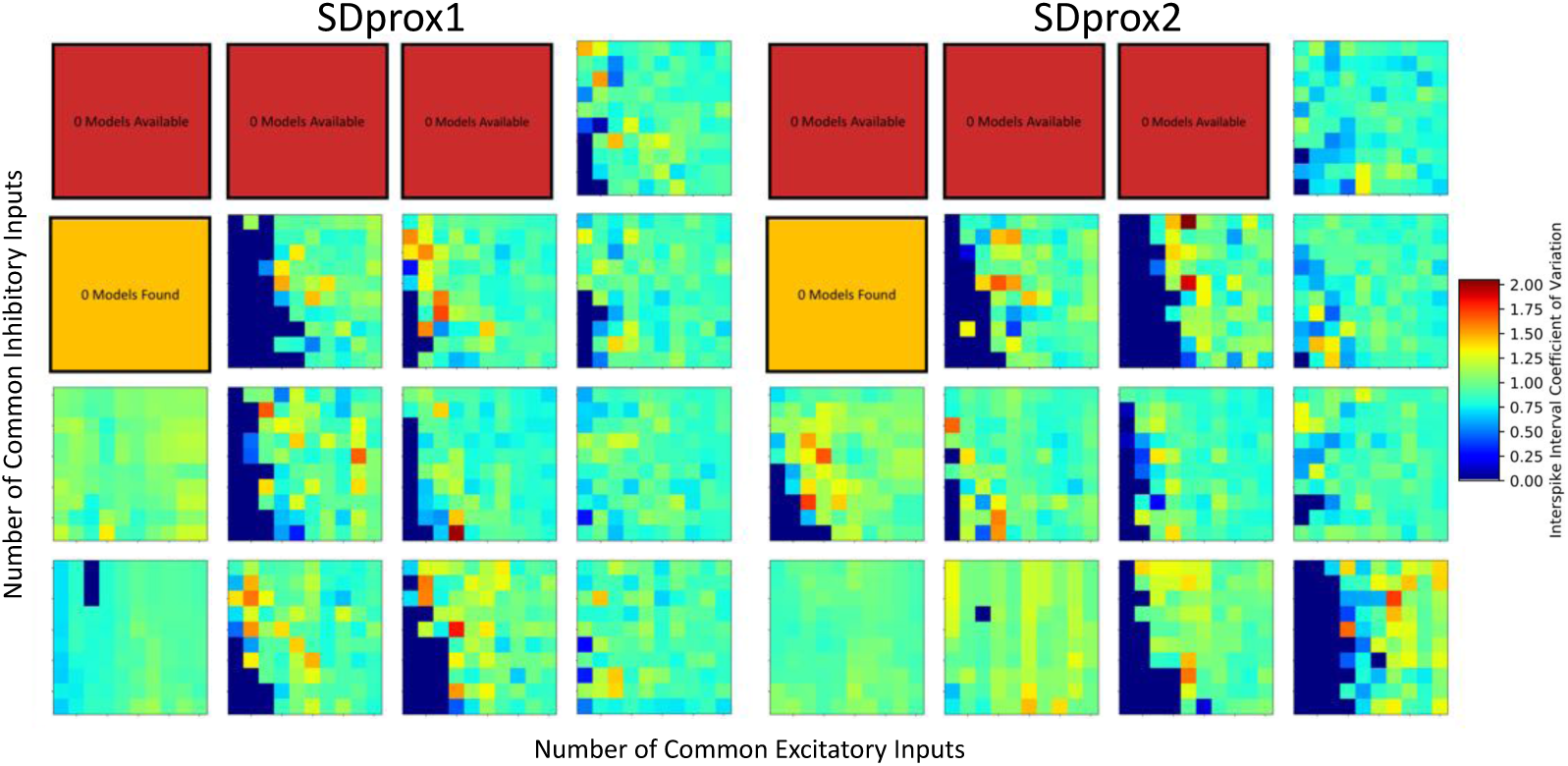
The influence of the number of common inputs on the interspike interval coefficient of variation (part of the HC metric) for each representative HC scenario from the given pool (same format as in ***Figure 6,Figure Supplement 4***).

**Figure 6–Figure supplement 8.**
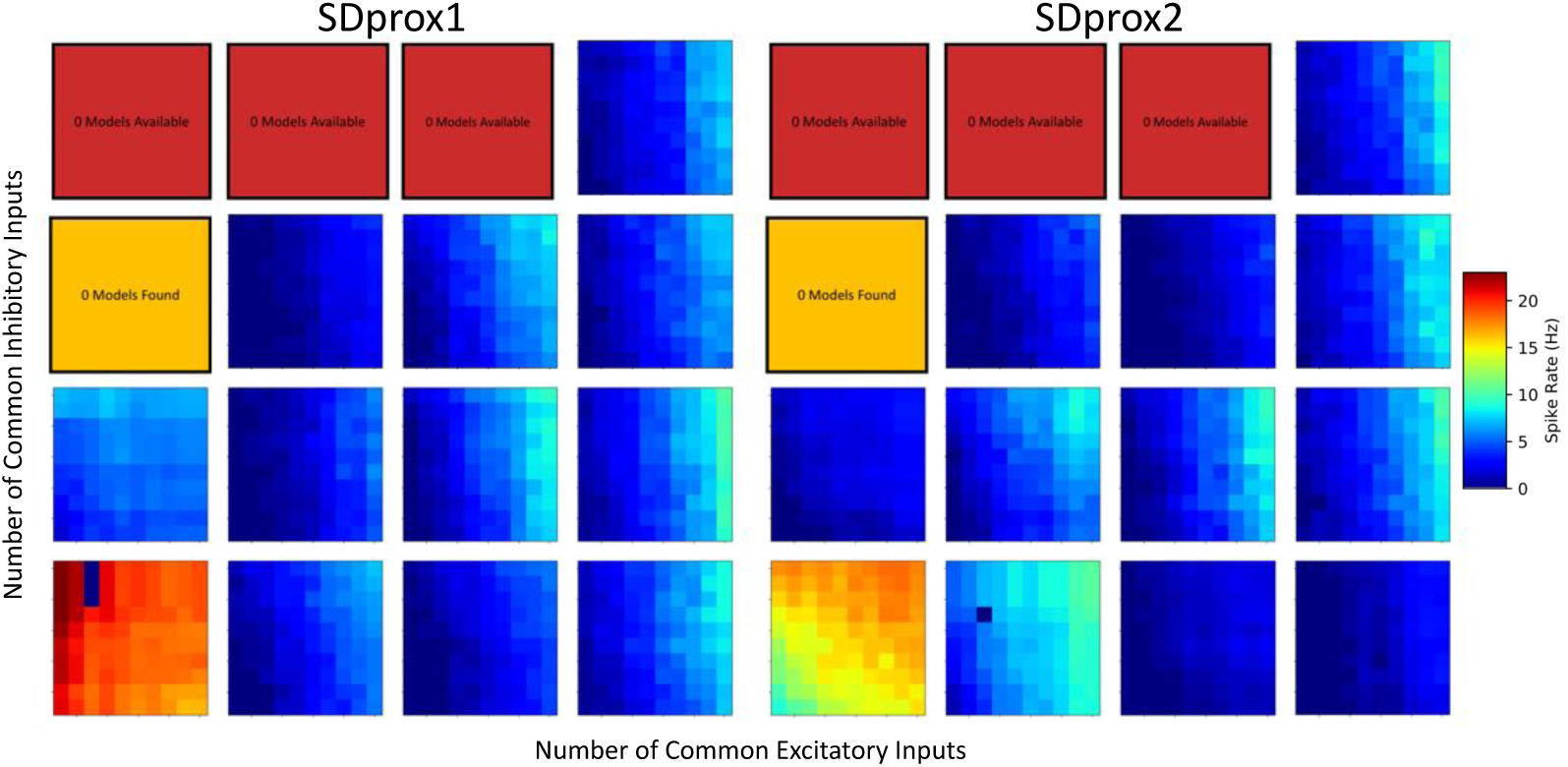
The influence of the number of common inputs on the resulting spike rate of the IS3 cell model for each representative HC scenario from the given pool (same format as in ***Figure 6, Figure Supplement 4***).

**Figure 6–Figure supplement 9.**
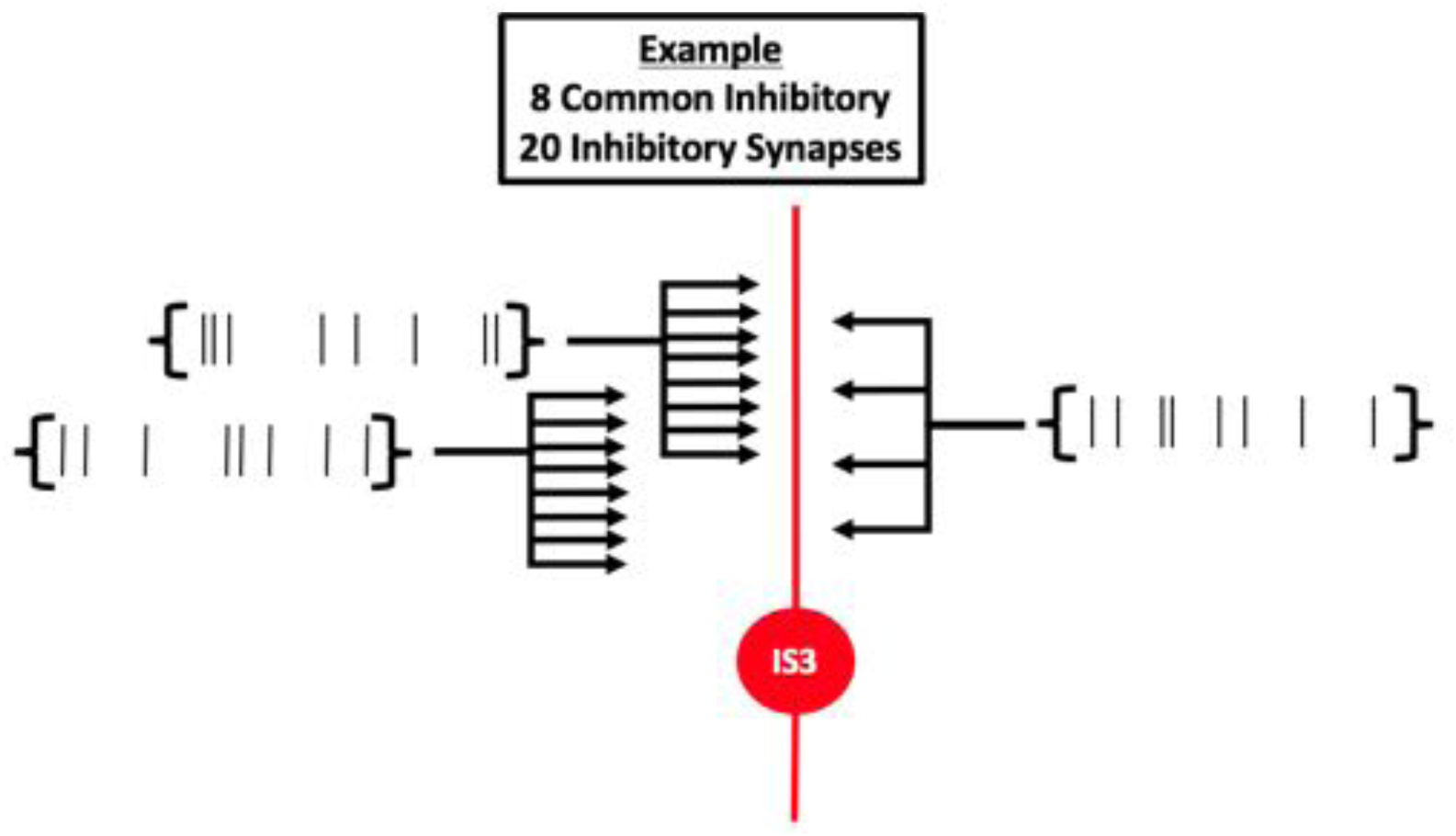
Schematic demonstrating an example where there are 20 inhibitory inputs available, but the number of common inhibitory inputs is set to 8. This will create two groups of 8 inhibitory synapses receiving common inputs, and then a remainder group of 4 inhibitory synapses receiving common inputs. Thus our parameter exploration is inherently inexact.

**Figure 8–Figure supplement 1.**
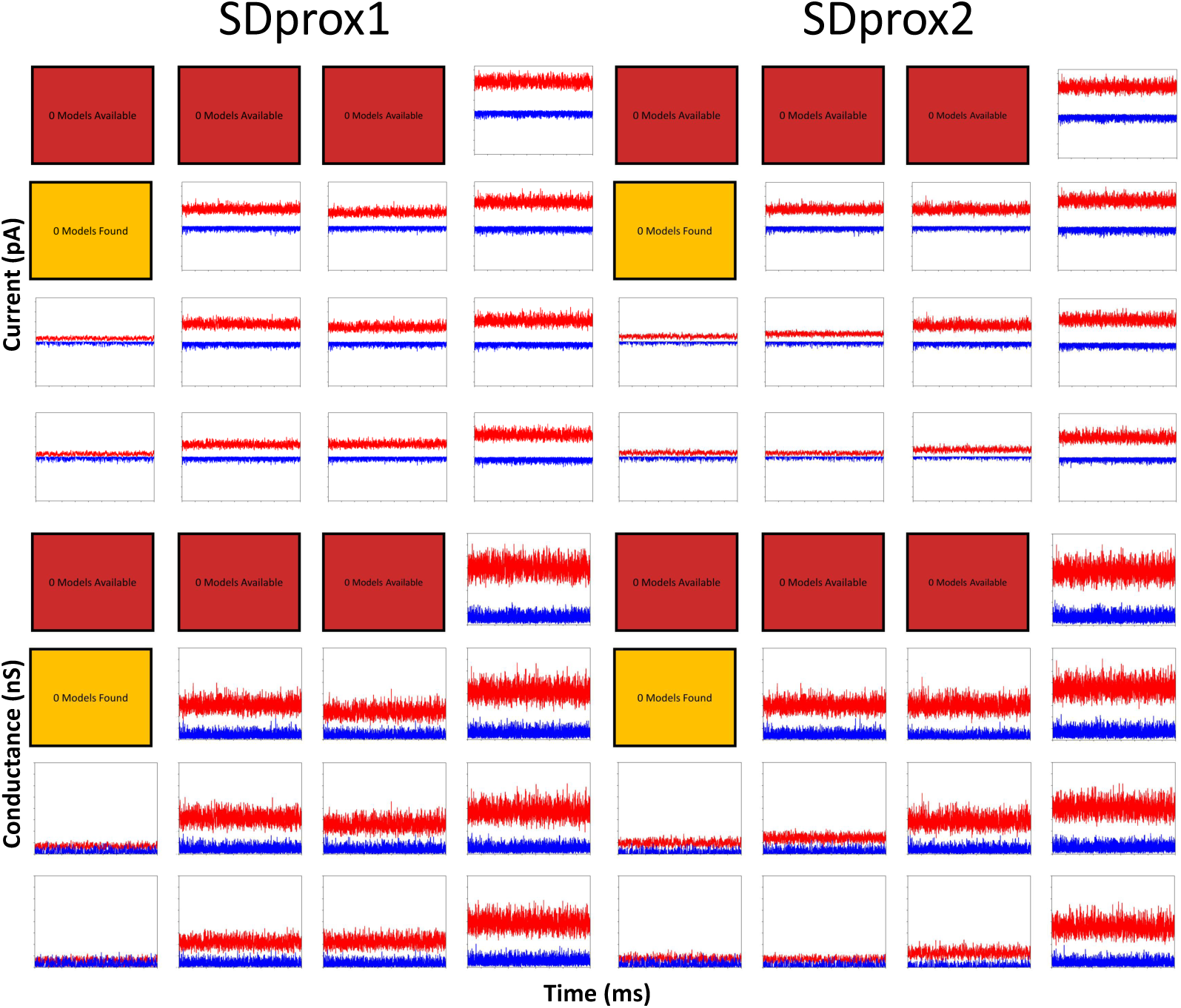
Inhibitory or excitatory currents (y-axes: -1100 pA to 1100 pA) and conductances (y-axes: 0 nS to 16 nS) recorded when excitatory or inhibitory currents are not removed completely (i.e. the only measure taken is by changing the holding potential to their respective reversal potentials; x-axes: 1000 ms to 10,000 ms). Top panels: excitatory (blue) and inhibitory (red) currents recorded in representative HC scenarios of the SDprox1 and SDprox2 models. Bottom panels: excitatory (blue) and inhibitory (red) conductances recorded in representative HC scenarios of the SDprox1 and SDprox2 models.

**Figure 10–Figure supplement 1.**
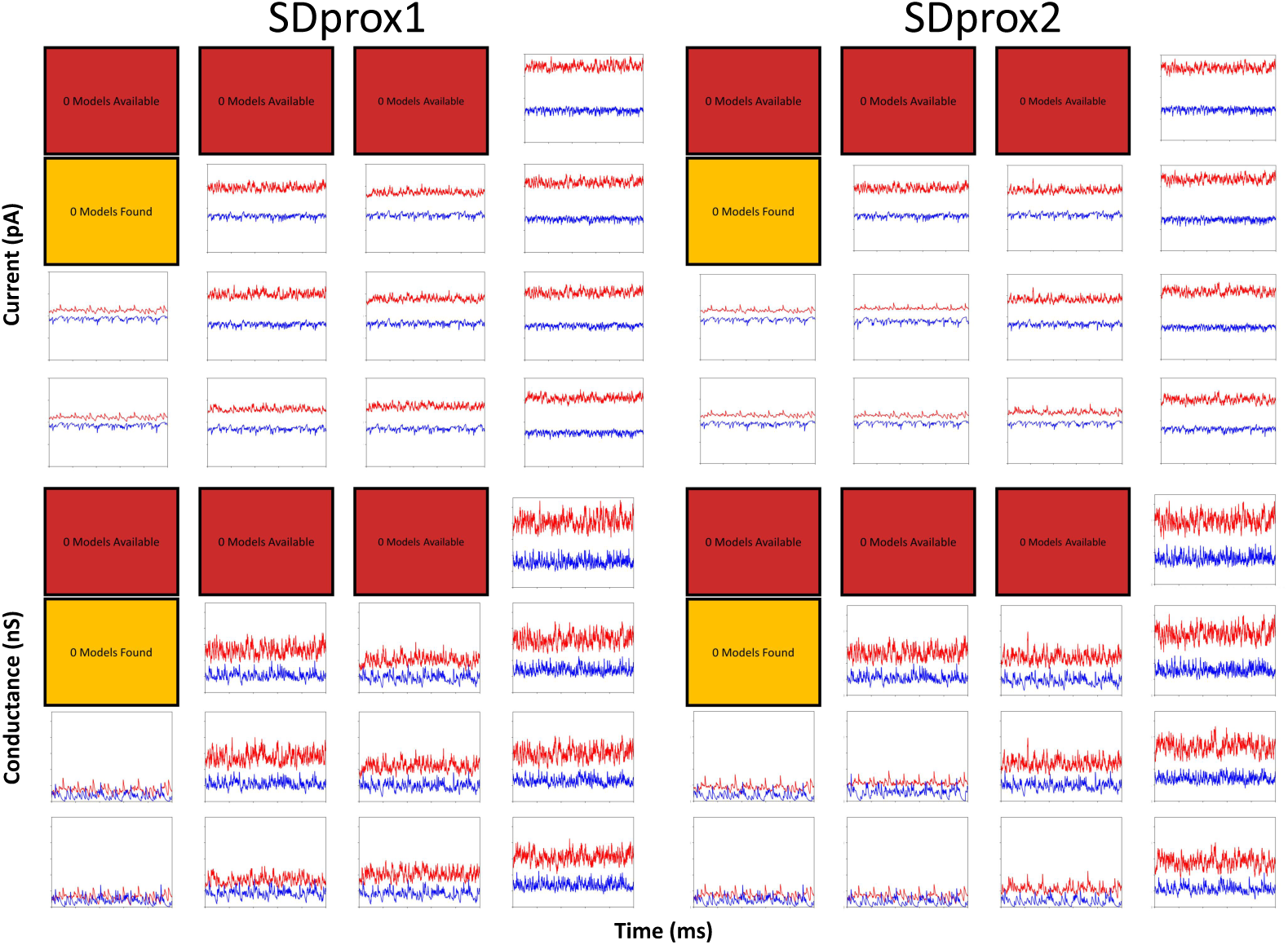
Inhibitory or excitatory currents (y-axes: -1000 pA to +1000 pA) and conductances (y-axes: 0 nS to 14 nS) recorded during HC input scenarios + theta-timed inputs (same protocol as in ***Figure 8***; x-axes: 9000 ms to 10,000 ms). Top panels: excitatory (blue) and inhibitory (red) currents recorded in representative HC scenarios of the SDprox1 and SDprox2 models. Bottom panels: excitatory (blue) and inhibitory (red) conductances recorded in representative HC scenarios of the SDprox1 and SDprox2 models. Note that we zoomed in to the last second of the simulation in order to better visualize the time-scale of theta

**Figure 10–Figure supplement 2.**
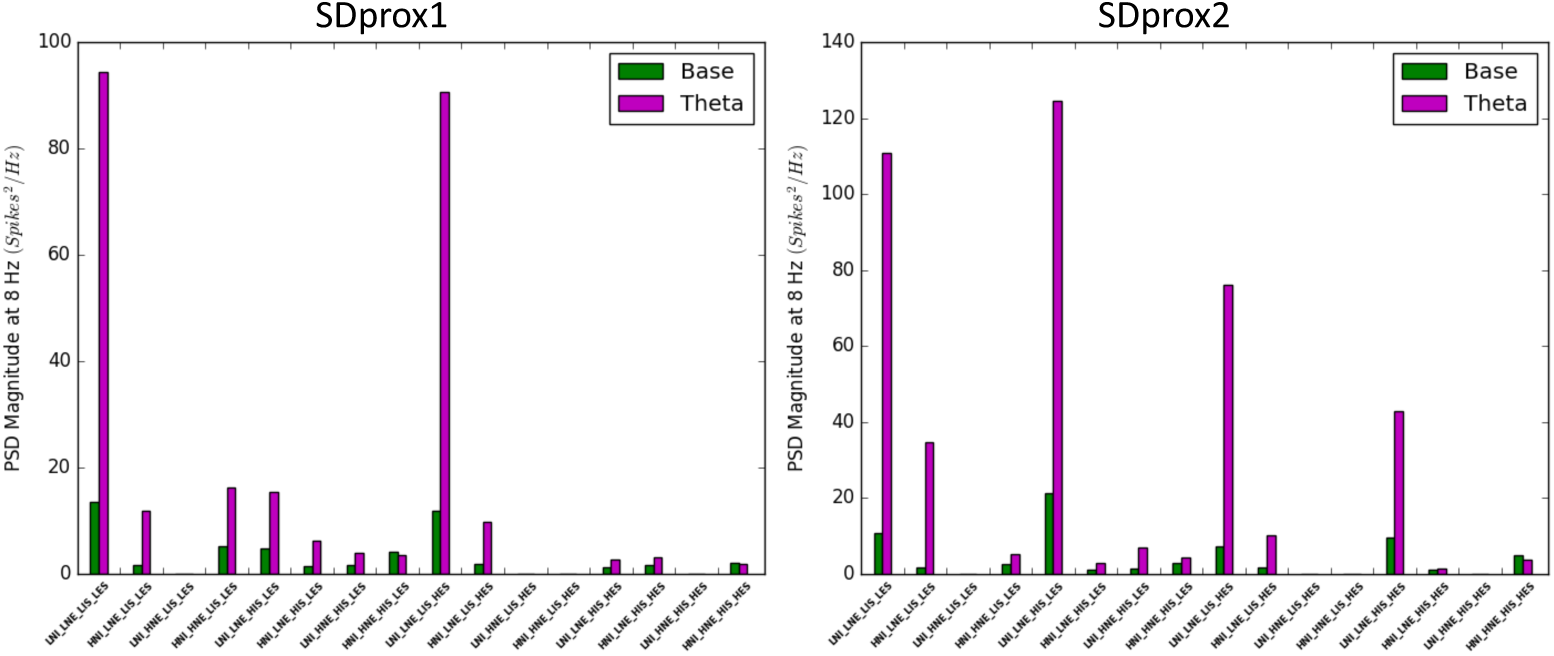
Power spectral densities (PSD) at 8 Hz computed from the spike trains of representative HC scenarios before (green) and after (magenta) the addition of theta-timed inputs.

**Figure 10–Figure supplement 3.**
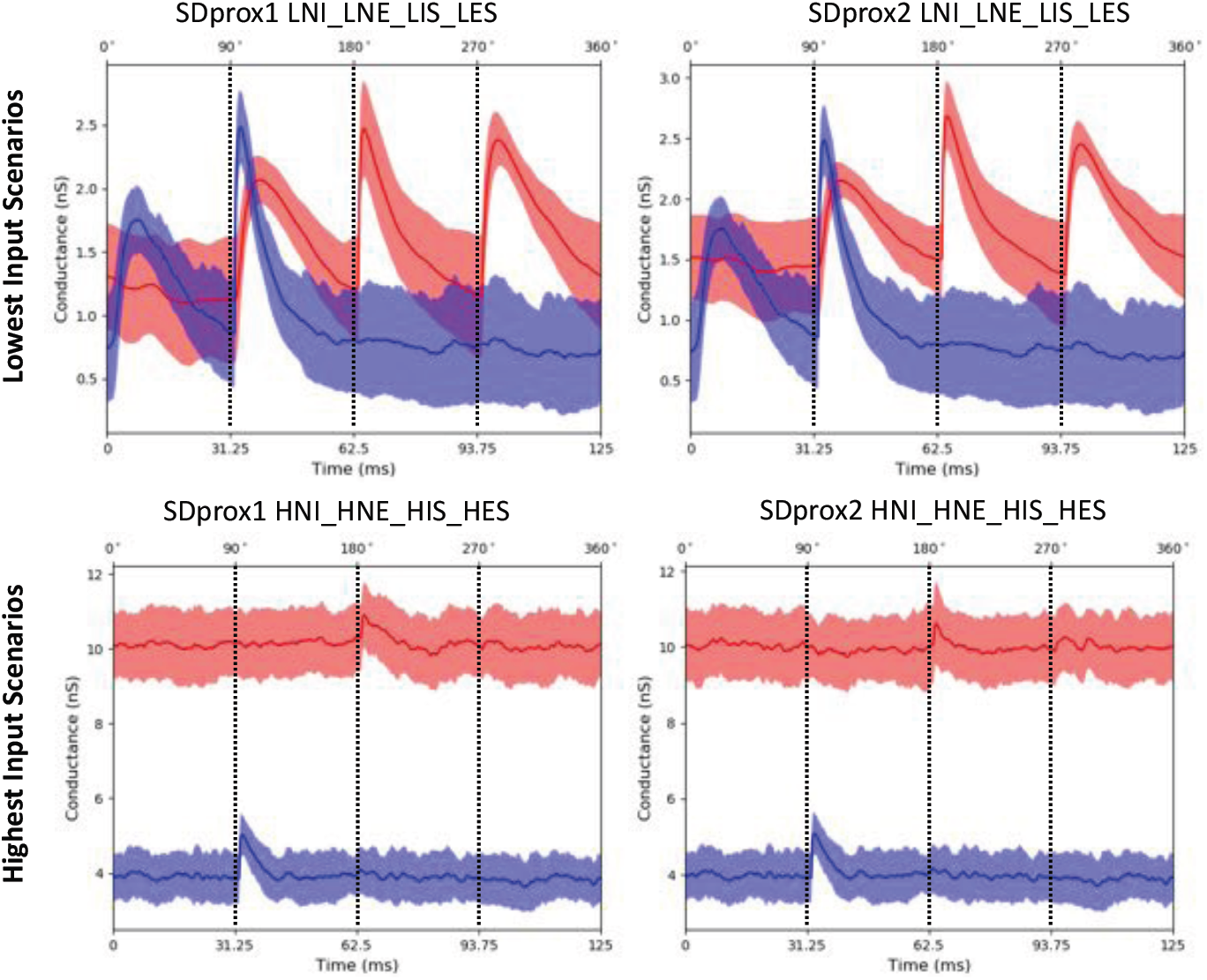
Example average inhibitory (red) and excitatory (blue) conductance traces during one theta cycle. We took the conductance traces (1000 ms to 10,000 ms), split them each into their 72 theta cycles (i.e. 9000 *ms*/125 *ms* = 72 *cycles*), and then computed the average 125 ms theta cycle traces for each representative scenario. Shaded areas show the amount of standard deviation above or below the mean. The top row of plots show results using the representative HC scenarios with the lowest amounts of inputs (LNI_LNE_LIS_LES). The bottom row of plots show the results using the representative HC scenarios with the highest amount of inputs (HNI_HNE_HIS_HES). In the LNI_LNE_LIS_LES scenario, clear peaks in conductance can be observed for when theta-timed inputs are occurring at 0°, 90°, 180°, and 270°.

**Figure 10–Figure supplement 4.**
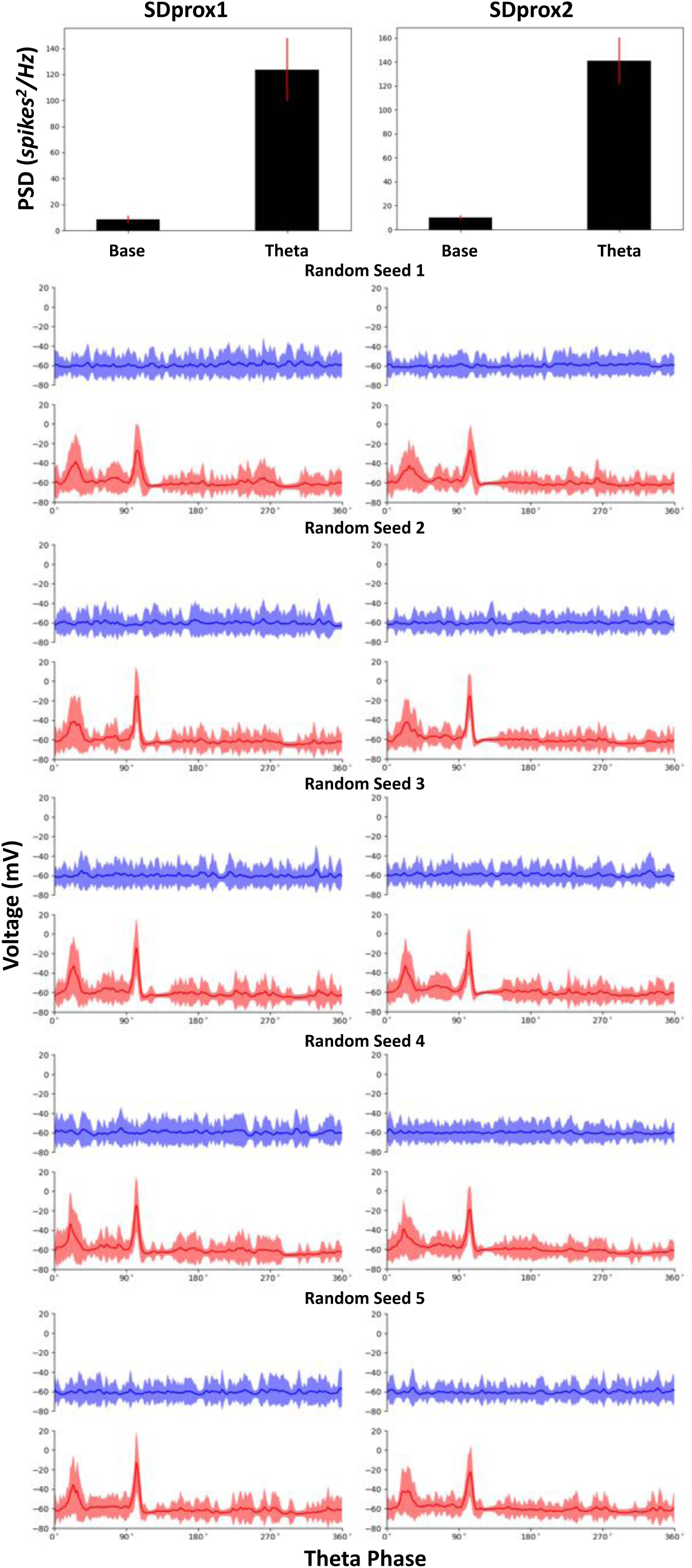
Consistency of theta recruitment in the LNI_LNE_LIS_LES representative scenario when re-randomizing synaptic locations and spike times. Top bar plots show the mean change in the PSD (red: standard deviation). Bottom plots show the mean *V*_*m*_ across all theta cycles in a trace (shaded areas: standard deviation; see ***Figure 10, Figure Supplement 3*** for more details). Blue traces show the baseline, and red traces show when theta-timed inputs are added.

## References

Acsády L, Arabadzisz D, Freund TF. Correlated morphological and neurochemical features identify different subsets of vasoactive intestinal polypeptide-immunoreactive interneurons in rat hippocampus. Neuroscience. 1996 Jul; 73(2):299–315. http://www.sciencedirect.com/science/article/pii/0306452295006109, doi: 10.1016/0306-4522(95)00610-9.

Acsády L, Görcs TJ, Freund TF. Different populations of vasoactive intestinal polypeptide-immunoreactive interneurons are specialized to control pyramidal cells or interneurons in the hippocampus. Neuroscience. 1996 Jul; 73(2):317–334. http://www.sciencedirect.com/science/article/pii/0306452295006095, doi: 10.1016/0306-4522(95)00609-5.

Adesnik H. Synaptic Mechanisms of Feature Coding in the Visual Cortex of Awake Mice. Neuron. 2017 Aug; 95(5):1147–1159.e4. http://www.cell.com/neuron/abstract/S0896-6273(17)30705-5, doi: 10.1016/j.neuron.2017.08.014.

Ahmed OJ, Kramer MA, Truccolo W, Naftulin JS, Potter NS, Eskandar EN, Cosgrove GR, Blum AS, Hochberg LR, Cash SS. Inhibitory single neuron control of seizures and epileptic traveling waves in humans. BMC Neuroscience. 2014 Jul; 15(1):F3. https://doi.org/10.1186/1471-2202-15-S1-F3, doi: 10.1186/1471-2202-15-S1-F3.

Atallah BV, Scanziani M. Instantaneous Modulation of Gamma Oscillation Frequency by Balancing Excitation with Inhibition. Neuron. 2009 May; 62(4):566–577. http://www.cell.com/neuron/abstract/S0896-6273(09)00351-1, doi: 10.1016/j.neuron.2009.04.027.

Bezaire MJ, Raikov I, Burk K, Vyas D, Soltesz I. Interneuronal mechanisms of hippocampal theta oscillations in a full-scale model of the rodent CA1 circuit. eLife. 2016 Dec; 5:e18566. https://elifesciences.org/articles/18566, doi: 10.7554/eLife.18566.

Bezaire MJ, Soltesz I. Quantitative Assessment of CA1 Local Circuits: Knowledge Base for Interneuron-Pyramidal Cell Connectivity. Hippocampus. 2013 Sep; 23(9):751–785. https://www.ncbi.nlm.nih.gov/pmc/articles/PMC3775914/, doi: 10.1002/hipo.22141.

Boyce R, Glasgow SD, Williams S, Adamantidis A. Causal evidence for the role of REM sleep theta rhythm in contextual memory consolidation. Science. 2016 May; 352(6287):812–816. http://science.sciencemag.org/content/352/6287/812, doi: 10.1126/science.aad5252.

Buzsáki G. Theta Oscillations in the Hippocampus. Neuron. 2002 Jan; 33(3):325–340. http://www.sciencedirect.com/science/article/pii/S089662730200586X, doi: 10.1016/S0896-6273(02)00586-X.

Canto CB, Witter MP. Cellular properties of principal neurons in the rat entorhinal cortex. I. The lateral entorhinal cortex. Hippocampus. 2012 Jun; 22(6):1256–1276. http://onlinelibrary.wiley.com/doi/10.1002/hipo.20997/abstract, doi: 10.1002/hipo.20997.

Canto CB, Witter MP. Cellular properties of principal neurons in the rat entorhinal cortex. II. The medial entorhinal cortex. Hippocampus. 2012 Jun; 22(6):1277–1299. http://onlinelibrary.wiley.com/doi/10.1002/hipo.20993/abstract, doi: 10.1002/hipo.20993.

Carnevale NT, Hines ML. The NEURON Book. 1 edition ed. Cambridge, UK; New York: Cambridge University Press; 2006.

Chamberland S, Salesse C, Topolnik D, Topolnik L. Synapse-Specific Inhibitory Control of Hippocampal Feedback Inhibitory Circuit. Frontiers in Cellular Neuroscience. 2010; 4. https://www.frontiersin.org/articles/10.3389/fncel.2010.00130/full, doi: 10.3389/fncel.2010.00130.

Chamberland S, Topolnik L. Inhibitory control of hippocampal inhibitory neurons. Frontiers in Neuroscience. 2012; 6. https://www.frontiersin.org/articles/10.3389/fnins.2012.00165/full, doi: 10.3389/fnins.2012.00165.

Chiu CQ, Martenson JS, Yamazaki M, Natsume R, Sakimura K, Tomita S, Tavalin SJ, Higley MJ. Input-Specific NMDAR-Dependent Potentiation of Dendritic GABAergic Inhibition. Neuron. 2018 Jan; 97(2):368–377.e3. http://www.cell.com/neuron/abstract/S0896-6273(17)31180-7, doi: 10.1016/j.neuron.2017.12.032.

Colgin LL. Rhythms of the hippocampal network. Nature Reviews Neuroscience. 2016 Apr; 17(4):239–249. https://www.nature.com/nrn/journal/v17/n4/full/nrn.2016.21.html, doi: 10.1038/nrn.2016.21.

David LS, Topolnik L. Target-specific alterations in the VIP inhibitory drive to hippocampal GABAergic cells after status epilepticus. Experimental Neurology. 2017 Jun; 292(Supplement C):102–112. http://www.sciencedirect.com/science/article/pii/S0014488617300687, doi: 10.1016/j.expneurol.2017.03.007.

Denève S, Machens CK. Effcient codes and balanced networks. Nature Neuroscience. 2016 Mar; 19(3):375–382. http://www.nature.com/neuro/journal/v19/n3/full/nn.4243.html?foxtrotcallback=true, doi: 10.1038/nn.4243.

Destexhe A, Paré D. Impact of Network Activity on the Integrative Properties of Neocortical Pyramidal Neurons In Vivo. Journal of Neurophysiology. 1999 Apr; 81(4):1531–1547. http://jn.physiology.org/content/81/4/1531.

Destexhe A, Rudolph M, Paré D. The high-conductance state of neocortical neurons in vivo. Nature Reviews Neuroscience. 2003 Sep; 4(9):739–751. http://www.nature.com/nrn/journal/v4/n9/full/nrn1198.html, doi: 10.1038/nrn1198.

Domnisoru C, Kinkhabwala AA, Tank DW. Membrane potential dynamics of grid cells. Nature. 2013 Mar; 495(7440):199–204. https://www.nature.com/nature/journal/v495/n7440/full/nature11973.html, doi: 10.1038/nature11973.

Ferguson KA, Huh CYL, Amilhon B, Manseau F, Williams S, Skinner FK. Network models provide insights into how oriens–lacunosum-moleculare and bistratified cell interactions influence the power of local hippocampal CA1 theta oscillations. Frontiers in Systems Neuroscience. 2015; 9. https://www.frontiersin.org/articles/10.3389/fnsys.2015.00110/full, doi: 10.3389/fnsys.2015.00110.

Frerking M, Schulte J, Wiebe SP, Stäubli U. Spike Timing in CA3 Pyramidal Cells During Behavior: Implications for Synaptic Transmission. Journal of Neurophysiology. 2005 Aug; 94(2):1528–1540. http://jn.physiology.org/content/94/2/1528, doi: 10.1152/jn.00108.2005.

Fu Y, Tucciarone JM, Espinosa JS, Sheng N, Darcy DP, Nicoll RA, Huang ZJ, Stryker MP. A Cortical Circuit for Gain Control by Behavioral State. Cell. 2014 Mar; 156(6):1139–1152. http://www.sciencedirect.com/science/article/pii/S0092867414001445, doi: 10.1016/j.cell.2014.01.050.

Goutagny R, Jackson J, Williams S. Self-generated theta oscillations in the hippocampus. Nature Neuroscience. 2009 Dec; 12(12):1491–1493. https://www.nature.com/neuro/journal/v12/n12/full/nn.2440.html, doi: 10.1038/nn.2440.

Guet-McCreight A, Camiré O, Topolnik L, Skinner FK. Using a Semi-Automated Strategy to Develop Multi-Compartment Models That Predict Biophysical Properties of Interneuron-Specific3 (IS3) Cells in Hippocampus. eNeuro. 2016 Aug; 3(4). doi: 10.1523/ENEURO.0087-16.2016.

Guet-McCreight A, Luo X, Francavilla R, Topolnik L, Skinner FK. Using computational modeling to estimate synaptic receptor densities along hippocampal CA1 interneuron specific 3 cell dendrites (Poster). F1000Research. 2017 Aug; 6. https://f1000research.com/posters/6-1552, doi: 10.7490/f1000research.1114780.1.

Gulyás AI, Hájos N, Freund TF. Interneurons Containing Calretinin Are Specialized to Control Other Interneurons in the Rat Hippocampus. Journal of Neuroscience. 1996 May; 16(10):3397–3411. http://www.jneurosci.org/content/16/10/3397.

Gulyás AI, Megias M, Emri Z, Freund TF. Total Number and Ratio of Excitatory and Inhibitory Synapses Converging onto Single Interneurons of Different Types in the CA1 Area of the Rat Hippocampus. Journal of Neuroscience. 1999 Nov; 19(22):10082–10097. http://www.jneurosci.org/content/19/22/10082.

Haider B, Duque A, Hasenstaub AR, McCormick DA. Neocortical Network Activity In Vivo Is Generated through a Dynamic Balance of Excitation and Inhibition. Journal of Neuroscience. 2006 Apr; 26(17):4535–4545. http://www.jneurosci.org/content/26/17/4535, doi: 10.1523/JNEUROSCI.5297-05.2006.

Harvey CD, Collman F, Dombeck DA, Tank DW. Intracellular dynamics of hippocampal place cells during virtual navigation. Nature. 2009 Oct; 461(7266):941–946. https://www.nature.com/nature/journal/v461/n7266/full/nature08499.html, doi: 10.1038/nature08499.

Higley MJ, Contreras D. Balanced Excitation and Inhibition Determine Spike Timing during Frequency Adaptation. Journal of Neuroscience. 2006 Jan; 26(2):448–457. http://www.jneurosci.org/content/26/2/448, doi: 10.1523/JNEUROSCI.3506-05.2006.

Ho ECY, Truccolo W. Interaction between synaptic inhibition and glial-potassium dynamics leads to diverse seizure transition modes in biophysical models of human focal seizures. Journal of Computational Neuroscience. 2016 Oct; 41(2):225–244. https://link.springer.com/article/10.1007/s10827-016-0615-7, doi: 10.1007/s10827-016-0615-7.

Huh CYL, Amilhon B, Ferguson KA, Manseau F, Torres-Platas SG, Peach JP, Scodras S, Mechawar N, Skinner FK, Williams S. Excitatory Inputs Determine Phase-Locking Strength and Spike-Timing of CA1 Stratum Oriens/Alveus Parvalbumin and Somatostatin Interneurons during Intrinsically Generated Hippocampal Theta Rhythm. Journal of Neuroscience. 2016 Jun; 36(25):6605–6622. http://www.jneurosci.org/content/36/25/6605, doi: 10.1523/JNEUROSCI.3951-13.2016.

Humphries MD. Motor Networks: The Goldilocks zone in neural circuits. eLife. 2016 Dec; 5:e22735. https://elifesciences.org/articles/22735, doi: 10.7554/eLife.22735.

Isaacson JS, Scanziani M. How Inhibition Shapes Cortical Activity. Neuron. 2011 Oct; 72(2):231–243. http://www.cell.com/neuron/abstract/S0896-6273(11)00879-8, doi: 10.1016/j.neuron.2011.09.027.

Jackson J, Ayzenshtat I, Karnani MM, Yuste R. VIP+ interneurons control neocortical activity across brain states. Journal of Neurophysiology. 2016 Jun; 115(6):3008–3017. http://jn.physiology.org/content/115/6/3008, doi: 10.1152/jn.01124.2015.

Kamigaki T, Dan Y. Delay activity of specific prefrontal interneuron subtypes modulates memory-guided behavior. Nature Neuroscience. 2017 Jun; 20(6):854–863. https://www.nature.com/neuro/journal/v20/n6/full/nn.4554.html, doi: 10.1038/nn.4554.

Kamondi A, Acsády L, Wang XJ, Buzsáki G. Theta oscillations in somata and dendrites of hippocampal pyramidal cells in vivo: Activity-dependent phase-precession of action potentials. Hippocampus. 1998 Jan; 8(3):244–261. http://onlinelibrary.wiley.com/doi/10.1002/(SICI)1098-1063(1998)8:3<244::AID-HIPO7>3.0.CO;2-J/abstract, doi: 10.1002/(SICI)1098-1063(1998)8:3<244::AID-HIPO7>3.0.CO;2-J.

Karnani MM, Jackson J, Ayzenshtat I, Sichani AH, Manoocheri K, Kim S, Yuste R. Opening Holes in the Blanket of Inhibition: Localized Lateral Disinhibition by VIP Interneurons. Journal of Neuroscience. 2016 Mar; 36(12):3471–3480. http://www.jneurosci.org/content/36/12/3471, doi: 10.1523/JNEUROSCI.3646-15.2016.

Karnani MM, Jackson J, Ayzenshtat I, Tucciarone J, Manoocheri K, Snider WG, Yuste R. Cooperative Subnetworks of Molecularly Similar Interneurons in Mouse Neocortex. Neuron. 2016 Apr; 90(1):86–100. http://www.cell.com/neuron/abstract/S0896-6273(16)00174-4, doi: 10.1016/j.neuron.2016.02.037.

Katona L, Lapray D, Viney TJ, Oulhaj A, Borhegyi Z, Micklem BR, Klausberger T, Somogyi P. Sleep and Movement Differentiates Actions of Two Types of Somatostatin-Expressing GABAergic Interneuron in Rat Hippocampus. Neuron. 2014 May; 82(4):872–886. http://www.cell.com/neuron/abstract/S0896-6273(14)00297-9, doi: 10.1016/j.neuron.2014.04.007.

Kepecs A, Fishell G. Interneuron cell types are fit to function. Nature. 2014 Jan; 505(7483):318. https://www.nature.com/articles/nature12983, doi: 10.1038/nature12983.

Klausberger T, Magill PJ, Márton LF, Roberts JDB, Cobden PM, Buzsáki G, Somogyi P. Brain-state-and cell-type-specific firing of hippocampal interneurons in vivo. Nature. 2003 Feb; 421(6925):844–848. https://www.nature.com/nature/journal/v421/n6925/full/nature01374.html, doi: 10.1038/nature01374.

Klausberger T, Somogyi P. Neuronal Diversity and Temporal Dynamics: The Unity of Hippocampal Circuit Operations. Science. 2008 Jul; 321(5885):53–57. http://www.sciencemag.org/content/321/5885/53, doi: 10.1126/science.1149381.

Kowalski J, Gan J, Jonas P, Pernía-Andrade A J. Intrinsic membrane properties determine hippocampal differential firing pattern in vivo in anesthetized rats. Hippocampus. 2016 May; 26(5):668–682. https://www.ncbi.nlm.nih.gov/pmc/articles/PMC5019144/, doi: 10.1002/hipo.22550.

Lapray D, Lasztoczi B, Lagler M, Viney TJ, Katona L, Valenti O, Hartwich K, Borhegyi Z, Somogyi P, Klausberger T. Behavior-dependent specialization of identified hippocampal interneurons. Nature Neuroscience. 2012 Sep; 15(9):1265–1271. https://www.nature.com/neuro/journal/v15/n9/full/nn.3176.html, doi: 10.1038/nn.3176.

Lee S, Kruglikov I, Huang ZJ, Fishell G, Rudy B. A disinhibitory circuit mediates motor integration in the somatosensory cortex. Nature Neuroscience. 2013 Nov; 16(11):1662–1670. https://www.nature.com/neuro/journal/v16/n11/full/nn.3544.html, doi: 10.1038/nn.3544.

Leutgeb S, Leutgeb JK, Treves A, Moser MB, Moser EI. Distinct Ensemble Codes in Hippocampal Areas CA3 and CA1. Science. 2004 Aug; 305(5688):1295–1298. http://science.sciencemag.org/content/305/5688/1295, doi: 10.1126/science.1100265.

Liu G. Local structural balance and functional interaction of excitatory and inhibitory synapses in hippocampal dendrites. Nature Neuroscience. 2004 Apr; 7(4):373–379. https://www.nature.com/articles/nn1206, doi: 10.1038/nn1206.

Luczak A, McNaughton BL, Harris KD. Packet-based communication in the cortex. Nature Reviews Neuroscience. 2015 Dec; 16(12):745–755. https://www.nature.com/articles/nrn4026, doi: 10.1038/nrn4026.

Magee J, Hoffman D, Colbert C, Johnston D. Electrical and Calcium Signaling in Dendrites of Hippocampal Pyramidal Neurons. Annual Review of Physiology. 1998; 60(1):327–346. https://doi.org/10.1146/annurev.physiol.60.1.327, doi: 10.1146/annurev.physiol.60.1.327.

Markram H, Toledo-Rodriguez M, Wang Y, Gupta A, Silberberg G, Wu C. Interneurons of the neocortical inhibitory system. Nature Reviews Neuroscience. 2004 Oct; 5(10):793. https://www.nature.com/articles/nrn1519, doi: 10.1038/nrn1519.

Marín O. Interneuron dysfunction in psychiatric disorders. Nature Reviews Neuroscience. 2012 Feb; 13(2):107–n120.https://www.nature.com/articles/nrn3155, doi: 10.1038/nrn3155.

McBain CJ, Fisahn A. Interneurons unbound. Nature Reviews Neuroscience. 2001 Jan; 2(1):11. https://www.nature.com/articles/35049047, doi: 10.1038/35049047.

Megias M, Emri Z, Freund TF, Gulyás AI. Total number and distribution of inhibitory and excitatory synapses on hippocampal CA1 pyramidal cells. Neuroscience. 2001 Feb; 102(3):527–540. https://www.sciencedirect.com/science/article/pii/S0306452200004966, doi: 10.1016/S0306-4522(00)00496-6.

Mizuseki K, Sirota A, Pastalkova E, Buzsáki G. Theta Oscillations Provide Temporal Windows for Local Circuit Computation in the Entorhinal-Hippocampal Loop. Neuron. 2009 Oct; 64(2):267–280. http://www.cell.com/neuron/abstract/S0896-6273(09)00673-4, doi: 10.1016/j.neuron.2009.08.037.

Monier C, Fournier J, Frégnac Y. In vitro and in vivo measures of evoked excitatory and inhibitory conductance dynamics in sensory cortices. Journal of Neuroscience Methods. 2008 Apr; 169(2):323–365. http://www.sciencedirect.com/science/article/pii/S0165027007005547, doi: 10.1016/j.jneumeth.2007.11.008.

Morin F, Haufler D, Skinner FK, Lacaille JC. Characterization of Voltage-Gated K+ Currents Contributing to Subthreshold Membrane Potential Oscillations in Hippocampal CA1 Interneurons. Journal of Neurophysiology. 2010 Apr; 103(6):3472–3489. http://www.physiology.org/doi/abs/10.1152/jn.00848.2009, doi: 10.1152/jn.00848.2009.

O’Donnell C, Gonçalves JT, Portera-Cailliau C, Sejnowski TJ. Beyond excitation/inhibition imbalance in multidimensional models of neural circuit changes in brain disorders. eLife. 2017 Oct; 6:e26724. https://elifesciences.org/articles/26724, doi: 10.7554/eLife.26724.

Ouyang YB, Ning S, Adler JR, Maciver B, Knox SJ, Giffard R. Alteration of Interneuron Immunoreactivity and Autophagic Activity in Rat Hippocampus after Single High-Dose Whole-Brain Irradiation. Cureus. 2017 Jun; 9(6). https://www.ncbi.nlm.nih.gov/pmc/articles/PMC5576964/, doi: 10.7759/cureus.1414.

Pelkey KA, Chittajallu R, Craig MT, Tricoire L, Wester JC, McBain CJ. Hippocampal GABAergic Inhibitory Interneurons. Physiological Reviews. 2017 Oct; 97(4):1619–1747. http://physrev.physiology.org/content/97/4/1619, doi: 10.1152/physrev.00007.2017.

Petersen PC, Berg RW. Lognormal firing rate distribution reveals prominent fluctuation–driven regime in spinal motor networks. eLife. 2016 Oct; 5:e18805. https://elifesciences.org/articles/18805, doi: 10.7554/eLife.18805.

Pi HJ, Hangya B, Kvitsiani D, Sanders JI, Huang ZJ, Kepecs A. Cortical interneurons that specialize in disinhibitory control. Nature. 2013 Nov; 503(7477):521–524. https://www.nature.com/nature/journal/v503/n7477/full/nature12676.html, doi: 10.1038/nature12676.

Piwkowska Z, Pospischil M, Brette R, Sliwa J, Rudolph-Lilith M, Bal T, Destexhe A. Characterizing synaptic conductance fluctuations in cortical neurons and their influence on spike generation. Journal of Neuroscience Methods. 2008 Apr; 169(2):302–322. http://www.sciencedirect.com/science/article/pii/S0165027007005572, doi: 10.1016/j.jneumeth.2007.11.010.

Ramaswamy S, Courcol JD, Abdellah M, Adaszewski SR, Antille N, Arsever S, Atenekeng G, Bilgili A, Brukau Y, Chalimourda A, Chindemi G, Delalondre F, Dumusc R, Eilemann S, Gevaert ME, Gleeson P, Graham JW, Hernando JB, Kanari L, Katkov Y, et al. The neocortical microcircuit collaboration portal: a resource for rat somatosensory cortex. Frontiers in Neural Circuits. 2015; p. 44. http://journal.frontiersin.org/article/10.3389/fncir.2015.00044/full, doi: 10.3389/fncir.2015.00044.

Reimann MW, King JG, Muller EB, Ramaswamy S, Markram H. An algorithm to predict the connectome of neural microcircuits. Frontiers in Computational Neuroscience. 2015; p. 120. http://journal.frontiersin.org/article/10.3389/fncom.2015.00120/abstract, doi: 10.3389/fncom.2015.00120.

Rosenbaum R, Smith MA, Kohn A, Rubin JE, Doiron B. The spatial structure of correlated neuronal variability. Nature Neuroscience. 2017 Jan; 20(1):107–114. https://www.nature.com/articles/nn.4433, doi: 10.1038/nn.4433.

Salz DM, Tiganj Z, Khasnabish S, Kohley A, Sheehan D, Howard MW, Eichenbaum H. Time Cells in Hippocampal Area CA3. The Journal of Neuroscience: The Official Journal of the Society for Neuroscience. 2016 Jul; 36(28):7476–7484. doi: 10.1523/JNEUROSCI.0087-16.2016.

Schreiber S, Samengo I, Herz AVM. Two Distinct Mechanisms Shape the Reliability of Neural Responses. Journal of Neurophysiology. 2009 May; 101(5):2239–2251. http://jn.physiology.org/content/101/5/2239, doi: 10.1152/jn.90711.2008.

Sekulic V, Skinner FK. Computational models of O-LM cells are recruited by low or high theta frequency inputs depending on h-channel distributions. eLife. 2017 Mar; 6:e22962. https://elifesciences.org/articles/22962, doi: 10.7554/eLife.22962.

Shadlen MN, Newsome WT. Noise, neural codes and cortical organization. Current Opinion in Neurobiology. 1994 Aug; 4(4):569–579. http://www.sciencedirect.com/science/article/pii/0959438894900590, doi: 10.1016/0959-4388(94)90059-0.

Sivagnanam S, Majumdar A, Yoshimoto K, Astakhov V B A, Martone M, Carnevale NT. Introducing The Neuroscience Gateway, vol. 993 of CEUR Workshop Proceedings of CEUR Workshop Proceedings; 2013. http://ceur-ws.org/Vol-993/paper10.pdf.

Softky WR, Koch C. The highly irregular firing of cortical cells is inconsistent with temporal integration of random EPSPs. The Journal of Neuroscience. 1993 Jan; 13(1):334–350. http://www.jneurosci.org/content/13/1/334.

Stevens CF, Zador AM. Input synchrony and the irregular firing of cortical neurons. Nature Neuroscience. 1998 Jul; 1(3):210–217. https://www.nature.com/neuro/journal/v1/n3/full/nn0798_210.html, doi: 10.1038/659.

Stringer C, Pachitariu M, Steinmetz NA, Okun M, Bartho P, Harris KD, Sahani M, Lesica NA. Inhibitory control of correlated intrinsic variability in cortical networks. eLife. 2016 Dec; 5:e19695. https://elifesciences.org/articles/19695, doi: 10.7554/eLife.19695.

Taylor AL, Hickey TJ, Prinz AA, Marder E. Structure and Visualization of High-Dimensional Conductance Spaces. Journal of Neurophysiology. 2006 Aug; 96(2):891–905. http://jn.physiology.org/content/96/2/891, doi: 10.1152/jn.00367.2006.

Trevelyan A J, Sussillo D, Watson BO, Yuste R. Modular Propagation of Epileptiform Activity: Evidence for an Inhibitory Veto in Neocortex. Journal of Neuroscience. 2006 Nov; 26(48):12447–12455. http://www.jneurosci.org/content/26/48/12447, doi: 10.1523/JNEUROSCI.2787-06.2006.

Treviño M. Inhibition Controls Asynchronous States of Neuronal Networks. Frontiers in Synaptic Neuroscience. 2016; 8. https://www.frontiersin.org/articles/10.3389/fnsyn.2016.00011/full, doi: 10.3389/fnsyn.2016.00011.

Turrigiano G. Too Many Cooks? Intrinsic and Synaptic Homeostatic Mechanisms in Cortical Circuit Refinement. Annual Review of Neuroscience. 2011; 34(1):89–103. https://doi.org/10.1146/annurev-neuro-060909-153238, doi: 10.1146/annurev-neuro-060909-153238.

Tyan L, Chamberland S, Magnin E, Camiré O, Francavilla R, David LS, Deisseroth K, Topolnik L. Dendritic Inhibition Provided by Interneuron-Specific Cells Controls the Firing Rate and Timing of the Hippocampal Feedback Inhibitory Circuitry. Journal of Neuroscience. 2014 Mar; 34(13):4534–4547. http://www.jneurosci.org/content/34/13/4534, doi: 10.1523/JNEUROSCI.3813-13.2014.

Varga C, Golshani P, Soltesz I. Frequency-invariant temporal ordering of interneuronal discharges during hippocampal oscillations in awake mice. Proceedings of the National Academy of Sciences. 2012 Oct; 109(40):E2726–E2734. http://www.pnas.org/content/109/40/E2726, doi: 10.1073/pnas.1210929109.

Varga C, Oijala M, Lish J, Szabo GG, Bezaire M, Marchionni I, Golshani P, Soltesz I. Functional fission of parvalbumin interneuron classes during fast network events. eLife. 2014 Nov; 3:e04006. https://elifesciences.org/articles/04006, doi: 10.7554/eLife.04006.

Vida I. Morphology of Hippocampal Neurons. In: Hippocampal Microcircuits Springer Series in Computational Neuroscience, Springer, New York, NY; 2010. p. 27–67. https://link.springer.com/chapter/10.1007/978-1-4419-0996-1_2, doi: 10.1007/978-1-4419-0996-1_2.

Villette V, Levesque M, Miled A, Gosselin B, Topolnik L. Simple platform for chronic imaging of hippocampal activity during spontaneous behaviour in an awake mouse. Scientific Reports. 2017 Feb; 7:srep43388. https://www.nature.com/articles/srep43388, doi: 10.1038/srep43388.

Vinet J, Sík A. Expression pattern of voltage-dependent calcium channel subunits in hippocampal inhibitory neurons in mice. Neuroscience. 2006 Nov; 143(1):189–212. http://www.sciencedirect.com/science/article/pii/S0306452206009997, doi: 10.1016/j.neuroscience.2006.07.019.

Vreeswijk Cv, Sompolinsky H. Chaos in Neuronal Networks with Balanced Excitatory and Inhibitory Activity. Science. 1996 Dec; 274(5293):1724–1726. http://science.sciencemag.org/content/274/5293/1724, doi: 10.1126/science.274.5293.1724.

Witter MP. Connectivity of the Hippocampus. In: Hippocampal Microcircuits Springer Series in Computational Neuroscience, Springer, New York, NY; 2010. p. 5–26. https://link.springer.com/chapter/10.1007/978-1-4419-0996-1_1, doi: 10.1007/978-1-4419-0996-1_1.

Xue M, Atallah BV, Scanziani M. Equalizing excitation-inhibition ratios across visual cortical neurons. Nature. 2014 Jul; 511(7511):596–600. https://www.nature.com/nature/journal/v511/n7511/full/nature13321.html, doi: 10.1038/nature13321.

Zeng H, Sanes JR. Neuronal cell-type classification: challenges, opportunities and the path forward. Nature Reviews Neuroscience. 2017 Sep; 18(9):530. https://www.nature.com/articles/nrn.2017.85, doi: 10.1038/nrn.2017.85.

Zhang S, Xu M, Kamigaki T, Do JPH, Chang WC, Jenvay S, Miyamichi K, Luo L, Dan Y. Long-range and local circuits for top-down modulation of visual cortex processing. Science. 2014 Aug; 345(6197):660–665. http://science.sciencemag.org/content/345/6197/660, doi: 10.1126/science.1254126.

Zhou S, Yu Y. Synaptic E-I Balance Underlies Effcient Neural Coding. Frontiers in Neuroscience. 2018; 12. https://www.frontiersin.org/articles/10.3389/fnins.2018.00046/full, doi: 10.3389/fnins.2018.00046.

